# Functional Characterization of 123 Genes from Mycobacteriophage LeBron Uncovers Cytotoxic Gene Families in Cluster L1

**DOI:** 10.64898/2025.12.17.694915

**Authors:** Earick J Cagang, Brian Nguyen, Elizabeth Paul, Elva Garcia, Alyssa Lee, Sabrina Benitez, Rita Dementyev, Ethan Dewri, Alexandra Falvo, Christian Figueroa, Dulce Guevara, Katie Jang, Michael Kelly, Caleb Kim, Soojeong Moon, Kristen Ngo, Ester Peiro, Daphne Prakash, Noboyuki Yano, Jiacheng Zhang, Danielle M. Heller, Arturo Diaz

**Affiliations:** Department of Biology, La Sierra University, Riverside, California, 92503, USA; Department of Science Education, Howard Hughes Medical Institute, Chevy Chase, Maryland 20815, USA

**Keywords:** mycobacteriophage, *Mycobacterium smegmatis*, cytotoxicity, genome-wide screen

## Abstract

Bacteriophages represent a vast reservoir of genetic diversity however, functional annotation remains a major challenge, as most predicted gene products lack detectable similarity to characterized gene families. Experimental approaches such as systematic overexpression screens provide an avenue for identifying phage genes that influence bacterial physiology. Here, we report an overexpression screen of all 123 predicted protein-coding genes from Cluster L1 mycobacteriophage LeBron, the first representative of this cluster to undergo genome-wide functional analysis. Expression assays in *Mycobacterium smegmatis* revealed that 39 genes (32%) impaired host growth, with nineteen of these toxic genes (49%) assigned no known function. The proportion of cytotoxic genes observed in LeBron is comparable to findings from Clusters K and F, despite minimal sequence conservation across clusters. Interestingly, a subset of LeBron’s toxic genes appear to be functionally analogous to previously identified toxic genes in other clusters, suggesting conserved biological activities carried out by non-homologous proteins. Additionally, this analysis uncovered several novel gene families that elicit strong cytotoxic effects, expanding the known catalog of phage-derived bacterial growth inhibitors. These results provide new insights into phage gene functions and demonstrate the value of genome-wide expression screening for uncovering conserved and cluster-specific interactions between bacteriophages and their hosts.

## Introduction

Bacteriophages and bacteria have coexisted in a constant evolutionary arms race, giving rise to extraordinary genetic diversity and a wide range of host–phage interactions (Hampton et al. 2020). Phage genomes encode both a conserved core of structural and replication functions and a variable array of accessory genes that can modulate host physiology, promote competition with other phages, or circumvent bacterial defense systems (Dedrick et al. 2013; Dedrick et al. 2017; Dion et al. 2020; Hatfull 2020; Yuan et al. 2023). Comparative analyses of sequenced phage genomes have revealed highly mosaic architectures composed of thousands of distinct gene families, or phamilies, with the vast majority lacking recognizable sequence motifs or functional annotation (Cresawn et al. 2011; Pope et al. 2011; Pope et al. 2015; Pope et al. 2017; Jacobs-Sera et al. 2020; Gauthier and Hatfull 2023; Hyde et al. 2023). The study of these poorly characterized genes has yielded insights into fundamental processes, from restriction–modification and CRISPR–Cas defense systems to novel mechanisms of inter-bacterial competition (Loenen et al. 2014; Adli 2018; Yuan et al. 2023). Thus, phage genomes represent a reservoir of untapped biological functions with implications for microbial evolution, therapeutic development, and biotechnology.

One productive experimental approach for probing these unknown functions is systematic overexpression screening, in which individual phage genes are assayed for phenotypic effects in the bacterial host. These assays often reveal growth-inhibitory genes, pointing to proteins that disrupt or interact with essential host processes (Liu et al. 2004; Molshanski-Mor et al. 2014; Ko and Hatfull 2018; Ko and Hatfull 2020; Freeman et al. 2024). Recent genetic overexpression screens carried out under the HHMI-supported SEA-GENES project (Science Education Alliance Gene-function Exploration by a Network of Emerging Scientists) revealed that roughly one-quarter to one-third of the genes encoded in a given mycobacteriophage genome inhibit *Mycobacterium smegmatis* growth to varying degrees (Heller et al. 2022; Amaya et al. 2023; Freeman et al. 2024; Pollenz et al. 2024; Tafoya et al. 2025). Actinobacteriophage genomes are assigned to clusters based on shared genetic content (Gauthier and Hatfull 2023), and these groupings provide a framework for comparing inhibitory gene functions across related viruses. In Cluster K, for example, genome-wide analyses of phages Hammy (K6), Waterfoul (K5), and Amelie (K1) identified toxicities in about one-third of their genes, with many inhibitors conserved across related subclusters but others unique to each genome (Heller et al. 2022; Amaya et al. 2023; Tafoya et al. 2025). Similarly, analyses of Cluster F1 phages Girr (Pollenz et al. 2024) and NormanBulbieJr (Wise et al. 2025) revealed 28% of tested genes as toxic to *M. smegmatis*, underscoring a comparable range of inhibitory potential across genetically distinct clusters despite minimal sequence conservation. Together, these studies highlight that cytotoxic genes are not confined to specific clusters but rather broadly distributed across phage lineages, suggesting that bacteriophages have independently evolved diverse genetic strategies for engaging and disrupting host cellular processes.

Systematic functional characterization of additional phage clusters is essential to capture new categories of bacterial growth inhibitors and to assess the extent of shared versus unique genetic strategies. Here we describe a genome-wide functional screen of *Mycobacterium smegmatis* genes from bacteriophage LeBron, a Cluster L1 mycobacteriophage. At the level of gene content similarity (GCS), which reflects the percentage of gene “phamilies” (groups of related genes) that are found in both genomes, LeBron shares only 9.4–9.5% similarity with Cluster K phages and just 3.6% with Cluster F1 phages, underscoring its distant relationship to clusters that have already been subjected to functional screening. Of its 123 predicted protein-coding genes, 39 were found to impair bacterial growth when overexpressed, including a set of novel gene phamilies with no functional homologs in other clusters. Comparative analysis further revealed instances where LeBron’s toxic genes resemble known bacterial inhibitors from unrelated clusters, underscoring the occurrence of functional convergence across diverse gene sequences. Together, these results broaden our understanding of the genetic strategies employed by Cluster L1 phages and emphasize the value of genome-wide expression assays in uncovering both conserved and cluster-specific mechanisms of phage-host interaction.

## Materials and Methods

### Bacterial and Phage Culture Conditions

*Mycobacterium smegmatis* mc^2^155 was cultured at 37°C in Middlebrook 7H9 broth (Millipore), supplemented with 10% albumin-dextrose (AD; 2% w/v dextrose, 145 mM NaCl, 5% w/v albumin fraction V), 0.05% Tween-80, and 10 µg/mL cycloheximide or grown on Middlebrook 7H10 agar (Millipore) supplemented with 10% AD and 10 µg/mL cycloheximide. For plasmid transformation, 40–200 ng of DNA was introduced into electrocompetent *M. smegmatis* via electroporation, followed by 90 minutes of recovery in liquid 7H9 medium at 37°C with shaking. Transformants were selected on 7H10 agar plates containing 10 µg/mL kanamycin (GoldBio) and incubated at 37°C for 5 days prior to phenotypic screening. For propagation of mycobacteriophage LeBron, host cultures were grown in 7H9 broth lacking Tween-80 but supplemented with 1 mM CaCl_2_ to promote phage adsorption, using standard top agar overlay methods.

### Construction of the pExTra-Amelie Expression Library

Each LeBron gene was individually inserted into the pExTra shuttle vector (Heller et al. 2022), which drives expression from an anhydrotetracycline (aTc)-inducible pTet promoter and includes an in-frame downstream transcriptional fusion to *mCherry*. Target genes were PCR amplified from high-titer LeBron lysates using Q5 Hot Start DNA Polymerase (New England Biolabs) and gene-specific primers (Integrated DNA Technologies). Forward primers annealed to the 5′ region of each coding sequence and standardized the start codon to ATG, while reverse primers annealed to the 3′ end of the gene and introduced a uniform TGA stop. All primers were designed with homology arms facilitating Gibson assembly into a linearized backbone. Primer sequences and homology arms are provided in Supplementary Table 1. The pExTra01 vector backbone was linearized by double-restriction enzyme digest with NdeI and SalI (New England Biolabs). Gene fragments and linearized pExTra were joined using NEB HiFi DNA assembly, and recombinant plasmids were recovered following transformation into *E. coli* NEB5α F’IQ cells, with selection on LB agar containing 50 µg/mL kanamycin. Plasmid inserts were verified by Sanger sequencing (Azenta) using universal primers (pExTra_seqF and pExTra_universalR). Longer gene constructs were sequenced using additional internal primers (see Supplementary Table 1). All cloned genes matched the published LeBron genome sequence (GenBank accession HM152763).

### Cytotoxicity Assays and Phenotypic Classification

To evaluate gene-specific cytotoxicity, three independent colonies of *M. smegmatis* harboring each pExTra-LeBron construct were resuspended in 7H9 broth, serially diluted, and spotted onto 7H10 agar supplemented with 10 µg/mL kanamycin and either 0, 10, or 100 ng/mL anhydrotetracycline (aTc; IBA LifeSciences). Spot assays were performed in triplicate and included two controls: pExTra02 (positive control, encoding the toxic Fruitloop gene *52*) and pExTra03 (negative control, encoding a non-toxic I70S mutant of the same gene) (Ko and Hatfull 2018; Heller et al. 2022). Plates were incubated at 37°C, and colony growth was monitored over a 5-day period.

Toxic phenotypes were scored based on both aTc-dependent growth inhibition and comparison to control strains. A four-point classification system was used: 0 = No visible growth inhibition relative to controls; 1 = Reduced colony size, indicating partial toxicity; 2 = 1–3 log reduction in colony-forming units; 3 = Severe toxicity with >3 log reduction or complete growth arrest. For constructs scoring 0, expression of *mCherry* under aTc conditions confirmed successful gene induction. Final cytotoxicity calls were based on results from 2–4 independent experiments per gene. Effects were only scored as toxic if consistent growth inhibition was observed and exceeded any background growth defects seen in negative controls. Where inter-experimental variability occurred for mildly toxic genes, the lowest observed toxicity was reported.

### Genome Annotation and Functional Prediction

The LeBron genome was analyzed using Phamerator (Cresawn et al. 2011) to generate genome maps. Initial annotations were based on the GenBank entry (accession KX808132) and then refined using the Phage Evidence Collection and Annotation Network (PECAAN). Functional evidence was drawn from multiple resources: BLASTp (Altschul et al. 1990; Russell and Hatfull 2017), HHPRED (Gabler et al. 2020), Conserved Domain Database (Marchler-Bauer et al. 2015), Pfam, SCOPe, DeepTMHMM (Hallgren et al. 2022), and TOPCONS (Tsirigos et al. 2015). Stringent thresholds were applied (HHPRED probability >90%, alignment coverage >50%, and e-value < 1×10^−5^) to assign putative protein functions. Proteins not meeting these standards were classified as hypothetical with no known function (NKF). Common domain features, including domains of unknown function (DUFs), were annotated through CDD searches. Common structural or regulatory motifs, such as domains of unknown function (DUFs) and helix-turn-helix (HTH) motifs, were identified through domain databases and verified using the NPS HTH predictor (https://npsa-prabi.ibcp.fr/).

## Results and Discussion

### Study Overview

To evaluate the impact of LeBron gene expression on *M. smegmatis* mc^2^155 cell growth, all 123 predicted protein-coding sequences were cloned into the pExTra plasmid, where transcription is driven by the inducible pTet promoter and monitored with an mCherry flourescent reporter (Fig. 1a). 122 genes were included in this analysis as *M. smegmatis* cells transformed with a pExtra plasmid encoding gene 46, a phosphoesterase, could not be recovered despite multiple attempts. Moreover, this analysis also excluded the nine tRNAs encoded in the LeBron genome. Each construct was sequence-verified and subsequently introduced into *M. smegmatis* for analysis in a semiquantitative, plate-based cytotoxicity assay. For every gene, three independent transformants were suspended and used to generate 10-fold dilution series, which were then spotted onto media containing either no inducer or increasing concentrations of aTc. Control strains expressing the wild-type cytotoxic gene Fruitloop *52* or its nontoxic mutant allele were included in each assay (Ko and Hatfull 2018).

**Figure 1.**
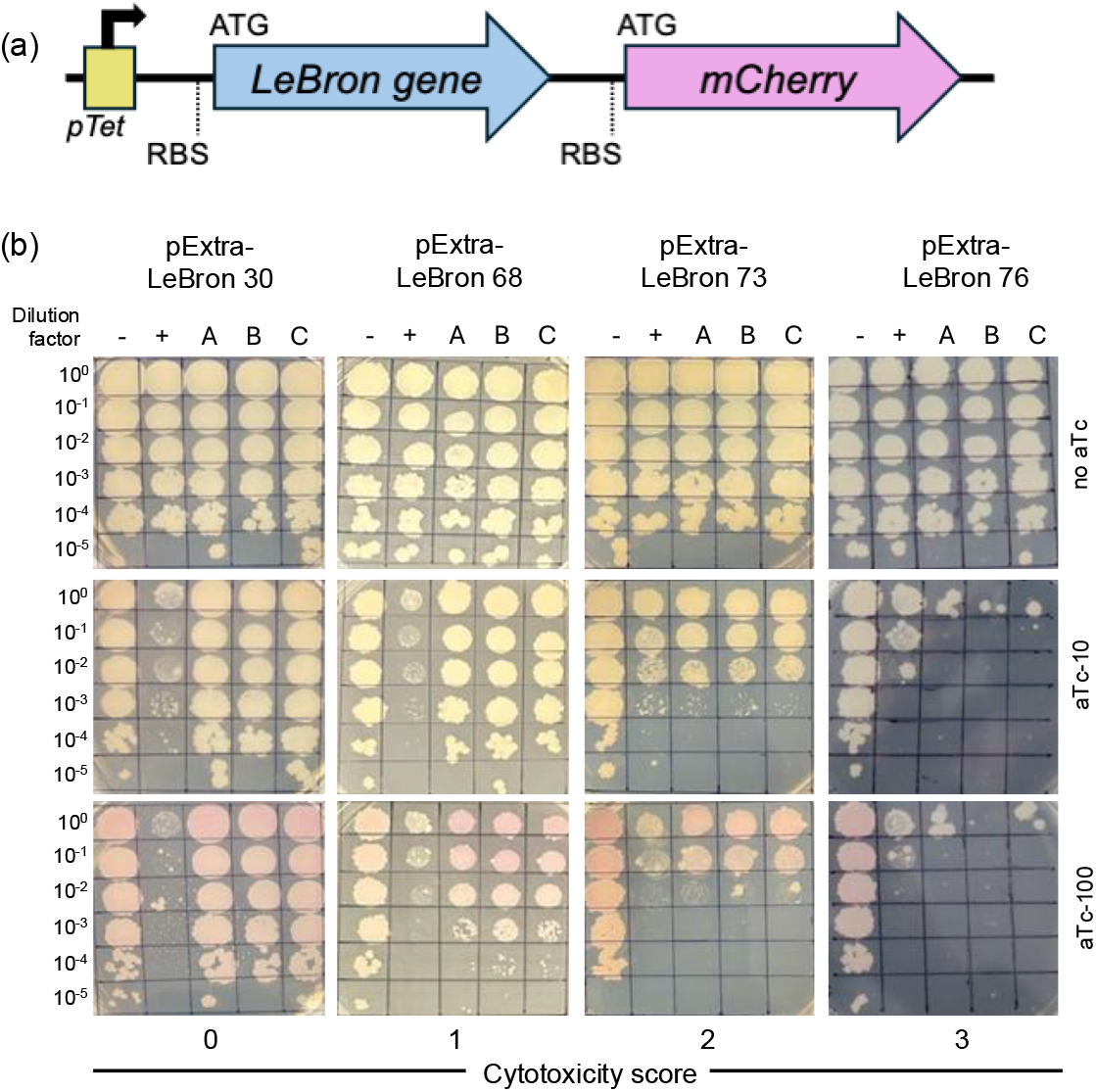
Cytotoxicity assay design and representative results for phage LeBron genes. a) Recombinant pExTra plasmids constructed in this study encode LeBron gene sequences downstream of the pTet promoter and upstream of *mCherry*. The two genes are arranged in an aTc-inducible operon, each with its own start and stop codons and ribosome binding site (RBS), ensuring independent translation of both protein products. b) Representative cytotoxicity assays illustrate the range of observed growth defects. Colonies of *M. smegmatis* mc2155 transformed with the indicated pExTra plasmids were resuspended, serially diluted, and spotted on 7H10 Kan media supplemented with 0, 10, or 100 ng/ml aTc. Triplicate colonies (A, B, C) were tested for each gene alongside a positive control strain (+, pExTra02 expressing wild-type Fruitloop gp52) and a negative control strain ( −, pExTra03 expressing Fruitloop gp52 I70S). Examples of each cytotoxicity score are shown: score 0 (LeBron gp30), score 1 (LeBron gp68), score 2 (LeBron gp73), and score 3 (LeBron gp47).

Cytotoxic effects were classified into four categories based on the extent of growth inhibition: no observable effect on viability (score 0), reduced colony size relative to controls (score 1), moderate toxicity with a 1–3 log reduction in viable cell count (score 2), and severe cytotoxicity, with complete or near-complete (>3 log) inhibition of growth (score 3). Representative examples are shown in Figure 1b: overexpression of LeBron 68 resulted in mild colony size reduction (score 1), LeBron 73 exhibited moderate cytotoxicity (score 2), and LeBron 76 caused severe growth inhibition across all replicates (score 3). Interestingly, nine LeBron genes (*1, 23, 26, 32, 34, 36, 117, 123*, and *126*) produced visibly pink colonies even in the absence of inducer, suggesting that these sequences may harbor internal promoter elements (Supplementary Fig. 1). For all other non-toxic genes, the appearance of pink colonies in the presence of aTc confirmed successful induction of the *pTet* operon and production of the linked mCherry reporter. The complete dataset and cytotoxicity scores for all 122 LeBron genes evaluated in this study are presented in Supplementary Figure 1.

### Overexpression screen reveals 39 cytotoxic LeBron genes

Expression of 84 out of the 122 LeBron protein-coding genes did not appreciably impact *M. smegmatis* growth in our assay (Fig. 2; Supplementary Fig. 1). For most of these nontoxic genes (70 out of 84), induction with the highest concentration of aTc resulted in visible pink coloration of colonies, confirming *pTet*-driven expression of the reporter. As in previous studies, protein accumulation was not directly measured, so it remains possible that some products classified as nontoxic were either poorly or not expressed in our system.

**Figure 2.**
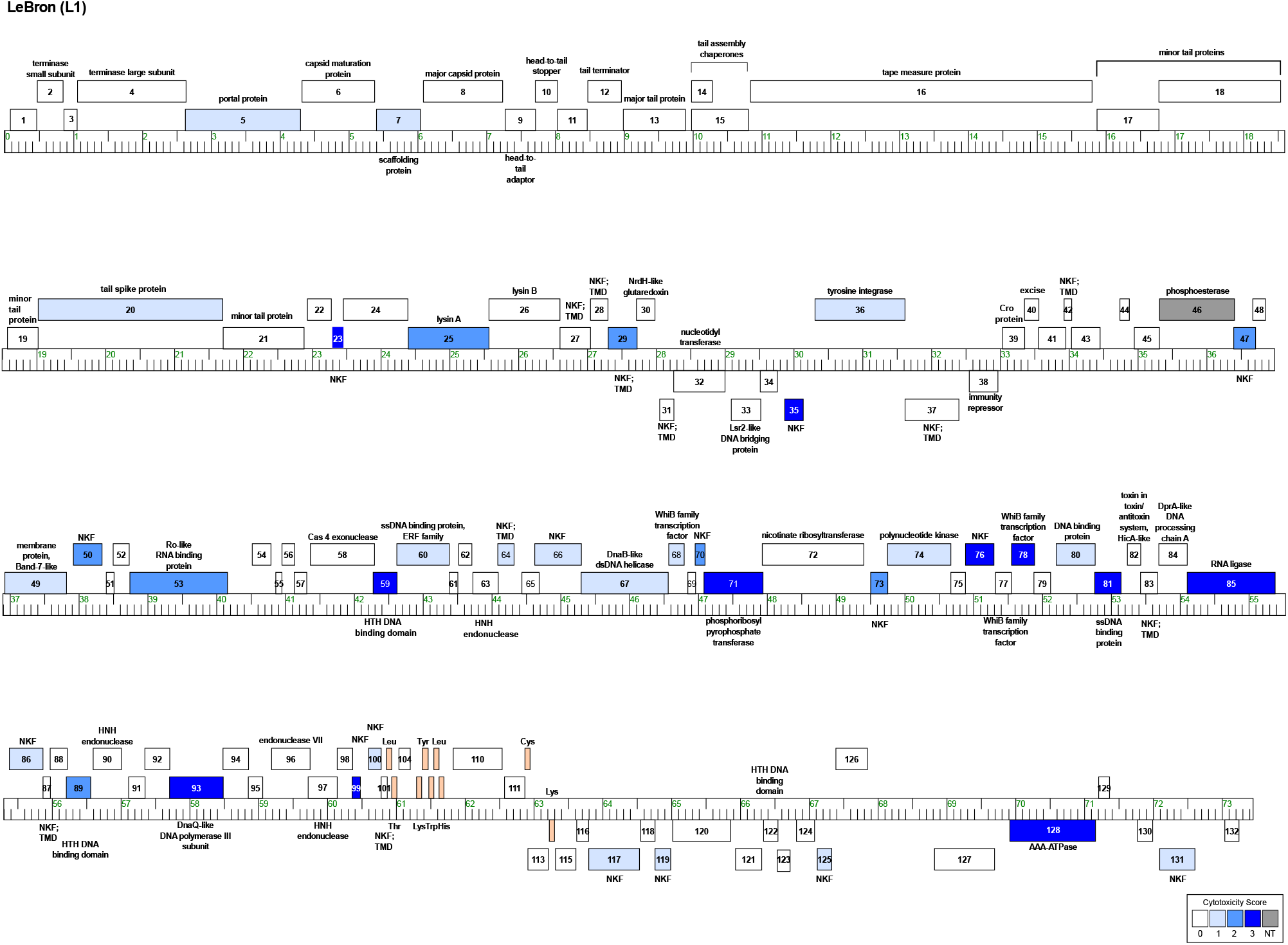
Genome map of phage Lebron highlighting cytotoxic genes. The linear genome of Lebron is shown with kilobase markers. Genes transcribed rightward are displayed above the line, and those transcribed leftward below. Numbers inside each box correspond to gene designations, with predicted functions labeled above or below. Box shading indicates cytotoxicity phenotypes: white, no effect on *M. smegmatis* growth (score 0); gray, no transformants recovered (NT); and blue, cytotoxic genes. Blue shading reflects severity: light blue (score *1*; reduction in colony size; genes *5, 7, 20, 36, 49, 60, 64, 66, 67, 68, 74, 80, 86, 100, 113, 117, 119, 125, 131*), medium blue (score *2*; 1–3 log reduction in viability; genes *25, 29, 47, 50, 53, 70, 73*, and *89*), and dark blue (score *3*; >3-log reduction in viability; genes *23, 35, 59, 71, 76, 78, 81, 85, 93, 99* and *128*).

In contrast, overexpression of 38 LeBron genes reduced *M. smegmatis* growth, with half of these (19/38) producing moderate to severe cytotoxic effects (Table 1; Fig. 3a). Overall, 32% of LeBron genes inhibited *M. smegmatis* growth, a frequency that closely parallels those observed for Amelie (34%) (Tafoya et al. 2025), Waterfoul (34%) (Heller et al. 2022), Hammy (26%) (Amaya et al. 2023), Girr (28%) (Pollenz et al. 2024), and NormanBulbieJr (28%) (Wise et al. 2025) (Fig. 3b). Among the cytotoxic genes, LeBron *99* (no known function) encoded the smallest toxic protein at only 39 amino acids, whereas LeBron *20* (tail spike protein) encoded the largest at 893 amino acids (Table 1).

**Table 1.**
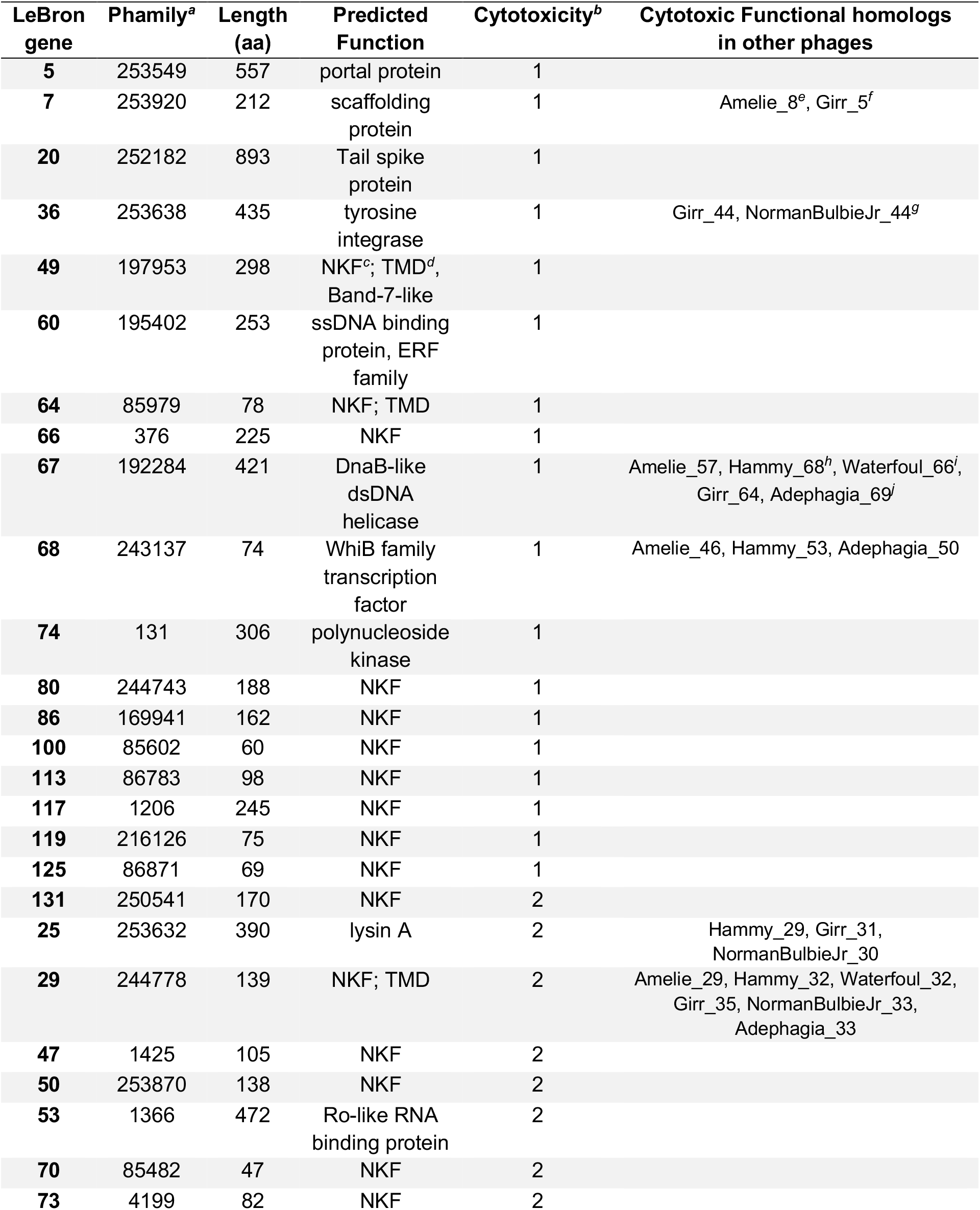
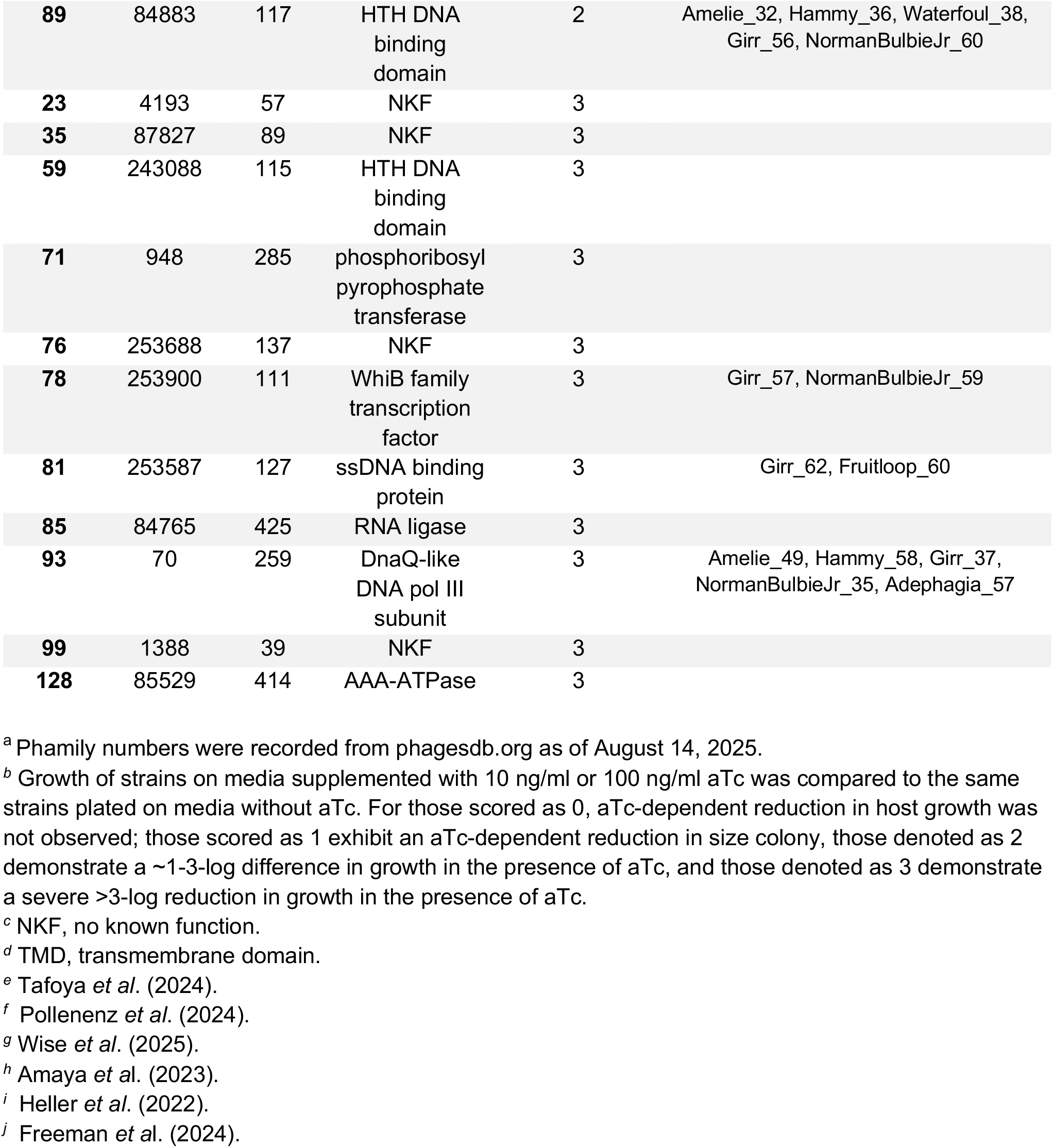
LeBron genes observed to inhibit *Mycobacterium smegmatis* growth upon overexpression.

**Figure 3.**
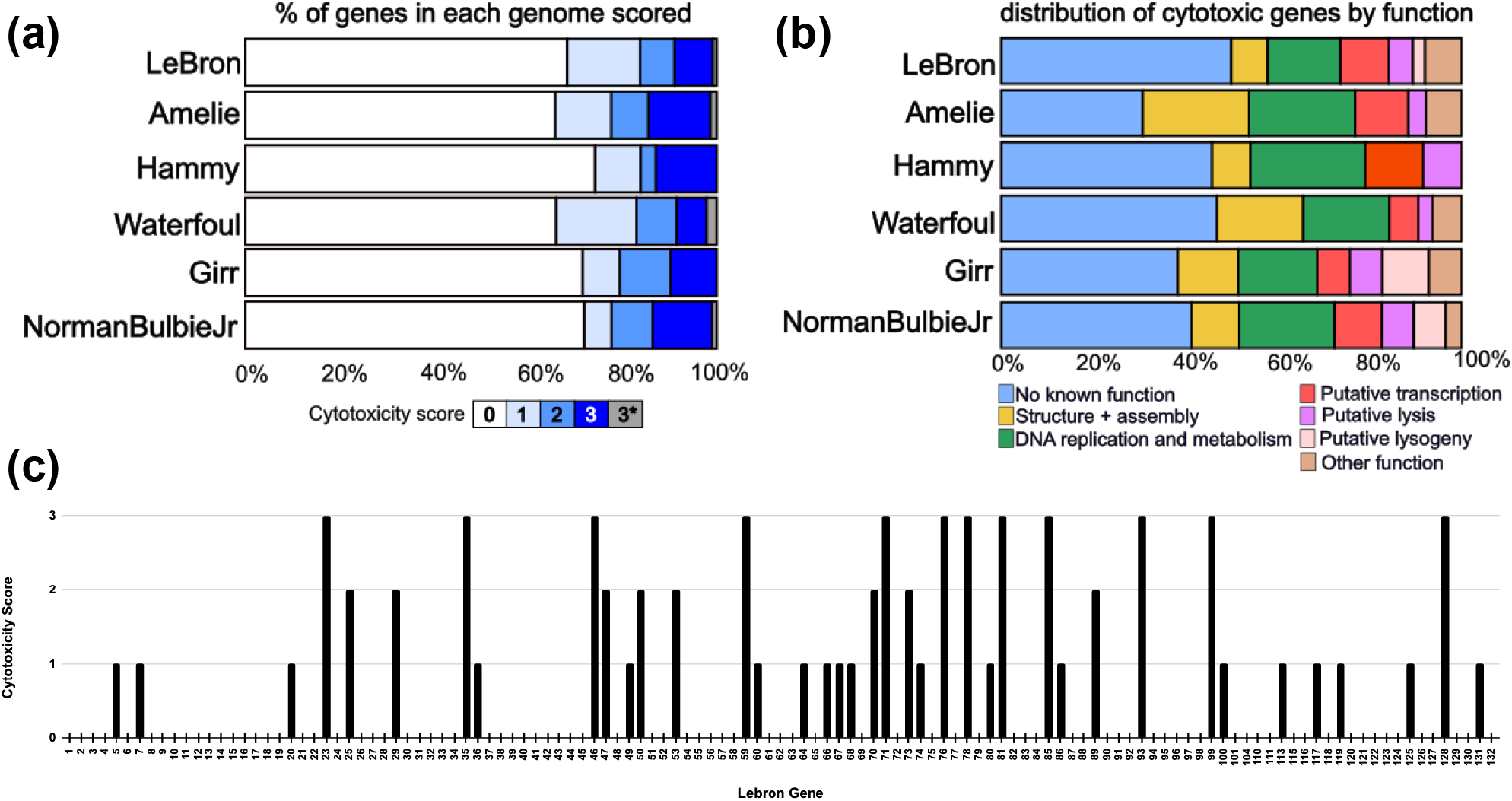
Comparative patterns of cytotoxicity across mycobacteriophages. a) Stacked bar chart showing the proportion of genes assigned to each cytotoxicity score (0–3) in LeBron (this study), Hammy (Amaya et al. 2023), Waterfoul (Heller et al. 2022), Girr (Pollenz et al. 2024), and NormanBoulbieJr (Wise et al. 2025). Genes for which no transformants could be recovered due to extreme toxicity (Lebron gp46, Amelie gp6, Waterfoul gp8, NormanBoulbieJr gp102) are designated as score 3*. b) Stacked bar chart showing the functional distribution of cytotoxic genes (scores 1–3) for each phage. Categories include NKF (no known function) and predicted functional classes, color-coded as indicated in the legend. c) Cytotoxicity scores (0–3) for each LeBron gene are plotted in genomic order along the horizontal axis, highlighting both the overall distribution of toxic genes and their tendency to cluster in specific regions of the genome.

Amongst the 38 LeBron genes identified as cytotoxic, 19 have predicted functions (Table 1). Three structural genes, the portal protein (gp5), the scaffolding protein (gp7), and the tail spike protein (gp20) produced mild cytotoxicity. LeBron encodes multiple lysis-associated proteins with distinct cytotoxic profiles. Lysin A (gp25) displayed moderate toxicity, whereas Lysin B (gp26) was not toxic. Immediately downstream, two small transmembrane proteins are encoded: gp28, which contains two predicted transmembrane helices but was not toxic in our assay, and gp29, which has a single predicted transmembrane helix and caused moderate toxicity. Phage lysis cassettes often encode multiple holin-like transmembrane proteins that act together to coordinate the timing of host cell lysis (Catalão et al. 2011; Pollenz et al. 2022). A recent study of *Corynebacterium glutamicum* phages revealed a conserved arrangement in which a two-TMD protein functions as the holin and a downstream single-TMD protein, termed LysZ, is essential for proper lysis (McKitterick et al. 2025). LeBron gp29 shares structural features, synteny, and cytotoxic phenotype with LysZ, suggesting it plays an analogous role in regulating cell lysis of *M. smegmatis*. Additionally, overexpression of the gene encoding the tyrosine integrase (gp36), which regulates the integration of the phage genome into the bacterial genome, resulted in a mild reduction in bacterial growth.

Six gene products with predicted roles in DNA replication or metabolism had an adverse impact on host growth. These included two single-stranded DNA binding proteins, gp60 and gp81, as well as two DNA helicases. gp67 encodes a DnaB-like dsDNA helicase, while the AAA+ ATPase gp128 showed a 99.9% probability match to the DNA helicase MCM6 in HHPRED (PBD ID: 6RAW), suggesting a potential role in DNA unwinding during genome replication (Malysa et al. 2025). In addition, the DnaQ-like DNA polymerase III subunit (gp93) and the phosphoribosyl pyrophosphate transferase (gp71), which participates in nucleotide biosynthesis (Hove-Jensen et al. 2016), were highly toxic.

Four cytotoxic LeBron genes are predicted to encode DNA-binding proteins that may regulate gene expression. Two belong to the WhiB family of transcription factors: overproduction of gp68 caused only mild toxicity, whereas gp78 was highly toxic. In addition, two proteins with predicted helix–turn–helix DNA-binding motifs, gp59 and gp89, were both highly toxic. Finally, three toxic proteins are predicted to function in RNA metabolism. gp53 encodes a Ro-like RNA-binding protein, gp85 is a predicted RNA ligase, and gp74 encodes a polynucleotide kinase, an enzyme class often involved in RNA or DNA repair (Wang et al. 2002).

The 19 toxic proteins of unknown function varied in severity, ranging from mild effects (gp49, gp64, gp66, gp80, gp86, gp100, gp113, gp117, gp119, gp125) to severe growth inhibition (gp23, gp47, gp50, gp54, gp70, gp73, gp76, gp99, gp131). Notably, some of the most toxic genes, gp23, gp47, gp54, gp70, gp73, and gp99, encode for small proteins under 100 amino acids. Several small phage-encoded proteins have recently been implicated in critical regulatory roles, including the modulation of host gene expression as well as directing lysogeny-lysis transitions (Beggs and Bassler 2024).

Overall, the distribution of toxic genes in LeBron underscores the broad functional diversity of phage products capable of perturbing host physiology. As seen in other phages, replication-associated proteins and certain transcriptional regulators are among the most potent growth inhibitors, but LeBron also encodes a number of poorly characterized small proteins with strong cytotoxicity, making them intriguing candidates for future mechanistic studies.

### Conservation of mycobacterial growth inhibition by related phage genes

Results from the LeBron overexpression screen highlight both conserved and distinctive features when compared with previously characterized phages from Clusters K and F1. At the nucleotide level, LeBron shares only 9.4–9.5% gene content similarity (GCS) with Cluster K phages and just 3.6% with Cluster F1 phages, underscoring its divergence in overall genome composition. Despite this low sequence similarity, the proportion of genes that were cytotoxic was comparable across all phages examined (Fig. 3a), as was the functional distribution of these toxic genes (Fig. 3b). This suggests that while LeBron and Cluster K and F1 phages are genetically distinct, they converge on a common set of gene functions that impair *M. smegmatis* growth when overexpressed. To further explore this relationship, we compared the cytotoxic phenotypes of LeBron gene products with those of functional homologs in Cluster K and Cluster F phages.

In each of the genomes screened to date, a variable subset of structural proteins has been observed to cause growth inhibition upon overexpression. Notably, unlike the Cluster K and F1 phages, LeBron did not show toxicity associated with its major capsid protein. Three structural proteins in LeBron were cytotoxic, the portal protein (gp5), the scaffolding protein (gp7), and the tail spike protein gp20. The scaffolding proteins of LeBron (gp7), Amelie (gp8), and Girr (gp5) were toxic to varying degrees, despite sharing less than 26% amino acid identity. LeBron gp5 represents the first reported instance of a portal protein causing toxicity. Although it belongs to the same phamily as Adephagia gp9, Amelie gp5, Hammy gp18, and Waterfoul gp7, portal proteins in Cluster K phages share high identity with one another (68–83%) but are much more divergent from LeBron gp5 (28–30% identity). At least one predicted component of the phage tail structures from Lebron, Waterfoul, Amelie, Girr, and NormanBulbieJr has been reported to cause growth inhibition upon cytoplasmic overexpression. Recent cryo-EM reconstructions of the Bxb1 tail during host infection have provided a comprehensive view of the mycobacteriophage siphoviral tail apparatus, allowing for more specific functional predictions of tail associated genes (Freeman et al. 2025). Based on these new structural insights, Lebron gp20 is predicted to be the tail spike protein, with a >99.9% probability HHPRED match to the Bxb1 tail spike protein gp29 (PDB ID: 9D93_Pc), whereas the cytotoxic tail proteins encoded by Girr (gp19) and NormanBulbieJr (gp19) match with >99.9% probability to the Bxb1 tail wing base and contain predicted D-ala-D-ala carboxypeptidase catalytic domains. Waterfoul gp21on the other hand is predicted to be the baseplate hub protein, whereas Amelie gp20 does not match to any of the resolved features on the Bxb1 tail. Most of these structural proteins cause only mild to moderate growth inhibition when overproduced in the cytoplasm, and we would expect that their primary role in host perturbation would occur at the cell surface during infection; however, these inter-cluster differences underscore the wide range of structural proteins capable of impairing *M. smegmatis* growth.

DNA replication and metabolism-associated proteins were also cytotoxic across clusters. Previous work has shown that forced accumulation of replisome components leads to abnormal DNA replication (Roseaulin et al. 2013). Moreover, polymerases and single-stranded DNA-binding proteins have recently been implicated in interactions with bacterial anti-phage defense systems (Stokar-Avihail et al. 2023), suggesting that their toxicity may reflect not only disruption of replication stoichiometry but also antagonism of host defense pathways. LeBron gp93, a DnaQ-like DNA polymerase III subunit, was among the most toxic, consistent with related homologs in Amelie (gp49), Hammy (gp58), Adephagia (gp57) and Girr (gp37). The DnaQ-like DNA polymerase III subunit acts as a 3’ →5’ exonuclease that removes incorrectly incorporated nucleotides, thereby increasing the fidelity of DNA replication (Scheuermann and Echols 1984). LeBron gp67, a DnaB-like helicase, caused only mild toxicity, a phenotype that closely parallels Adephagia gp69, Amelie gp57, Girr gp64, Hammy gp68, and Waterfoul gp66. Additionally, based on structural similarity to DNA helicase MCM6, the AAA+ ATPase gp128 also has a potential role in DNA unwinding during genome replication. Two single-stranded DNA-binding proteins, gp60 and gp81, were also toxic, consistent with similar phenotypes reported for Girr gp62 and Fruitloop gp60 (Ko and Hatfull 2020; Pollenz et al. 2024 Mar 8). Finally, LeBron gp71, a phosphoribosyl pyrophosphate transferase, was highly toxic, underscoring the essential role of nucleotide biosynthesis in supporting both host and phage replication (Hove-Jensen et al. 2016). Additional investigation is needed to understand whether cytotoxicity by phage replication proteins is due to interference with the host DNA or replication machinery or due to another mechanism.

Four LeBron proteins predicted to regulate transcription were cytotoxic. Among its three WhiB family transcription factors, gp77 was not toxic, gp68 caused mild toxicity, and gp78 was highly toxic. The difference in phenotype is likely explained by the fact that these proteins share only 18-32% amino acid identity. Although there’s not significant sequence similarity to LeBron gp68, WhiB family proteins from Amelie (gp46), Hammy (gp53), and Adephagia (gp50) also showed mild reduction in host growth. In contrast, Girr gp57 and NormanBulbieJr gp 59 were highly toxic, paralleling the phenotype observed for LeBron gp78. WhiB proteins are known to function in cooperation with other proteins and play central roles in processes such as cell division, development, antibiotic resistance and virulence (Bush 2018). Supporting this, the WhiB-like protein of mycobacteriophage TM4 was shown to inhibit the essential *M. smegmatis* WhiB2 by binding to the whiB2 promoter and downregulating its expression, thereby interfering with septation and cell division (Rybniker et al. 2010). These findings suggest that LeBron gp68 and gp78 may impair host growth through similar disruption of WhiB-regulated pathways. In addition, LeBron encodes two helix–turn–helix (HTH) DNA-binding proteins, gp59 and gp89. Gp89 caused moderate toxicity, while gp59 was highly toxic. By contrast, homologous HTH proteins from Amelie (gp32), Hammy (gp36), Girr (gp56), NormanBulbieJr (gp60), and Waterfoul (gp38) all produced only mild phenotypes. This suggests that while HTH DNA-binding proteins are broadly cytotoxic across phages, the magnitude of their effects may vary depending on sequence divergence or host interactions.

The lysis cassette of LeBron encodes a lysin A (gp25) that was moderately toxic, a lysin B (gp26) that was not toxic, and two small transmembrane proteins immediately downstream. gp28, which has two predicted transmembrane domains, was not toxic, whereas gp29, which has a single predicted transmembrane domain, displayed moderate toxicity. A similar lysis cassette organization is observed in Cluster F1 phages Girr and NormanBulbieJr, where both the lysin A and the single-TMD proteins were cytotoxic but lysin B and a double-TMD protein were not toxic (Pollenz et al. 2024; Wise et al. 2025). The arrangement in Cluster K phages differs slightly: lysin A and lysin B are followed by a four TMD protein and then a single-TMD protein. Among the lysin A proteins only Hammy gp29 showed toxicity, whereas the single-TMD proteins in all Cluster K phages examined (Amelie gp29, Hammy gp32, Waterfoul gp32, and Adephagia gp33) were all toxic. The seven single-TMD proteins found to be consistently cytotoxic across these screens belong to three different gene phamilies, signifying their low sequence identity. Recent work on *Corynebacterium glutamicum* phages described an analogous lysis cassette configuration in which a single-TMD protein, termed LysZ, located downstream of lysin genes and a multi-TMD holin, was found to be cytotoxic to the host upon overexpression and essential for host lysis (McKitterick et al. 2025). The conserved synteny and cytotoxicity of these single-TMD proteins despite having low sequence conservation, being from diverse phage clusters, and even targeting different host bacteria suggest that they may play a key conserved role in regulating lysis of the complex actinobacterial cell envelope.

LeBron’s tyrosine integrase (gp36), which regulates the integration of the phage genome into the bacterial genome, was mildly toxic, similar to related pham members Girr gp44 which was moderately toxic and NormanBulbieJr gp44 which was highly toxic. Interestingly, Hammy gp42, which belongs to the same phamily and shares 45% amino acid identity with these integrases, did not cause toxicity.

Of note, three toxic LeBron genes are predicted to function in RNA metabolism, a category not previously reported for other phages. These include gp53, a Ro-like RNA-binding protein; gp85, an RNA ligase; and gp74, a polynucleotide kinase. In phage T4, RNA ligase plays key roles in RNA repair, splicing, and editing pathways (Amitsur et al. 1987; Gonzalez et al. 1999). Together with T4 polynucleotide kinase, the RNA ligase participates in an RNA repair system that restores functionality to damaged tRNAs by remodeling and sealing broken RNA ends (Amitsur et al. 1987). The presence of analogous functions in LeBron suggests that RNA repair or modification may represent a conserved but previously underappreciated strategy by which phages manipulate host RNA metabolism. Another notable deviation observed in the Lebron dataset was the absence of toxicity observed with overexpression of gene *82* which encodes a predicted HicA-like ribonuclease toxin. Toxin-antitoxin (TA) systems are abundant with bacterial and phage genomes (LeRoux and Laub 2022), and other ribonuclease toxins—such as the HicA-like toxin gp73 from Amelie, the HicA-like toxin gp90 from Adephagia, and the BrnT-like toxin gp86 from Waterfoul—exhibit strong cytotoxicity in comparable overexpression assays.

Lebron gp82 shares 44-49% sequence identity with the HicA-like toxins from Adephagia and Amelie, but unlike them, appears to be an orphan toxin, without a predicted cognate antitoxin encoded nearby. Therefore, although differences in expression levels across constructs cannot be excluded, this finding suggests that phage-encoded TA systems can vary widely in activity, even among homologs with significant sequence similarity.

Overall, across different mycobacteriophage clusters, patterns of phenotypic conservation are complex; however, we consistently see common functional themes emerge, regardless of whether proteins have high sequence similarity, thereby supporting that functional category, rather than precise sequence conservation, can often be a better predictor of overexpression phenotypes.

### Additional insights

This study represents the first genome-wide screen for cytotoxic genes from a Cluster L phage. Despite LeBron’s limited gene content overlap with Cluster K and F1 phages, a consistent pattern emerged: mycobacteriophages are rich in cytotoxic genes. Across genomes tested to date, roughly one-quarter to one-third of genes impaired host growth when overexpressed (Heller et al. 2022; Freeman et al. 2024; Pollenz et al. 2024; Tafoya et al. 2025; Wise et al. 2025). While some effects, particularly those scored as mild (score 1), may reflect artifacts of overexpression rather than biologically meaningful interactions, more than half of LeBron’s cytotoxic genes (52%), including 10 with no known function, produced moderate to severe phenotypes even at intermediate inducer concentrations (Table 1). These are especially compelling candidates for future functional analysis. Another notable trend is that cytotoxic genes, though dispersed across the genome, frequently occur in close proximity, often within the same predicted operon. This clustering suggests possible cooperative functions during different stages of the phage life cycle. In LeBron, three such clusters were evident (Fig. 2 and 3c): Cluster 1 (genes 46, 47, 49, 50, 53, 54), Cluster 2 (genes 64, 66, 67, 68, 70, 71), and Cluster 3 (genes 73, 74, 76, 78, 80, 81). These regions contain both proteins of unknown function and those predicted to participate in DNA replication, DNA metabolism, transcription, and RNA metabolism, highlighting them as genomic hotspots of cytotoxic activity.

Out of 123 LeBron genes tested, 38 showed measurable toxicity when expressed in *M. smegmatis*. Additionally, gp46 was so toxic that no transformants could be recovered despite multiple attempts with different pExtra constructs. This extreme phenotype suggests that leaky expression of gp46, even in the absence of inducer, is sufficient to kill host cells. Similar outcomes have been reported in other systems, including Amelie gp6, Waterfoul gp8, and NormanBulbieJr gp102, where no colonies could be recovered due to toxicity (Heller et al. 2022; Tafoya et al. 2025; Wise et al. 2025). In the case of Waterfoul gp8, a nonsense mutation restored transformant recovery, confirming that low-level protein production underlies the phenotype (Heller et al. 2022). Based on these parallels, LeBron gp46 is designated as a toxic gene (scored as 3* in Fig. 3a), bringing the total number of cytotoxic genes in LeBron to 39.

The patterns of cytotoxicity observed for LeBron’s genes reinforce the emerging view that functional conservation, rather than pham assignment or high sequence identity, is the best predictor of growth inhibition in *M. smegmatis* upon gene overexpression. Across diverse clusters, structural proteins, lysis-associated genes, replication and repair enzymes, transcriptional regulators, lysogeny-associated integrases, RNA metabolism enzymes, and many proteins of unknown function repeatedly displayed cytotoxicity. Notably, several highly toxic LeBron proteins lack characterized homologs yet mirror phenotypes reported in Cluster K and F1 phages, underscoring the importance of functionally conserved strategies despite extensive sequence divergence.

Recent advances in structure-based homology tools, such as those demonstrated by Guo *et al*. (Guo and He 2025) are revealing distant functional relationships that escape detection by sequence-based comparisons and may provide more powerful predictors of protein function and phenotype. Altogether, our findings highlight the dual nature of phage evolution: remarkable genetic diversity coupled with phenotypic redundancy in how phages manipulate their bacterial hosts. Such patterns not only expand our understanding of phage–host interactions but also point to new avenues for uncovering gene functions and identifying phage-derived proteins with potential as antimicrobial tools.

## Data availability statement

All plasmids and plasmid sequences reported in this study are available upon request. The authors affirm that all data necessary for confirming the conclusions of this article are represented fully within the article and its tables and figures. Extended data, including plasmid Sanger sequencing data and confirmatory cytotoxicity assay data can be found at the SEA-GENES project open access database GenesDB (https://genesdb.org). To access these data on genesDB.org, any user can register for a free account, and once logged in to this account, navigate from the home page to Cluster L1 phage LeBron.

Supplementary material is available at G3 online.

## Acknowledgements

We are grateful to the members of the Science Education Alliance for their invaluable research support, particularly Viknesh Sivanathan, Graham Hatfull, Deborah Jacobs-Sera, Steven Cresawn, and Dan Russell. We also thank New England Biolabs (NEB) and Integrated DNA Technologies (IDT) for generously providing reagents. We would also like to thank the Biology Department and the College of Arts and Sciences at La Sierra University for their continued support of SEA programs on campus.

