## Supplemental Figure 1 and Table 1 for "Functional Characterization of 123 Genes from Mycobacteriophage LeBron Uncovers Cytotoxic Gene Families in Cluster L1"

#### Gene *1*; Score *0*

Images taken after 5 days at 37 ºC

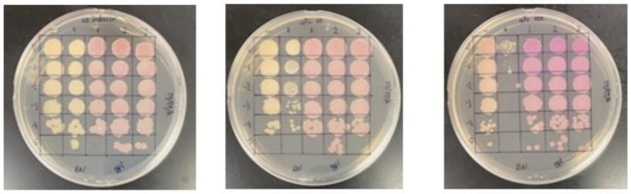


0 ng/ml aTc | 10 ng/ml aTc | 100 ng/ml aTc

| Lane | Gene ID | Plasmid name | Gene name | Toxic/Non-toxic | Colony color on 100 ng/ml aTc plate* |
|---|---|---|---|---|---|
| + Toxic control | -- | pExTra02 | Fruitloop *52* | Toxic | - |
| - Non-toxic control | -- | pExTra03 | Fruitloop *52* mutant | Non-toxic | + |
| 1 | | pExTra-Lebron 1 | Lebron *1* replicate 1 | Non-toxic | +++ |
| 2 | | pExTra-Lebron 1 | Lebron *1* replicate 2 | Non-toxic | +++ |
| 3 | | pExTra-Lebron 1 | Lebron *1* replicate 3 | Non-toxic | +++ |

*Key: NG (no growth) - (no pink color) +(faint pink color) ++(obvious pink color) +++ (dark pink color)

#### Gene *2*; Score *0*

Images taken after 5 days at 37 ºC

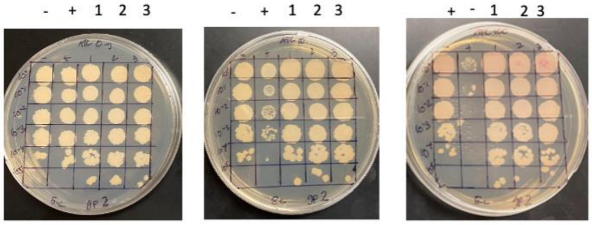


0 ng/ml aTc | 10 ng/ml aTc | 100 ng/ml aTc

| Lane | Gene ID | Plasmid name | Gene name | Toxic/Non-toxic | Colony color on 100 ng/ml aTc plate* |
|---|---|---|---|---|---|
| + Toxic control | -- | pExTra02 | Fruitloop *52* | Toxic | - |
| - Non-toxic control | -- | pExTra03 | Fruitloop *52* mutant | Non-toxic | + |
| 1 | | pExTra-Lebron 2 | Lebron 2 replicate 1 | Non-toxic | ++ |
| 2 | | pExTra-Lebron 2 | Lebron 2 replicate 2 | Non-toxic | ++ |
| 3 | | pExTra-Lebron 2 | Lebron 2 replicate 3 | Non-toxic | ++ |

*Key: NG (no growth) - (no pink color) +(faint pink color) ++(obvious pink color) +++ (dark pink color)

#### Gene *3*; Score *0*

Images taken after 5 days at 37 ºC

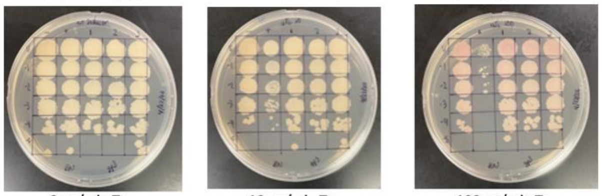


0 ng/ml aTc | 10 ng/ml aTc | 100 ng/ml aTc

| Lane | Gene ID | Plasmid name | Gene name | Toxic/Non-toxic | Colony color on 100 ng/ml aTc plate* |
|---|---|---|---|---|---|
| + Toxic control | -- | pExTra02 | Fruitloop *52* | Toxic | - |
| - Non-toxic control | -- | pExTra03 | Fruitloop *52* mutant | Non-toxic | + |
| 1 | | pExTra-Lebron 3 | Lebron *3* replicate 1 | Non-toxic | + |
| 2 | | pExTra-Lebron 3 | Lebron *3* replicate 2 | Non-toxic | + |
| 3 | | pExTra-Lebron 3 | Lebron *3* replicate 3 | Non-toxic | + |

*Key: NG (no growth) - (no pink color) +(faint pink color) ++(obvious pink color) +++ (dark pink color)

#### Gene *4*; Score *0*

Images taken after 5 days at 37 ºC

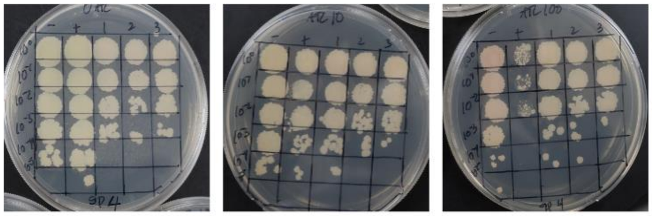


0 ng/ml aTc | 10 ng/ml aTc | 100 ng/ml aTc

| Lane | Gene ID | Plasmid name | Gene name | Toxic/Non-toxic | Colony color on 100 ng/ml aTc plate* |
|---|---|---|---|---|---|
| + Toxic control | -- | pExTra02 | Fruitloop *52* | Toxic | - |
| - Non-toxic control | -- | pExTra03 | Fruitloop *52* mutant | Non-toxic | + |
| 1 | | pExTra-Lebron 4 | Lebron *4* replicate 1 | Non-toxic | + |
| 2 | | pExTra-Lebron 4 | Lebron *4* replicate 2 | Non-toxic | + |
| 3 | | pExTra-Lebron 4 | Lebron *4* replicate 3 | Non-toxic | + |

*Key: NG (no growth) - (no pink color) +(faint pink color) ++(obvious pink color) +++ (dark pink color)

#### Gene *5*; Score *1*

Images taken after 5 days at 37 ºC

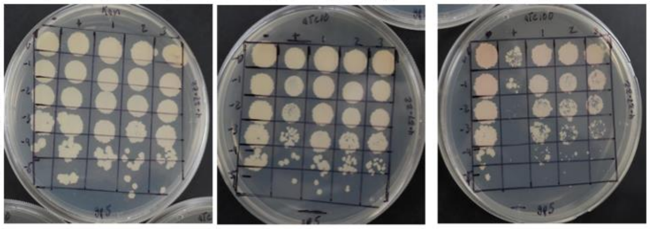


0 ng/ml aTc | 10 ng/ml aTc | 100 ng/ml aTc

| Lane | Gene ID | Plasmid name | Gene name | Toxic/Non-toxic | Colony color on 100 ng/ml aTc plate* |
|---|---|---|---|---|---|
| + Toxic control | -- | pExTra02 | Fruitloop *52* | Toxic | - |
| - Non-toxic control | -- | pExTra03 | Fruitloop *52* mutant | Non-toxic | + |
| 1 | 131436 | pExTra-Lebron 5 | Lebron *5* replicate 1 | Toxic | ++ |
| 2 | 131436 | pExTra-Lebron 5 | Lebron *5* replicate 2 | Toxic | ++ |
| 3 | 131436 | pExTra-Lebron 5 | Lebron *5* replicate 3 | Toxic | ++ |

*Key: NG (no growth) - (no pink color) +(faint pink color) ++(obvious pink color) +++ (dark pink color)

#### Gene *6*; Score *0*

Images taken after 5 days at 37 ºC

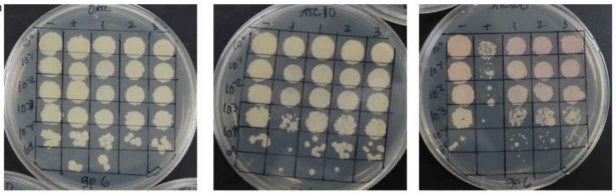


0 ng/ml aTc | 10 ng/ml aTc | 100 ng/ml aTc

| Lane | Gene ID | Plasmid name | Gene name | Toxic/Non-toxic | Colony color on 100 ng/ml aTc plate* |
|---|---|---|---|---|---|
| - Non-toxic control | -- | pExTra03 | Fruitloop *52* mutant | Non-toxic | + |
| + Toxic control | -- | pExTra02 | Fruitloop *52* | Toxic | - |
| 1 | | pExTra-Lebron 6 | Lebron *6* replicate 1 | Non-toxic | ++ |
| 2 | | pExTra-Lebron 6 | Lebron *6* replicate 2 | Non-toxic | ++ |
| 3 | | pExTra-Lebron 6 | Lebron *6* replicate 3 | Non-toxic | ++ |

*Key: NG (no growth) - (no pink color) +(faint pink color) ++(obvious pink color) +++ (dark pink color)

#### Gene *7*; Score *1*

Images taken after 5 days at 37 ºC

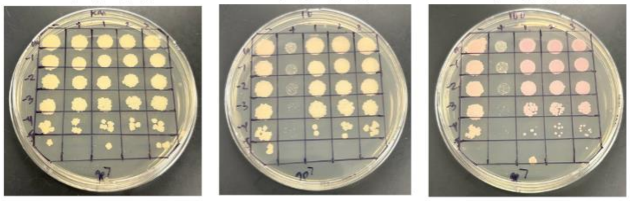


0 ng/ml aTc | 10 ng/ml aTc | 100 ng/ml aTc

| Lane | Gene ID | Plasmid name | Gene name | Toxic/Non-toxic | Colony color on 100 ng/ml aTc plate* |
|---|---|---|---|---|---|
| - Non-toxic control | -- | pExTra03 | Fruitloop *52* mutant | Non-toxic | + |
| + Toxic control | -- | pExTra02 | Fruitloop *52* | Toxic | - |
| 1 | | pExTra-Lebron 7 | Lebron *7* replicate 1 | Toxic | ++ |
| 2 | | pExTra-Lebron 7 | Lebron *7* replicate 2 | Toxic | ++ |
| 3 | | pExTra-Lebron 7 | Lebron *7* replicate 3 | Toxic | ++ |

*Key: NG (no growth) - (no pink color) +(faint pink color) ++(obvious pink color) +++ (dark pink color)

#### Gene *8*; Score *0*

Images taken after 5 days at 37 ºC

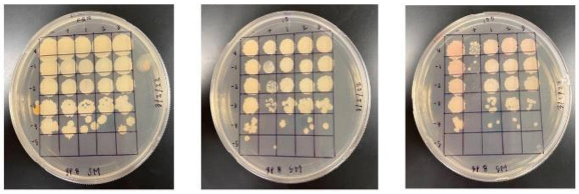


0 ng/ml aTc | 10 ng/ml aTc | 100 ng/ml aTc

| Lane | Gene ID | Plasmid name | Gene name | Toxic/Non-toxic | Colony color on 100 ng/ml aTc plate* |
|---|---|---|---|---|---|
| + Toxic control | -- | pExTra02 | Fruitloop *52* | Toxic | - |
| - Non-toxic control | -- | pExTra03 | Fruitloop *52* mutant | Non-toxic | + |
| 1 | | pExTra-Lebron 8 | Lebron *8* replicate 1 | Non-Toxic | + |
| 2 | | pExTra-Lebron 8 | Lebron *8* replicate 2 | Non-Toxic | + |
| 3 | | pExTra-Lebron 8 | Lebron 8 replicate 3 | Non-Toxic | + |

*Key: NG (no growth) - (no pink color) +(faint pink color) ++(obvious pink color) +++ (dark pink color)

### Gene *9*; Score *0*

Images taken after 5 days at 37 ºC

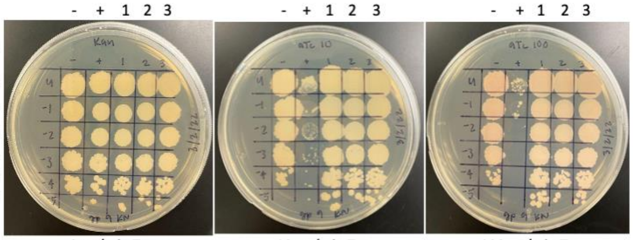


0 ng/ml aTc | 10 ng/ml aTc | 100 ng/ml aTc

| Lane | Gene ID | Plasmid name | Gene name | Toxic/Non-toxic | Colony color on 100 ng/ml aTc plate* |
|---|---|---|---|---|---|
| - Non-toxic control | -- | pExTra03 | Fruitloop *52* mutant | Non-toxic | + |
| + Toxic control | -- | pExTra02 | Fruitloop *52* | Toxic | - |
| 1 | | pExTra-Lebron 9 | Lebron 9 replicate 1 | Non-toxic | + |
| 2 | | pExTra-Lebron 9 | Lebron 9 replicate 2 | Non-toxic | + |
| 3 | | pExTra-Lebron 9 | Lebron 9 replicate 3 | Non-toxic | + |

*Key: NG (no growth) - (no pink color) +(faint pink color) ++(obvious pink color) +++ (dark pink color)

### Gene *10*; Score *0*

Images taken after 5 days at 37 ºC

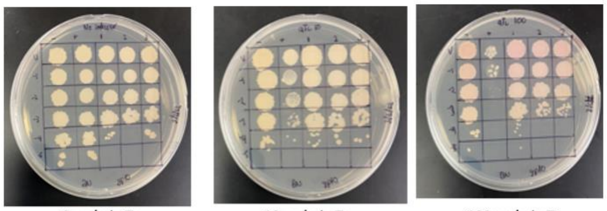


0 ng/ml aTc | 10 ng/ml aTc | 100 ng/ml aTc

| Lane | Gene ID | Plasmid name | Gene name | Toxic/Non-toxic | Colony color on 100 ng/ml aTc plate* |
|---|---|---|---|---|---|
| - Non-toxic control | -- | pExTra03 | Fruitloop *52* mutant | Non-toxic | + |
| + Toxic control | -- | pExTra02 | Fruitloop *52* | Toxic | + |
| 1 | | pExTra-Lebron 10 | Lebron *10* replicate 1 | Non-toxic | ++ |
| 2 | | pExTra-Lebron 10 | Lebron *10* replicate 2 | Non-toxic | ++ |
| 3 | | pExTra-Lebron 10 | Lebron *10* replicate 3 | Non-toxic | ++ |

*Key: NG (no growth) - (no pink color) +(faint pink color) ++(obvious pink color) +++ (dark pink color)

### Gene *11*; Score *0*

Images taken after 5 days at 37 ºC

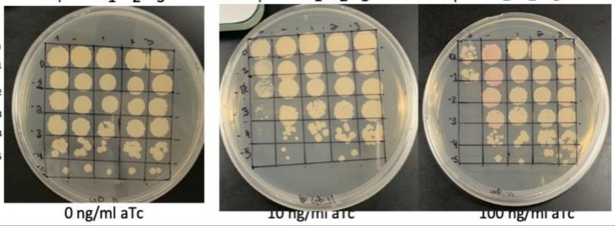


0 ng/ml aTc | 10 ng/ml aTc | 100 ng/ml aTc

| Lane | Gene ID | Plasmid name | Gene name | Toxic/Non-toxic | Colony color on 100 ng/ml aTc plate* |
|---|---|---|---|---|---|
| + Toxic control | -- | pExTra02 | Fruitloop *52* | Toxic | - |
| - Non-toxic control | -- | pExTra03 | Fruitloop *52* mutant | Non-toxic | ++ |
| 1 | | pExTra-Lebron 11 | Lebron *11* replicate 1 | Non-toxic | - |
| 2 | | pExTra-Lebron 11 | Lebron *11* replicate 2 | Non-toxic | - |
| 3 | | pExTra-Lebron 11 | Lebron *11* replicate 3 | Non-toxic | - |

*Key: NG (no growth) - (no pink color) +(faint pink color) ++(obvious pink color) +++ (dark pink color)

### Gene *12*; Score *0*

Images taken after 5 days at 37 ºC

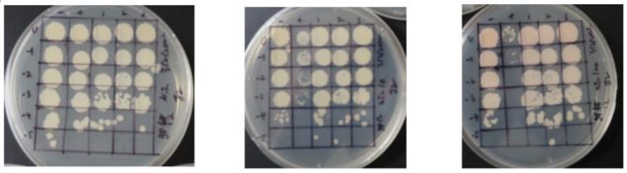


0 ng/ml aTc | 10 ng/ml aTc | 100 ng/ml aTc

| Lane | Gene ID | Plasmid name | Gene name | Toxic/Non-toxic | Colony color on 100 ng/ml aTc plate* |
|---|---|---|---|---|---|
| - Non-toxic control | -- | pExTra03 | Fruitloop *52* mutant | Non-toxic | + |
| + Toxic control | -- | pExTra02 | Fruitloop *52* | Toxic | - |
| 1 | | pExTra-Lebron 12 | Lebron *12* replicate 1 | Non-toxic | + |
| 2 | | pExTra-Lebron 12 | Lebron *12* replicate 2 | Non-toxic | + |
| 3 | | pExTra-Lebron 12 | Lebron *12* replicate 3 | Non-toxic | + |

*Key: NG (no growth) - (no pink color) +(faint pink color) ++(obvious pink color) +++ (dark pink color)

### Gene *13*; Score *0*

Images taken after 5 days at 37 ºC

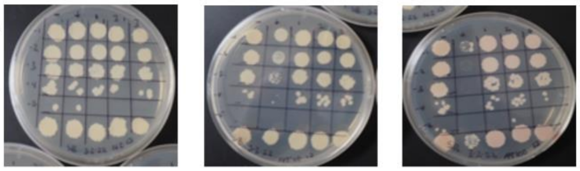


0 ng/ml aTc | 10 ng/ml aTc | 100 ng/ml aTc

| Lane | Gene ID | Plasmid name | Gene name | Toxic/Non-toxic | Colony color on 100 ng/ml aTc plate* |
|---|---|---|---|---|---|
| + Toxic control | -- | pExTra02 | Fruitloop *52* | Non-toxic | + |
| - Non-toxic control | -- | pExTra03 | Fruitloop *52* mutant | Toxic | - |
| 1 | | pExTra-Lebron 13 | Lebron 13 replicate 1 | Non-toxic | + |
| 2 | | pExTra-Lebron 13 | Lebron 13 replicate 2 | Non-toxic | + |
| 3 | | pExTra-Lebron 13 | Lebron 13 replicate 3 | Non-toxic | + |

*Key: NG (no growth) - (no pink color) +(faint pink color) ++(obvious pink color) +++ (dark pink color)

### Gene *14*; Score *0*

Images taken after 5 days at 37 ºC

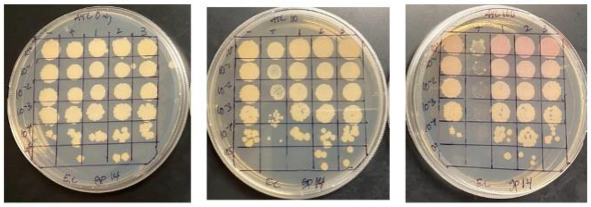


0 ng/ml aTc | 10 ng/ml aTc | 100 ng/ml aTc

| Lane | Gene ID | Plasmid name | Gene name | Toxic/Non-toxic | Colony color on 100 ng/ml aTc plate* |
|---|---|---|---|---|---|
| + Toxic control | -- | pExTra02 | Fruitloop *52* | Toxic | - |
| - Non-toxic control | -- | pExTra03 | Fruitloop *52* mutant | Non-toxic | + |
| 1 | | pExTra-Lebron 14 | Lebron 14 replicate 1 | Non-toxic | ++ |
| 2 | | pExTra-Lebron 14 | Lebron 14 replicate 2 | Non-toxic | ++ |
| 3 | | pExTra-Lebron 14 | Lebron 14 replicate 3 | Non-toxic | ++ |

*Key: NG (no growth) - (no pink color) +(faint pink color) ++(obvious pink color) +++ (dark pink color)

### Gene *15*; Score *0*

Images taken after 5 days at 37 ºC

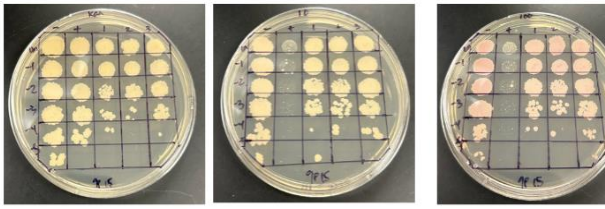


0 ng/ml aTc | 10 ng/ml aTc | 100 ng/ml aTc

| Lane | Gene ID | Plasmid name | Gene name | Toxic/Non-toxic | Colony color on 100 ng/ml aTc plate* |
|---|---|---|---|---|---|
| - Non-toxic control | -- | pExTra03 | Fruitloop *52* mutant | Non-toxic | ++ |
| + Toxic control | -- | pExTra02 | Fruitloop *52* | Toxic | - |
| 1 | | pExTra-Lebron 15 | Lebron *15* replicate 1 | Non-toxic | + |
| 2 | | pExTra-Lebron 15 | Lebron *15* replicate 2 | Non-toxic | + |
| 3 | | pExTra-Lebron 15 | Lebron *15* replicate 3 | Non-toxic | + |

*Key: NG (no growth) - (no pink color) +(faint pink color) ++(obvious pink color) +++ (dark pink color)

### Gene *16*; Score *0*

Images taken after 5 days at 37 ºC

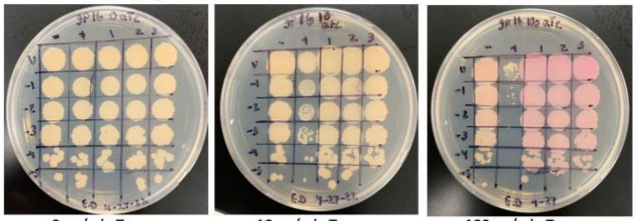


0 ng/ml aTc | 10 ng/ml aTc | 100 ng/ml aTc

| Lane | Gene ID | Plasmid name | Gene name | Toxic/Non-toxic | Colony color on 100 ng/ml aTc plate* |
|---|---|---|---|---|---|
| + Toxic control | -- | pExTra02 | Fruitloop *52* | Toxic | - |
| - Non-toxic control | -- | pExTra03 | Fruitloop *52* mutant | Non-toxic | ++ |
| 1 | | pExTra-Lebron 16 | Lebron *16* replicate 1 | Non-toxic | +++ |
| 2 | | pExTra-Lebron 16 | Lebron *16* replicate 2 | Non-toxic | +++ |
| 3 | | pExTra-Lebron 16 | Lebron *16* replicate 3 | Non-toxic | +++ |

*Key: NG (no growth) - (no pink color) +(faint pink color) ++(obvious pink color) +++ (dark pink color)

#### Gene *17*; Score *0*

Images taken after 5 days at 37 ºC

- + 1 2 3 (each plate)

Dilution factor: $10^0$, $10^{-1}$, $10^{-2}$, $10^{-3}$, $10^{-4}$, $10^{-5}$

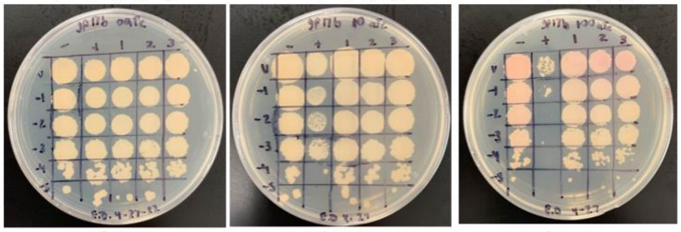

0 ng/ml aTc | 10 ng/ml aTc | 100 ng/ml aTc

| Lane | Gene ID | Plasmid name | Gene name | Toxic/Non-toxic | Colony color on 100 ng/ml aTc plate* |
|---|---|---|---|---|---|
| + Toxic control | -- | pExTra02 | Fruitloop *52* | Toxic | - |
| - Non-toxic control | -- | pExTra03 | Fruitloop *52* mutant | Non-toxic | + |
| 1 | | pExTra-LeBron 17 | LeBron *17* replicate 1 | Non-toxic | + |
| 2 | | pExTra-LeBron 17 | LeBron *17* replicate 2 | Non-toxic | + |
| 3 | | pExTra-LeBron 17 | LeBron *17* replicate 3 | Non-toxic | + |

*Key: NG (no growth) - (no pink color) +(faint pink color) ++(obvious pink color) +++ (dark pink color)

#### Gene *18*; Score 0

Images taken after 5 days at 37 ºC

- + 1 2 3 (each plate)

Dilution factor: $10^0$, $10^{-1}$, $10^{-2}$, $10^{-3}$, $10^{-4}$, $10^{-5}$

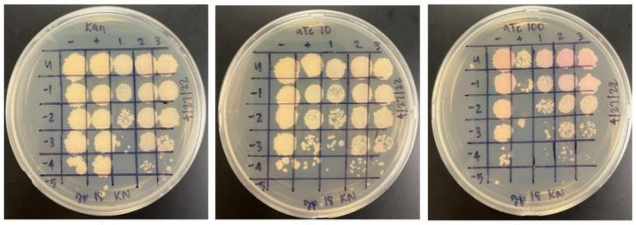

0 ng/ml aTc | 10 ng/ml aTc | 100 ng/ml aTc

| Lane | Gene ID | Plasmid name | Gene name | Toxic/Non-toxic | Colony color on 100 ng/ml aTc plate* |
|---|---|---|---|---|---|
| - Non-toxic control | -- | pExTra03 | Fruitloop 52 mutant | Non-toxic | + |
| + Toxic control | -- | pExTra02 | Fruitloop *52* | Toxic | - |
| 1 | | pExTra-Lebron 18 | Lebron 18 replicate 1 | Non-toxic | ++ |
| 2 | | pExTra-Lebron 18 | Lebron 18 replicate 2 | Non-toxic | ++ |
| 3 | | pExTra-Lebron 18 | Lebron 18 replicate 3 | Non-toxic | ++ |

*Key: NG (no growth) - (no pink color) +(faint pink color) ++(obvious pink color) +++ (dark pink color)

#### Gene *19*; Score *0*

Images taken after 5 days at 37 ºC

- + 1 2 3 (each plate)

Dilution factor: $10^0$, $10^{-1}$, $10^{-2}$, $10^{-3}$, $10^{-4}$, $10^{-5}$

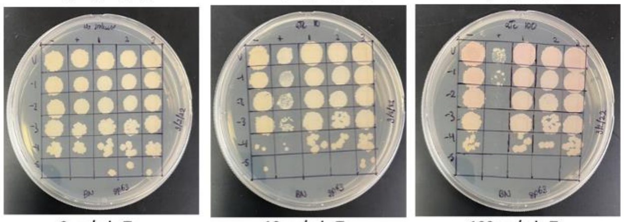

0 ng/ml aTc | 10 ng/ml aTc | 100 ng/ml aTc

| Lane | Gene ID | Plasmid name | Gene name | Toxic/Non-toxic | Colony color on 100 ng/ml aTc plate* |
|---|---|---|---|---|---|
| - Non-toxic control | -- | pExTra03 | Fruitloop *52* mutant | Non-toxic | ++ |
| + Toxic control | -- | pExTra02 | Fruitloop *52* | Toxic | + |
| 1 | | pExTra-Lebron 19 | Lebron *19* replicate 1 | Non-toxic | ++ |
| 2 | | pExTra-Lebron 19 | Lebron *19* replicate 2 | Non-toxic | ++ |
| 3 | | pExTra-Lebron 19 | Lebron *19* replicate 3 | Non-toxic | ++ |

*Key: NG (no growth) - (no pink color) +(faint pink color) ++(obvious pink color) +++ (dark pink color)

#### Gene *20*; Score *1*

Images taken after 5 days at 37 ºC

- + 1 2 3 (each plate)

Dilution factor: $10^0$, $10^{-1}$, $10^{-2}$, $10^{-3}$, $10^{-4}$, $10^{-5}$

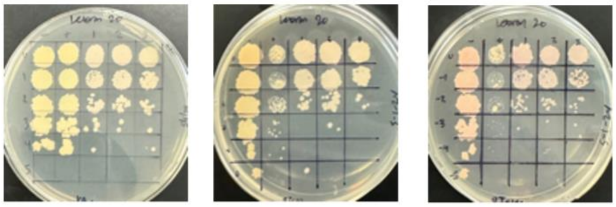

0 ng/ml aTc | 10 ng/ml aTc | 100 ng/ml aTc

| Lane | Gene ID | Plasmid name | Gene name | Toxic/Non-toxic | Colony color on 100 ng/ml aTc plate* |
|---|---|---|---|---|---|
| - Non-toxic control | -- | pExTra03 | Fruitloop *52* mutant | Non-toxic | + |
| + Toxic control | -- | pExTra02 | Fruitloop *52* | Toxic | - |
| 1 | | pExTra-Lebron 20 | Lebron *20* replicate 1 | Toxic | ++ |
| 2 | | pExTra-Lebron 20 | Lebron *20* replicate 2 | Toxic | ++ |
| 3 | | pExTra-Lebron 20 | Lebron *20* replicate 3 | Toxic | ++ |

*Key: NG (no growth) - (no pink color) +(faint pink color) ++(obvious pink color) +++ (dark pink color)

#### Gene *21*; Score *0*

Images taken after 5 days at 37 ºC

- + 1 2 3 (each plate)

Dilution factor: $10^0$, $10^{-1}$, $10^{-2}$, $10^{-3}$, $10^{-4}$, $10^{-5}$

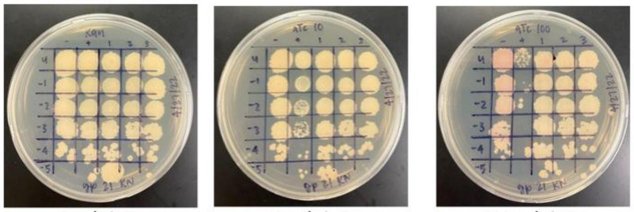

0 ng/ml aTc | 10 ng/ml aTc | 100 ng/ml aTc

| Lane | Gene ID | Plasmid name | Gene name | Toxic/Non-toxic | Colony color on 100 ng/ml aTc plate* |
|---|---|---|---|---|---|
| - Non-toxic control | -- | pExTra03 | Fruitloop *52* mutant | Non-toxic | + |
| + Toxic control | -- | pExTra02 | Fruitloop *52* | Toxic | - |
| 1 | | pExTra-Lebron 21 | Lebron 21 replicate 1 | Non-toxic | - |
| 2 | | pExTra-Lebron 21 | Lebron 21 replicate 2 | Non-toxic | - |
| 3 | | pExTra-Lebron 21 | Lebron 21 replicate 3 | Non-toxic | - |

*Key: NG (no growth) - (no pink color) +(faint pink color) ++(obvious pink color) +++ (dark pink color)

#### Gene *22*; Score *0*

Images taken after 5 days at 37 ºC

- + 1 2 3 (each plate)

Dilution factor: $10^0$, $10^{-1}$, $10^{-2}$, $10^{-3}$, $10^{-4}$, $10^{-5}$

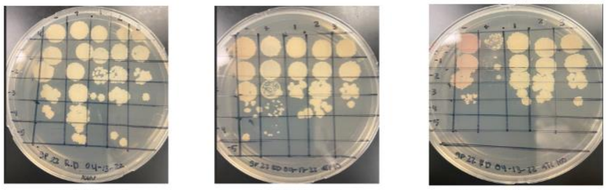

0 ng/ml aTc | 10 ng/ml aTc | 100 ng/ml aTc

| Lane II | Gene ID | Plasmid name | Gene name | Toxic/Non-toxic | Colony color on 100 ng/ml aTc plate* |
|---|---|---|---|---|---|
| - Non-toxic control | -- | pExTra03 | Fruitloop *52 mutant* | Non-toxic | ++ |
| + Toxic control | -- | pExTra02 | Fruitloop *52* | Toxic | + |
| 1 | | pExTra-Lebron 22 | Lebron 22 replicate 1 | Non-toxic | - |
| 2 | | pExTra-Lebron 22 | Lebron 22 replicate 2 | Non-toxic | - |
| 3 | | pExTra-Lebron 22 | Lebron 22 replicate 3 | Non-toxic | - |

*Key: NG (no growth) - (no pink color) +(faint pink color) ++(obvious pink color) +++ (dark pink color)

#### Gene *23*; Score *3*

Images taken after 5 days at 37 ºC

- + 1 2 3 (each plate)

Dilution factor: $10^0$, $10^{-1}$, $10^{-2}$, $10^{-3}$, $10^{-4}$, $10^{-5}$

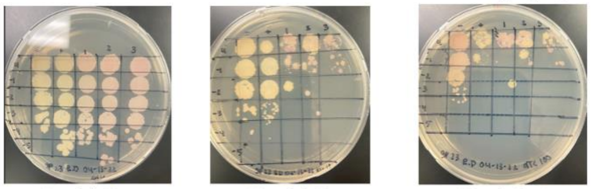

0 ng/ml aTc | 10 ng/ml aTc | 100 ng/ml aTc

| Lane II | Gene ID | Plasmid name | Gene name | Toxic/Non-toxic | Colony color on 100 ng/ml aTc plate* |
|---|---|---|---|---|---|
| - Non-toxic control | -- | pExTra03 | Fruitloop *52 mutant* | Non-toxic | +++ |
| + Toxic control | -- | pExTra02 | Fruitloop *52* | Toxic | + |
| 1 | | pExTra-Lebron 23 | Lebron 23 replicate 1 | Toxic | +++ |
| 2 | | pExTra-Lebron 23 | Lebron 23 replicate 2 | Toxic | + |
| 3 | | pExTra-Lebron 23 | Lebron 23 replicate 3 | Toxic | ++ |

*Key: NG (no growth) - (no pink color) +(faint pink color) ++(obvious pink color) +++ (dark pink color)

#### Gene *24*; Score *0*

Images taken after 5 days at 37 ºC

- + 1 2 3 (each plate)

Dilution factor: $10^0$, $10^{-1}$, $10^{-2}$, $10^{-3}$, $10^{-4}$, $10^{-5}$

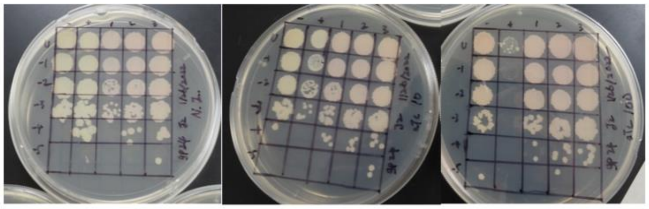

0 ng/ml aTc | 10 ng/ml aTc | 100 ng/ml aTc

| Lane | Gene ID | Plasmid name | Gene name | Toxic/Non-toxic | Colony color on 100 ng/ml aTc plate* |
|---|---|---|---|---|---|
| - Non-toxic control | -- | pExTra03 | Fruitloop 52 mutant | Non-toxic | + |
| + Toxic control | -- | pExTra02 | Fruitloop 52 | Toxic | - |
| 1 | | pExTra-Lebron 24 | Lebron *24* replicate 1 | Non-toxic | + |
| 2 | | pExTra-Lebron 24 | Lebron *24* replicate 2 | Non-toxic | + |
| 3 | | pExTra-Lebron 24 | Lebron *24* replicate 3 | Non-toxic | + |

*Key: NG (no growth) - (no pink color) +(faint pink color) ++(obvious pink color) +++ (dark pink color)

#### Gene *25*; Score *2*

Images taken after 5 days at 37 ºC

Dilution factor: $10^0$, $10^{-1}$, $10^{-2}$, $10^{-3}$, $10^{-4}$, $10^{-5}$

Lanes: - + 1 2 3 | - + 1 2 3 | - + 1 2 3

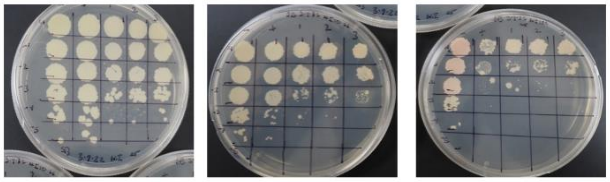

0 ng/ml aTc | 10 ng/ml aTc | 100 ng/ml aTc

| Lane | Gene ID | Plasmid name | Gene name | Toxic/Non-toxic | Colony color on 100 ng/ml aTc plate* |
|---|---|---|---|---|---|
| + Toxic control | -- | pExTra02 | Fruitloop *52* | Non-toxic | + |
| - Non-toxic control | -- | pExTra03 | Fruitloop *52* mutant | Toxic | - |
| 1 | 131436 | pExTra-Lebron 25 | Lebron 25 replicate 1 | Toxic | - |
| 2 | 131436 | pExTra-Lebron 25 | Lebron 25 replicate 2 | Toxic | - |
| 3 | 131436 | pExTra-Lebron 25 | Lebron 25 replicate 3 | Toxic | - |

*Key: NG (no growth) - (no pink color) +(faint pink color) ++(obvious pink color) +++ (dark pink color)

#### Gene *26*; Score *0*

Images taken after 5 days at 37 ºC

Dilution factor: $10^0$, $10^{-1}$, $10^{-2}$, $10^{-3}$, $10^{-4}$, $10^{-5}$

Lanes: - + 1 2 3 | - + 1 2 3 | - + 1 2 3

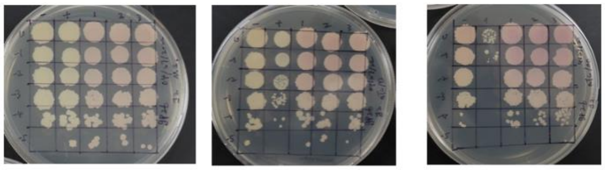

0 ng/ml aTc | 10 ng/ml aTc | 100 ng/ml aTc

| Lane | Gene ID | Plasmid name | Gene name | Toxic/Non-toxic | Colony color on 100 ng/ml aTc plate* |
|---|---|---|---|---|---|
| - Non-toxic control | -- | pExTra03 | Fruitloop *52* mutant | Non-toxic | + |
| + Toxic control | -- | pExTra02 | Fruitloop *52* | Toxic | - |
| 1 | | pExTra-Lebron 26 | Lebron *26* replicate 1 | Non-toxic | ++ |
| 2 | | pExTra-Lebron 26 | Lebron *26* replicate 2 | Non-toxic | ++ |
| 3 | | pExTra-Lebron 26 | Lebron *26* replicate 3 | Non-toxic | ++ |

*Key: NG (no growth) - (no pink color) +(faint pink color) ++(obvious pink color) +++ (dark pink color)

#### Gene *27*; Score *0*

Images taken after 5 days at 37 ºC

Dilution factor: $10^0$, $10^{-1}$, $10^{-2}$, $10^{-3}$, $10^{-4}$, $10^{-5}$

Lanes: - + 1 2 3 | - + 1 2 3 | - + 1 2 3

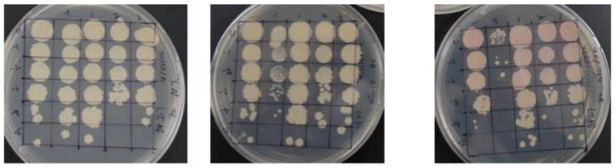

0 ng/ml aTc | 10 ng/ml aTc | 100 ng/ml aTc

| Lane | Gene ID | Plasmid name | Gene name | Toxic/Non-toxic | Colony color on 100 ng/ml aTc plate* |
|---|---|---|---|---|---|
| - Non-toxic control | -- | pExTra03 | Fruitloop *52* mutant | Non-toxic | + |
| + Toxic control | -- | pExTra02 | Fruitloop *52* | Toxic | - |
| 1 | | pExTra-Lebron 27 | Lebron *27* replicate 1 | Non-toxic | + |
| 2 | | pExTra-Lebron 27 | Lebron *27* replicate 2 | Non-toxic | + |
| 3 | | pExTra-Lebron 27 | Lebron *27* replicate 3 | Non-toxic | + |

*Key: NG (no growth) - (no pink color) +(faint pink color) ++(obvious pink color) +++ (dark pink color)

#### Gene *28*; Score *0*

Images taken after 5 days at 37 ºC

Dilution factor: $10^0$, $10^{-1}$, $10^{-2}$, $10^{-3}$, $10^{-4}$, $10^{-5}$

Lanes: - + 1 2 3 | - + 1 2 3 | - + 1 2 3

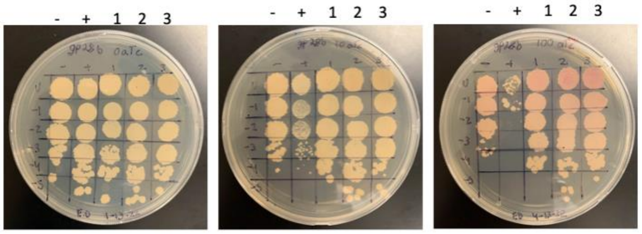

0 ng/ml aTc | 10 ng/ml aTc | 100 ng/ml aTc

| Lane | Gene ID | Plasmid name | Gene name | Toxic/Non-toxic | Colony color on 100 ng/ml aTc plate* |
|---|---|---|---|---|---|
| + Toxic control | -- | pExTra02 | Fruitloop *52* | Toxic | - |
| - Non-toxic control | -- | pExTra03 | Fruitloop *52* mutant | Non-toxic | + |
| 1 | | pExTra-Lebron 28 | Lebron *28* replicate 1 | Non-toxic | ++ |
| 2 | | pExTra-Lebron 28 | Lebron *28* replicate 2 | Non-toxic | ++ |
| 3 | | pExTra-Lebron 28 | Lebron *28* replicate 3 | Non-toxic | ++ |

*Key: NG (no growth) - (no pink color) +(faint pink color) ++(obvious pink color) +++ (dark pink color)

#### Gene *29*; Score *2*

Images taken after 5 days at 37 ºC

Dilution factor: $10^0$, $10^{-1}$, $10^{-2}$, $10^{-3}$, $10^{-4}$, $10^{-5}$

Lanes: - + 1 2 3 | - + 1 2 3 | - + 1 2 3

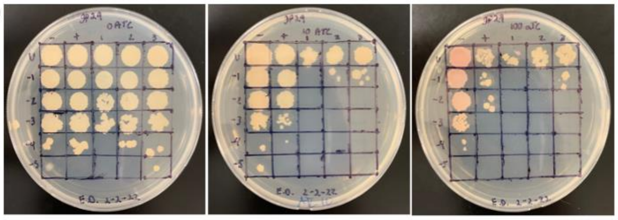

0 ng/ml aTc | 10 ng/ml aTc | 100 ng/ml aTc

| Lane | Gene ID | Plasmid name | Gene name | Toxic/Non-toxic | Colony color on 100 ng/ml aTc plate* |
|---|---|---|---|---|---|
| + Toxic control | -- | pExTra02 | Fruitloop *52* | Toxic | - |
| - Non-toxic control | -- | pExTra03 | Fruitloop *52* mutant | Non-toxic | ++ |
| 1 | | pExTra-LeBron 29 | LeBeon *29* replicate 1 | Toxic | + |
| 2 | | pExTra-LeBron 29 | LeBron *29* replicate 2 | Toxic | - |
| 3 | | pExTra-LeBron 29 | LeBron *29* replicate 3 | Toxic | + |

*Key: NG (no growth) - (no pink color) +(faint pink color) ++(obvious pink color) +++ (dark pink color)

#### Gene *30*; Score *0*

Images taken after 5 days at 37 ºC

Dilution factor: $10^0$, $10^{-1}$, $10^{-2}$, $10^{-3}$, $10^{-4}$, $10^{-5}$

Lanes: - + 1 2 3 | - + 1 2 3 | - + 1 2 3

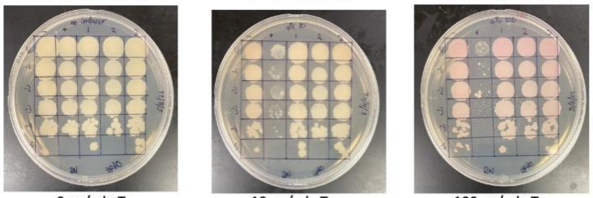

0 ng/ml aTc | 10 ng/ml aTc | 100 ng/ml aTc

| Lane | Gene ID | Plasmid name | Gene name | Toxic/Non-toxic | Colony color on 100 ng/ml aTc plate* |
|---|---|---|---|---|---|
| + Toxic control | -- | pExTra02 | Fruitloop *52* | Toxic | + |
| - Non-toxic control | -- | pExTra03 | Fruitloop *52* mutant | Non-toxic | + |
| 1 | | pExTra-Lebron 30 | Lebron *30* replicate 1 | Non-toxic | ++ |
| 2 | | pExTra-Lebron 30 | Lebron *30* replicate 2 | Non-toxic | ++ |
| 3 | | pExTra-Lebron 30 | Lebron *30* replicate 3 | Non-toxic | ++ |

*Key: NG (no growth) - (no pink color) +(faint pink color) ++(obvious pink color) +++ (dark pink color)

#### Gene *31*; Score *0*

Images taken after 5 days at 37 ºC

Dilution factor: $10^0$, $10^{-1}$, $10^{-2}$, $10^{-3}$, $10^{-4}$, $10^{-5}$

Lanes: - + 1 2 3 | - + 1 2 3 | - + 1 2 3

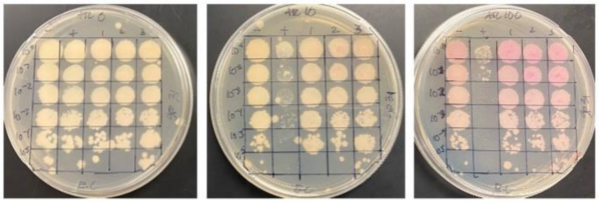

0 ng/ml aTc | 10 ng/ml aTc | 100 ng/ml aTc

| Lane | Gene ID | Plasmid name | Gene name | Toxic/Non-toxic | Colony color on 100 ng/ml aTc plate* |
|---|---|---|---|---|---|
| + Toxic control | -- | pExTra02 | Fruitloop *52* | Toxic | - |
| - Non-toxic control | -- | pExTra03 | Fruitloop *52* mutant | Non-toxic | + |
| 1 | | pExTra-Lebron 31 | Lebron 31 replicate 1 | Non-toxic | ++ |
| 2 | | pExTra-Lebron 31 | Lebron 31 replicate 2 | Non-toxic | ++ |
| 3 | | pExTra-Lebron 31 | Lebron 31 replicate 3 | Non-toxic | ++ |

*Key: NG (no growth) - (no pink color) +(faint pink color) ++(obvious pink color) +++ (dark pink color)

#### Gene *32*; Score *0*

Images taken after 5 days at 37 ºC

Dilution factor: $10^0$, $10^{-1}$, $10^{-2}$, $10^{-3}$, $10^{-4}$, $10^{-5}$

Lanes: - + 1 2 3 | - + 1 2 3 | - + 1 2 3

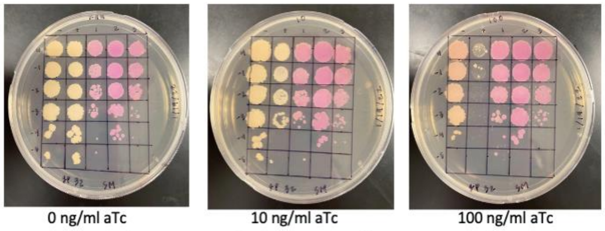

0 ng/ml aTc | 10 ng/ml aTc | 100 ng/ml aTc

| Lane | Gene ID | Plasmid name | Gene name | Toxic/Non-toxic | Colony color on 100 ng/ml aTc plate* |
|---|---|---|---|---|---|
| + Toxic control | -- | pExTra02 | Fruitloop *52* | Toxic | - |
| - Non-toxic control | -- | pExTra03 | Fruitloop *52* mutant | Non-toxic | + |
| 1 | | pExTra-Lebron 32 | Lebron *32* replicate 1 | Non-toxic | +++ |
| 2 | | pExTra-Lebron 32 | Lebron *32* replicate 2 | Non-toxic | +++ |
| 3 | | pExTra-Lebron 32 | Lebron *32* replicate 3 | Non-toxic | +++ |

*Key: NG (no growth) - (no pink color) +(faint pink color) ++(obvious pink color) +++ (dark pink color)

#### Gene *33*; Score *0*

Images taken after 5 days at 37 ºC

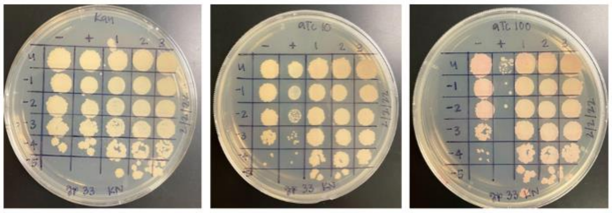


| Lane | Gene ID | Plasmid name | Gene name | Toxic/Non-toxic | Colony color on 100 ng/ml aTc plate* |
|---|---|---|---|---|---|
| - Non-toxic control | -- | pExTra03 | Fruitloop *52* mutant | Non-toxic | + |
| + Toxic control | -- | pExTra02 | Fruitloop *52* | Toxic | - |
| 1 | | pExTra-Lebron 33 | Lebron 33 replicate 1 | Non-toxic | + |
| 2 | | pExTra-Lebron 33 | Lebron 33 replicate 2 | Non-toxic | + |
| 3 | | pExTra-Lebron 33 | Lebron 33 replicate 3 | Non-toxic | + |

*Key: NG (no growth) - (no pink color) +(faint pink color) ++(obvious pink color) +++ (dark pink color)

#### Gene *34*; Score *0*

Images taken after 5 days at 37 ºC

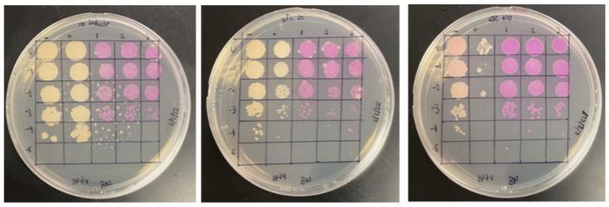


| Lane | Gene ID | Plasmid name | Gene name | Toxic/Non-toxic | Colony color on 100 ng/ml aTc plate* |
|---|---|---|---|---|---|
| - Non-toxic control | -- | pExTra03 | Fruitloop *52* mutant | Non-toxic | + |
| + Toxic control | -- | pExTra02 | Fruitloop *52* | Toxic | - |
| 1 | | pExTra-Lebron 34 | Lebron *34* replicate 1 | Non-toxic | +++ |
| 2 | | pExTra-Lebron 34 | Lebron *34* replicate 2 | Non-toxic | +++ |
| 3 | | pExTra-Lebron 34 | Lebron *34* replicate 3 | Non-toxic | +++ |

*Key: NG (no growth) - (no pink color) +(faint pink color) ++(obvious pink color) +++ (dark pink color)

#### Gene *35*; Score *3*

Images taken after 5 days at 37 ºC

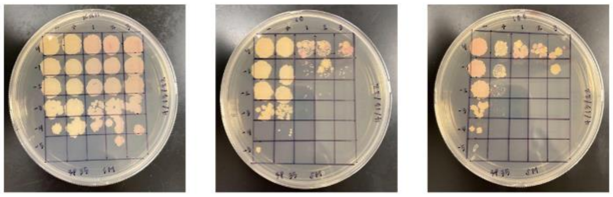


| Lane | Gene ID | Plasmid name | Gene name | Toxic/Non-toxic | Colony color on 100 ng/ml aTc plate* |
|---|---|---|---|---|---|
| + Toxic control | -- | pExTra02 | Fruitloop *52* | Toxic | - |
| - Non-toxic control | -- | pExTra03 | Fruitloop *52* mutant | Non-toxic | + |
| 1 | | pExTra-Lebron35 | Lebron *35* replicate 1 | Toxic | + |
| 2 | | pExTra-Lebron35 | Lebron *35* replicate 2 | Toxic | + |
| 3 | | pExTra-Lebron35 | Lebron *35* replicate 3 | Toxic | + |

*Key: NG (no growth) - (no pink color) +(faint pink color) ++(obvious pink color) +++ (dark pink color)

#### Gene *36*; Score *1*

Images taken after 5 days at 37 ºC

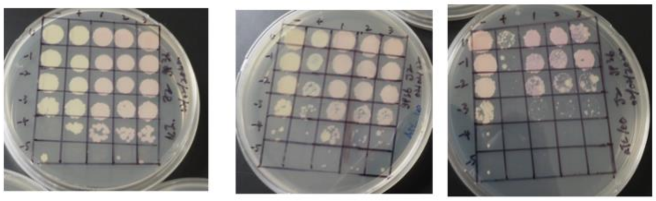


| Lane | Gene ID | Plasmid name | Gene name | Toxic/Non-toxic | Colony color on 100 ng/ml aTc plate* |
|---|---|---|---|---|---|
| - Non-toxic control | -- | pExTra03 | Fruitloop *52* mutant | Non-toxic | + |
| + Toxic control | -- | pExTra02 | Fruitloop *52* | Toxic | - |
| 1 | | pExTra-Lebron 36 | Lebron *36* replicate 1 | Toxic | ++ |
| 2 | | pExTra-Lebron 36 | Lebron *36* replicate 2 | Toxic | ++ |
| 3 | | pExTra-Lebron 36 | Lebron *36* replicate 3 | Toxic | ++ |

*Key: NG (no growth) - (no pink color) +(faint pink color) ++(obvious pink color) +++ (dark pink color)

#### Gene *37*; Score *0*

Images taken after 5 days at 37 ºC

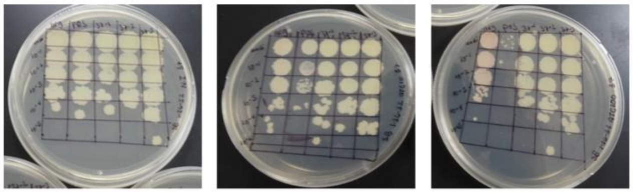


| Lane | Gene ID | Plasmid name | Gene name | Toxic/Non-toxic | Colony color on 100 ng/ml aTc plate* |
|---|---|---|---|---|---|
| + Toxic control | -- | pExTra02 | Fruitloop *52* | Non-toxic | + |
| - Non-toxic control | -- | pExTra03 | Fruitloop *52* mutant | Toxic | - |
| 1 | | pExTra-Lebron 37 | Lebron 37 replicate 1 | Non-toxic | - |
| 2 | | pExTra-Lebron 37 | Lebron 37 replicate 2 | Non-toxic | - |
| 3 | | pExTra-Lebron 37 | Lebron 37 replicate 3 | Non-toxic | - |

*Key: NG (no growth) - (no pink color) +(faint pink color) ++(obvious pink color) +++ (dark pink color)

#### Gene *38*; Score *0*

Images taken after 5 days at 37 ºC

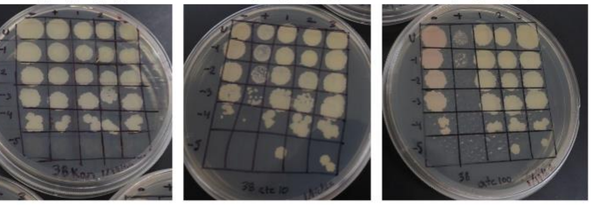


| Lane | Gene ID | Plasmid name | Gene name | Toxic/Non-toxic | Colony color on 100 ng/ml aTc plate* |
|---|---|---|---|---|---|
| - Non-toxic control | -- | pExTra03 | Fruitloop *52* mutant | Non-toxic | + |
| + Toxic control | -- | pExTra02 | Fruitloop *52* | Toxic | - |
| 1 | | pExTra-Lebron38 | Lebron *38* replicate 1 | Non-toxic | - |
| 2 | | pExTra-Lebron 38 | Lebron *38* replicate 2 | Non-toxic | - |
| 3 | | pExTra-Lebron 38 | Lebron *38* replicate 3 | Non-toxic | - |

*Key: NG (no growth) - (no pink color) +(faint pink color) ++(obvious pink color) +++ (dark pink color)

#### Gene *39*; Score *0*

Images taken after 5 days at 37 ºC

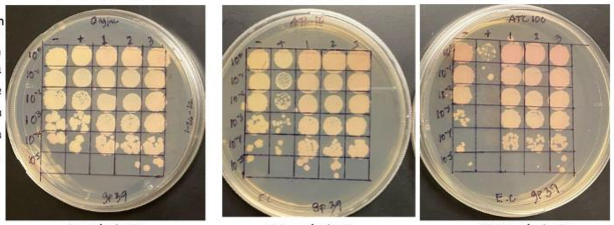


| Lane | Gene ID | Plasmid name | Gene name | Toxic/Non-toxic | Colony color on 100 ng/ml aTc plate* |
|---|---|---|---|---|---|
| + Toxic control | -- | pExTra02 | Fruitloop *52* | Toxic | - |
| - Non-toxic control | -- | pExTra03 | Fruitloop *52* mutant | Non-toxic | + |
| 1 | | pExTra-Lebron 39 | Lebron *39* replicate 1 | Non-toxic | ++ |
| 2 | | pExTra-Lebron 39 | Lebron *39* replicate 2 | Non-toxic | ++ |
| 3 | | pExTra-Lebron 39 | Lebron *39* replicate 3 | Non-toxic | ++ |

*Key: NG (no growth) - (no pink color) +(faint pink color) ++(obvious pink color) +++ (dark pink color)

#### Gene *40*; Score *0*

Images taken after 5 days at 37 ºC

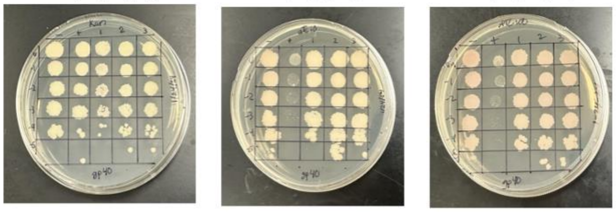


| Lane | Gene ID | Plasmid name | Gene name | Toxic/Non-toxic | Colony color on 100 ng/ml aTc plate* |
|---|---|---|---|---|---|
| - Non-toxic control | -- | pExTra03 | Fruitloop *52* mutant | Non-toxic | ++ |
| + Toxic control | -- | pExTra02 | Fruitloop *52* | Toxic | - |
| 1 | | pExTra-Lebron 40 | Lebron *40* replicate 1 | Non-toxic | + |
| 2 | | pExTra-Lebron 40 | Lebron *40* replicate 2 | Non-toxic | + |
| 3 | | pExTra-Lebron 40 | Lebron *40* replicate 3 | Non-toxic | + |

*Key: NG (no growth) - (no pink color) +(faint pink color) ++(obvious pink color) +++ (dark pink color)

Images taken after 5 days at 37 ºC

#### Gene *41*; Score *0*

- + 1 2 3    - + 1 2 3    - + 1 2 3

Dilution factor: $10^0$, $10^{-1}$, $10^{-2}$, $10^{-3}$, $10^{-4}$, $10^{-5}$

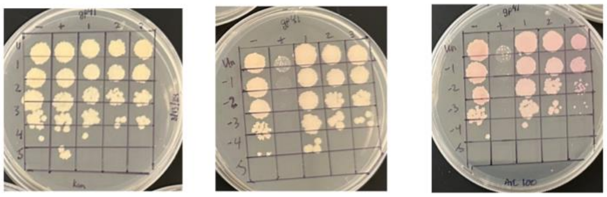

0 ng/ml aTc    10 ng/ml aTc    100 ng/ml aTc

| Lane | Gene ID | Plasmid name | Gene name | Toxic/Non-toxic | Colony color on 100 ng/ml aTc plate* |
|---|---|---|---|---|---|
| - Non-toxic control | -- | pExTra03 | Fruitloop *52* mutant | Non-toxic | ++ |
| + Toxic control | -- | pExTra02 | Fruitloop *52* | Toxic | - |
| 1 | | pExTra-Lebron 41 | Lebron *41* replicate 1 | Non-toxic | ++ |
| 2 | | pExTra-Lebron 41 | Lebron *41* replicate 2 | Non-toxic | ++ |
| 3 | | pExTra-Lebron 41 | Lebron *41* replicate 3 | Non-toxic | ++ |

*Key: NG (no growth) - (no pink color) +(faint pink color) ++(obvious pink color) +++ (dark pink color)

Images taken after 5 days at 37 ºC

#### Gene *42*; Score *0*

- + 1 2 3    - + 1 2 3    - + 1 2 3

Dilution factor: $10^0$, $10^{-1}$, $10^{-2}$, $10^{-3}$, $10^{-4}$, $10^{-5}$

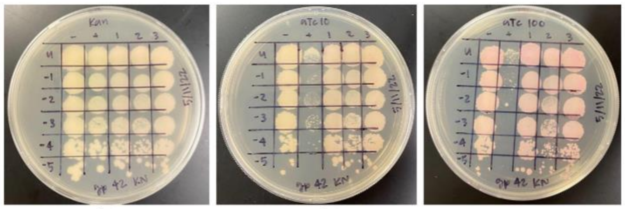

0 ng/ml aTc    10 ng/ml aTc    100 ng/ml aTc

| Lane | Gene ID | Plasmid name | Gene name | Toxic/Non-toxic | Colony color on 100 ng/ml aTc plate* |
|---|---|---|---|---|---|
| - Non-toxic control | -- | pExTra03 | Fruitloop *52* mutant | Non-toxic | + |
| + Toxic control | -- | pExTra02 | Fruitloop *52* | Toxic | - |
| 1 | | pExTra-Lebron 42 | Lebron 42 replicate 1 | Non-toxic | ++ |
| 2 | | pExTra-Lebron 42 | Lebron 42 replicate 2 | Non-toxic | ++ |
| 3 | | pExTra-Lebron 42 | Lebron 42 replicate 3 | Non-toxic | ++ |

*Key: NG (no growth) - (no pink color) +(faint pink color) ++(obvious pink color) +++ (dark pink color)

Images taken after 5 days at 37 ºC

#### Gene *43*; Score *0*

- + 1 2 3    - + 1 2 3    - + 1 2 3

Dilution factor: $10^0$, $10^{-1}$, $10^{-2}$, $10^{-3}$, $10^{-4}$, $10^{-5}$

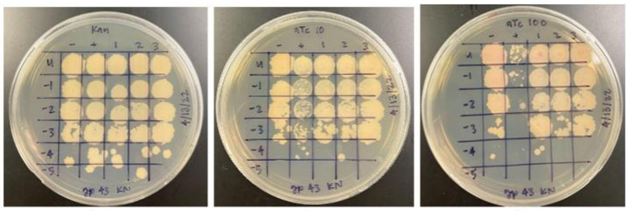

0 ng/ml aTc    10 ng/ml aTc    100 ng/ml aTc

| Lane | Gene ID | Plasmid name | Gene name | Toxic/Non-toxic | Colony color on 100 ng/ml aTc plate* |
|---|---|---|---|---|---|
| - Non-toxic control | -- | pExTra03 | Fruitloop *52* mutant | Non-toxic | + |
| + Toxic control | -- | pExTra02 | Fruitloop *52* | Toxic | - |
| 1 | | pExTra-Lebron 43 | Lebron 43 replicate 1 | Non-toxic | + |
| 2 | | pExTra-Lebron 43 | Lebron 43 replicate 2 | Non-toxic | + |
| 3 | | pExTra-Lebron 43 | Lebron 43 replicate 3 | Non-toxic | + |

*Key: NG (no growth) - (no pink color) +(faint pink color) ++(obvious pink color) +++ (dark pink color)

Images taken after 5 days at 37 ºC

#### Gene *44*; Score *0*

- + 1 2 3    - + 1 2 3    - + 1 2 3

Dilution factor: $10^0$, $10^{-1}$, $10^{-2}$, $10^{-3}$, $10^{-4}$, $10^{-5}$

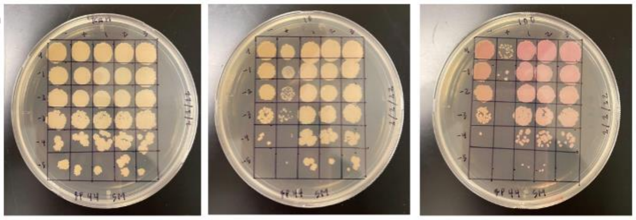

0 ng/ml aTc    10 ng/ml aTc    100 ng/ml aTc

| Lane | Gene ID | Plasmid name | Gene name | Toxic/Non-toxic | Colony color on 100 ng/ml aTc plate* |
|---|---|---|---|---|---|
| + Toxic control | -- | pExTra02 | Fruitloop *52* | Toxic | - |
| - Non-toxic control | -- | pExTra03 | Fruitloop *52* mutant | Non-toxic | + |
| 1 | | pExTra-Lebron 44 | Lebron *44* replicate 1 | Non-toxic | ++ |
| 2 | | pExTra-Lebron 44 | Lebron *44* replicate 2 | Non-toxic | ++ |
| 3 | | pExTra-Lebron 44 | Lebron *44* replicate 3 | Non-toxic | ++ |

*Key: NG (no growth) - (no pink color) +(faint pink color) ++(obvious pink color) +++ (dark pink color)

Images taken after 5 days at 37 ºC

#### Gene *45*; Score *0*

- + 1 2 3    - + 1 2 3    - + 1 2 3

Dilution factor: $10^0$, $10^{-1}$, $10^{-2}$, $10^{-3}$, $10^{-4}$, $10^{-5}$

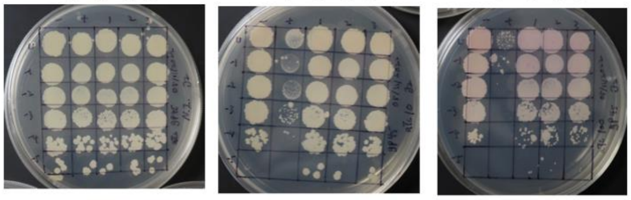

0 ng/ml aTc    10 ng/ml aTc    100 ng/ml aTc

| Lane | Gene ID | Plasmid name | Gene name | Toxic/Non-toxic | Colony color on 100 ng/ml aTc plate* |
|---|---|---|---|---|---|
| - Non-toxic control | -- | pExTra03 | Fruitloop *52* mutant | Non-toxic | + |
| + Toxic control | -- | pExTra02 | Fruitloop *52* | Non-Toxic | - |
| 1 | | pExTra-Lebron 45 | Lebron *45* replicate 1 | Non-Toxic | ++ |
| 2 | | pExTra-Lebron 45 | Lebron *45* replicate 2 | Non-Toxic | ++ |
| 3 | | pExTra-Lebron 45 | Lebron *45* replicate 3 | Non-Toxic | ++ |

*Key: NG (no growth) - (no pink color) +(faint pink color) ++(obvious pink color) +++ (dark pink color)

Images taken after 5 days at 37 ºC

#### Gene *47*; Score *2*

- + 1 2 3    - + 1 2 3    - + 1 2 3

Dilution factor: $10^0$, $10^{-1}$, $10^{-2}$, $10^{-3}$, $10^{-4}$, $10^{-5}$

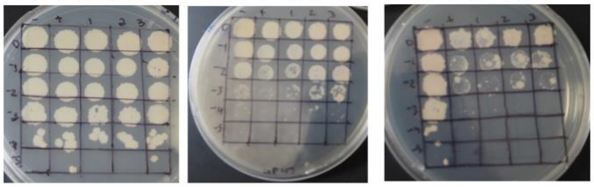

0 ng/ml aTc    10 ng/ml aTc    100 ng/ml aTc

| Lane | Gene ID | Plasmid name | Gene name | Toxic/Non-toxic | Colony color on 100 ng/ml aTc plate* |
|---|---|---|---|---|---|
| + Toxic control | -- | pExTra02 | Fruitloop *52* | Toxic | + |
| - Non-toxic control | -- | pExTra03 | Fruitloop *52* mutant | Non-toxic | - |
| 1 | | pExTra-Lebron 47 | Lebron *47* replicate 1 | Toxic | + |
| 2 | | pExTra-Lebron 47 | Lebron *47* replicate 2 | Toxic | + |
| 3 | | pExTra-Lebron 47 | Lebron *47* replicate 3 | Toxic | + |

*Key: NG (no growth) - (no pink color) +(faint pink color) ++(obvious pink color) +++ (dark pink color)

Images taken after 5 days at 37 ºC

#### Gene *48*; Score *0*

- + 1 2 3    - + 1 2 3    - + 1 2 3

Dilution factor: $10^0$, $10^{-1}$, $10^{-2}$, $10^{-3}$, $10^{-4}$, $10^{-5}$

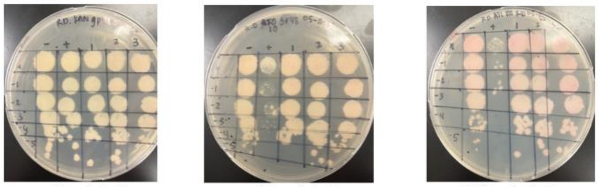

0 ng/ml aTc    10 ng/ml aTc    100 ng/ml aTc

| Lane ll | Gene ID | Plasmid name | Gene name | Toxic/Non-toxic | Colony color on 100 ng/ml aTc plate* |
|---|---|---|---|---|---|
| - Non-toxic control | -- | pExTra03 | Fruitloop *52 mutant* | Non-toxic | +++ |
| + Toxic control | -- | pExTra02 | Fruitloop *52* | Toxic | + |
| 1 | | pExTra-Lebron 48 | Lebron 48 replicate 1 | Non-toxic | ++ |
| 2 | | pExTra-Lebron 48 | Lebron 48 replicate 2 | Non-toxic | ++ |
| 3 | | pExTra-Lebron 48 | Lebron 48 replicate 3 | Non-toxic | ++ |

*Key: NG (no growth) - (no pink color) +(faint pink color) ++(obvious pink color) +++ (dark pink color)

Images taken after 5 days at 37 ºC

#### Gene *49*; Score *1*

- + 1 2 3    - + 1 2 3    - + 1 2 3

Dilution factor: $10^0$, $10^{-1}$, $10^{-2}$, $10^{-3}$, $10^{-4}$, $10^{-5}$

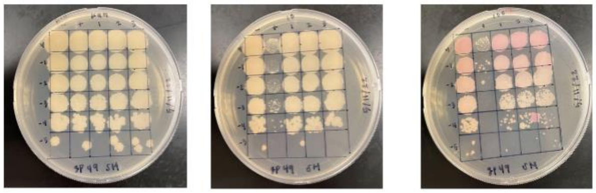

0 ng/ml aTc    10 ng/ml aTc    100 ng/ml aTc

| Lane | Gene ID | Plasmid name | Gene name | Toxic/Non-toxic | Colony color on 100 ng/ml aTc plate* |
|---|---|---|---|---|---|
| + Toxic control | -- | pExTra02 | Fruitloop *52* | Toxic | - |
| - Non-toxic control | -- | pExTra03 | Fruitloop *52* mutant | Non-toxic | ++ |
| 1 | | pExTra-Lebron 49 | Lebron *49* replicate 1 | Toxic | ++ |
| 2 | | pExTra-Lebron 49 | Lebron *49* replicate 2 | Toxic | ++ |
| 3 | | pExTra-Lebron 49 | Lebron *49* replicate 3 | Toxic | ++ |

*Key: NG (no growth) - (no pink color) +(faint pink color) ++(obvious pink color) +++ (dark pink color)

#### Gene *50*; Score *2*

Images taken after 5 days at 37 ºC

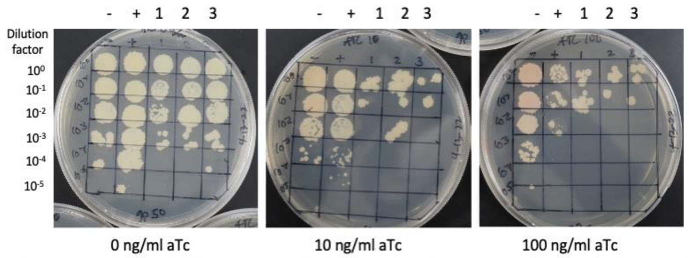


| Lane | Gene ID | Plasmid name | Gene name | Toxic/Non-toxic | Colony color on 100 ng/ml aTc plate* |
|---|---|---|---|---|---|
| + Toxic control | -- | pExTra02 | Fruitloop *52* | Toxic | - |
| - Non-toxic control | -- | pExTra03 | Fruitloop *52* mutant | Non-toxic | ++ |
| 1 | | pExTra-Lebron 50 | Lebron 50 replicate 1 | toxic | + |
| 2 | | pExTra-Lebron 50 | Lebron 50 replicate 2 | toxic | + |
| 3 | | pExTra-Lebron 50 | Lebron 50 replicate 3 | toxic | + |

*Key: NG (no growth) - (no pink color) +(faint pink color) ++(obvious pink color) +++ (dark pink color)

#### Gene *51*; Score *0*

Images taken after 5 days at 37 ºC

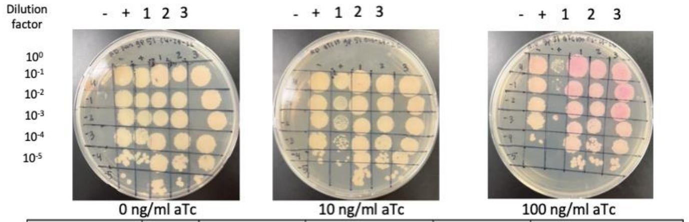


| Lane | Gene ID | Plasmid name | Gene name | Toxic/Non-toxic | Colony color on 100 ng/ml aTc plate* |
|---|---|---|---|---|---|
| - Non-toxic control | -- | pExTra03 | Fruitloop *52 mutant* | Non-toxic | ++ |
| + Toxic control | -- | pExTra02 | Fruitloop *52* | Toxic | + |
| 1 | | pExTra-Lebron 51 | Lebron 51 replicate 1 | Non-Toxic | +++ |
| 2 | | pExTra-Lebron 51 | Lebron 51 replicate 2 | Non-Toxic | +++ |
| 3 | | pExTra-Lebron 51 | Lebron 51 replicate 3 | Non-Toxic | +++ |

*Key: NG (no growth) - (no pink color) +(faint pink color) ++(obvious pink color) +++ (dark pink color)

#### Gene *52*; Score *0*

Images taken after 5 days at 37 ºC

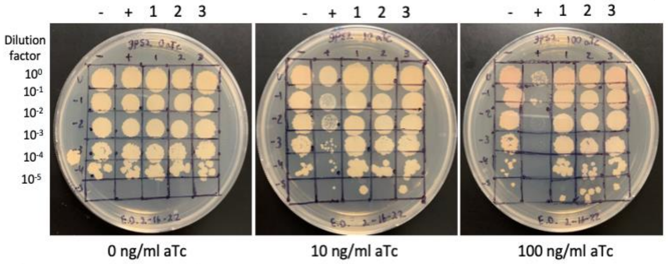


| Lane | Gene ID | Plasmid name | Gene name | Toxic/Non-toxic | Colony color on 100 ng/ml aTc plate* |
|---|---|---|---|---|---|
| + Toxic control | -- | pExTra02 | Fruitloop *52* | Toxic | - |
| - Non-toxic control | -- | pExTra03 | Fruitloop *52* mutant | Non-toxic | ++ |
| 1 | | pExTra-Lebron 52 | Lebron *52* replicate 1 | Non-toxic | ++ |
| 2 | | pExTra-Lebron 52 | Lebron *52* replicate 2 | Non-toxic | ++ |
| 3 | | pExTra-Lebron 52 | Lebron *52*replicate 3 | Non-toxic | ++ |

*Key: NG (no growth) - (no pink color) +(faint pink color) ++(obvious pink color) +++ (dark pink color)

#### Gene *53*; Score *2*

Images taken after 5 days at 37 ºC

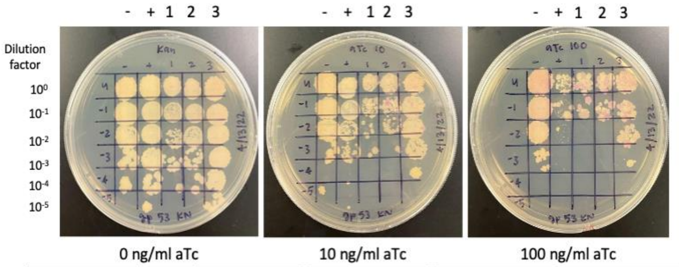


| Lane | Gene ID | Plasmid name | Gene name | Toxic/Non-toxic | Colony color on 100 ng/ml aTc plate* |
|---|---|---|---|---|---|
| - Non-toxic control | -- | pExTra03 | Fruitloop *52* mutant | Non-toxic | + |
| + Toxic control | -- | pExTra02 | Fruitloop *52* | Toxic | - |
| 1 | | pExTra-Lebron 53 | Lebron 53 replicate 1 | Toxic | + |
| 2 | | pExTra-Lebron 53 | Lebron 53 replicate 2 | Toxic | + |
| 3 | | pExTra-Lebron 53 | Lebron 53 replicate 3 | Toxic | + |

*Key: NG (no growth) - (no pink color) +(faint pink color) ++(obvious pink color) +++ (dark pink color)

#### Gene *54*; Score *0*

Images taken after 5 days at 37 ºC

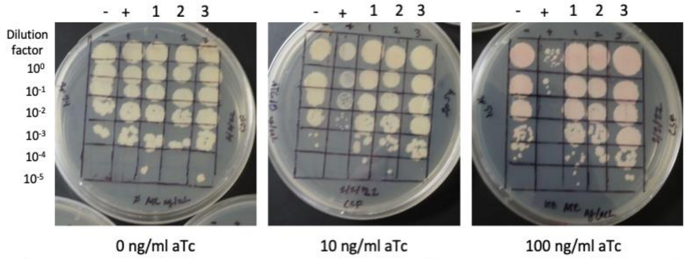


| Lane | Gene ID | Plasmid name | Gene name | Toxic/Non-toxic | Colony color on 100 ng/ml aTc plate* |
|---|---|---|---|---|---|
| - Non-toxic control | -- | pExTra03 | Fruitloop *52* mutant | Non-toxic | + |
| + Toxic control | -- | pExTra02 | Fruitloop *52* | Toxic | - |
| 1 | | pExTra-Lebron 54 | Lebron *54* replicate 1 | Non-toxic | ++ |
| 2 | | pExTra-Lebron 54 | Lebron *54* replicate 2 | Non-toxic | ++ |
| 3 | | pExTra-Lebron 54 | Lebron *54* replicate 3 | Non-toxic | ++ |

*Key: NG (no growth) - (no pink color) +(faint pink color) ++(obvious pink color) +++ (dark pink color)

#### Gene *55*; Score *0*

Images taken after 5 days at 37 ºC

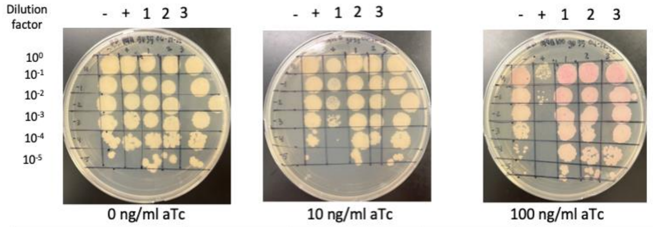


| Lane II | Gene ID | Plasmid name | Gene name | Toxic/Non-toxic | Colony color on 100 ng/ml aTc plate* |
|---|---|---|---|---|---|
| - Non-toxic control | -- | pExTra03 | Fruitloop *52 mutant* | Non-toxic | ++ |
| + Toxic control | -- | pExTra02 | Fruitloop *52* | Toxic | + |
| 1 | | pExTra-Lebron 55 | Lebron 55 replicate 1 | Non-Toxic | +++ |
| 2 | | pExTra-Lebron 55 | Lebron 55 replicate 2 | Non-Toxic | +++ |
| 3 | | pExTra-Lebron 55 | Lebron 55 replicate 3 | Non-Toxic | +++ |

*Key: NG (no growth) - (no pink color) +(faint pink color) ++(obvious pink color) +++ (dark pink color)

#### Gene *56*; Score *0*

Images taken after 5 days at 37 ºC

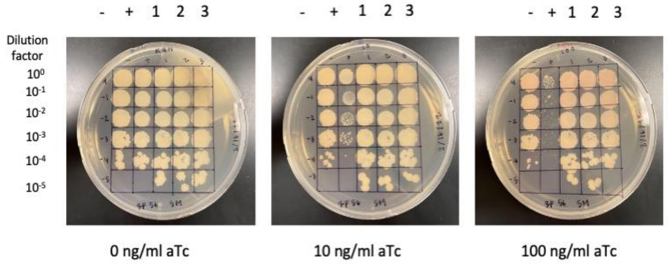


| Lane | Gene ID | Plasmid name | Gene name | Toxic/Non-toxic | Colony color on 100 ng/ml aTc plate* |
|---|---|---|---|---|---|
| + Toxic control | -- | pExTra02 | Fruitloop *52* | Toxic | - |
| - Non-toxic control | -- | pExTra03 | Fruitloop *52* mutant | Non-toxic | + |
| 1 | | pExTra-Lebron 56 | Lebron *56* replicate 1 | Non-toxic | + |
| 2 | | pExTra-Lebron 56 | Lebron *56* replicate 2 | Non-toxic | + |
| 3 | | pExTra-Lebron 56 | Lebron 56 replicate 3 | Non-toxic | + |

*Key: NG (no growth) - (no pink color) +(faint pink color) ++(obvious pink color) +++ (dark pink color)

#### Gene *57*; Score *0*

Images taken after 5 days at 37 ºC

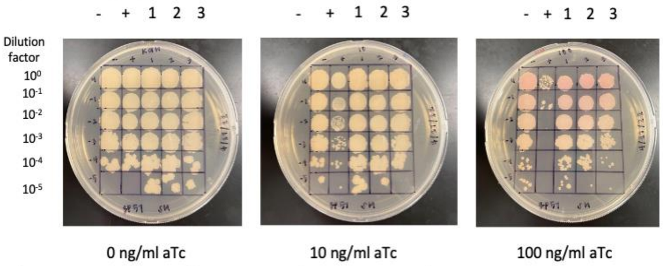


| Lane | Gene ID | Plasmid name | Gene name | Toxic/Non-toxic | Colony color on 100 ng/ml aTc plate* |
|---|---|---|---|---|---|
| + Toxic control | -- | pExTra02 | Fruitloop *52* | Toxic | - |
| - Non-toxic control | -- | pExTra03 | Fruitloop *52* mutant | Non-toxic | ++ |
| 1 | | pExTra-Lebron 57 | Lebron *57* replicate 1 | Non-toxic | ++ |
| 2 | | pExTra-Lebron 57 | Lebron *57* replicate 2 | Non-toxic | ++ |
| 3 | | pExTra-Lebron 57 | Lebron *57* replicate 3 | Non-toxic | ++ |

*Key: NG (no growth) - (no pink color) +(faint pink color) ++(obvious pink color) +++ (dark pink color)

#### Gene *58*; Score *0*

Images taken after 5 days at 37 ºC

- + 1 2 3   - + 1 2 3   - + 1 2 3

Dilution factor: $10^{0}$, $10^{-1}$, $10^{-2}$, $10^{-3}$, $10^{-4}$, $10^{-5}$

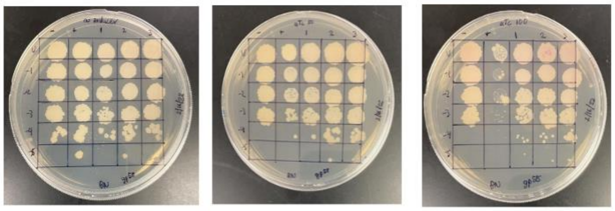

0 ng/ml aTc   10 ng/ml aTc   100 ng/ml aTc

| Lane | Gene ID | Plasmid name | Gene name | Toxic/Non-toxic | Colony color on 100 ng/ml aTc plate* |
|---|---|---|---|---|---|
| - Non-toxic control | -- | pExTra03 | Fruitloop *52* mutant | Non-toxic | + |
| + Toxic control | -- | pExTra02 | Fruitloop *52* | Toxic | - |
| 1 | | pExTra-Lebron 58 | Lebron *58* replicate 1 | Non-toxic | + |
| 2 | | pExTra-Lebron 58 | Lebron *58* replicate 2 | Non-toxic | + |
| 3 | | pExTra-Lebron 58 | Lebron *58* replicate 3 | Non-toxic | + |

*Key: NG (no growth) - (no pink color) +(faint pink color) ++(obvious pink color) +++ (dark pink color)

#### Gene *59*; Score *3*

Images taken after 5 days at 37 ºC

- + 1 2 3   - + 1 2 3   - + 1 2 3

Dilution factor: $10^{0}$, $10^{-1}$, $10^{-2}$, $10^{-3}$, $10^{-4}$, $10^{-5}$

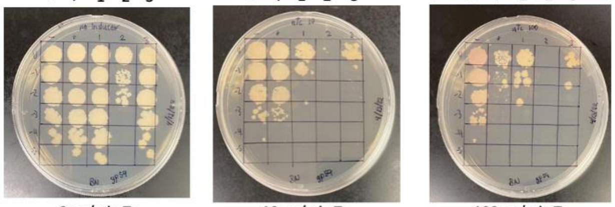

0 ng/ml aTc   10 ng/ml aTc   100 ng/ml aTc

| Lane | Gene ID | Plasmid name | Gene name | Toxic/Non-toxic | Colony color on 100 ng/ml aTc plate* |
|---|---|---|---|---|---|
| + Toxic control | -- | pExTra02 | Fruitloop *52* | Toxic | - |
| - Non-toxic control | -- | pExTra03 | Fruitloop *52* mutant | Non-toxic | + |
| 1 | | pExTra-Lebron 59 | Lebron *59* replicate 1 | Toxic | - |
| 2 | | pExTra-Lebron 59 | Lebron *59* replicate 2 | Toxic | NG |
| 3 | | pExTra-Lebron 59 | Lebron *59* replicate 3 | Toxic | - |

*Key: NG (no growth) - (no pink color) +(faint pink color) ++(obvious pink color) +++ (dark pink color)

#### Gene *60*; Score *1*

Images taken after 5 days at 37 ºC

- + 1 2 3   - + 1 2 3   - + 1 2 3

Dilution factor: $10^{0}$, $10^{-1}$, $10^{-2}$, $10^{-3}$, $10^{-4}$, $10^{-5}$

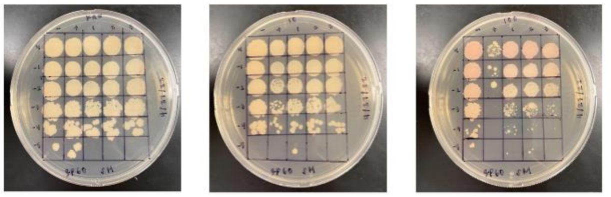

0 ng/ml aTc   10 ng/ml aTc   100 ng/ml aTc

| Lane | Gene ID | Plasmid name | Gene name | Toxic/Non-toxic | Colony color on 100 ng/ml aTc plate* |
|---|---|---|---|---|---|
| + Toxic control | -- | pExTra02 | Fruitloop *52* | Toxic | - |
| - Non-toxic control | -- | pExTra03 | Fruitloop *52* mutant | Non-toxic | + |
| 1 | | pExTra-Lebron 60 | Lebron *60* replicate 1 | Toxic | + |
| 2 | | pExTra-Lebron 60 | Lebron *60* replicate 2 | Toxic | + |
| 3 | | pExTra-Lebron 60 | Lebron *60* replicate 3 | Toxic | + |

*Key: NG (no growth) - (no pink color) +(faint pink color) ++(obvious pink color) +++ (dark pink color)

#### Gene *61*; Score *0*

Images taken after 5 days at 37 ºC

- + 1 2 3   - + 1 2 3   - + 1 2 3

Dilution factor: $10^{0}$, $10^{-1}$, $10^{-2}$, $10^{-3}$, $10^{-4}$, $10^{-5}$

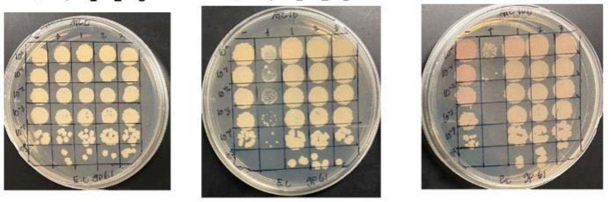

0 ng/ml aTc   10 ng/ml aTc   100 ng/ml aTc

| Lane | Gene ID | Plasmid name | Gene name | Toxic/Non-toxic | Colony color on 100 ng/ml aTc plate* |
|---|---|---|---|---|---|
| + Toxic control | -- | pExTra02 | Fruitloop *52* | Toxic | - |
| - Non-toxic control | -- | pExTra03 | Fruitloop *52* mutant | Non-toxic | + |
| 1 | | pExTra-Lebron 61 | Lebron 61 replicate 1 | Non-toxic | + |
| 2 | | pExTra-Lebron 61 | Lebron 61 replicate 2 | Non-toxic | + |
| 3 | | pExTra-Lebron 61 | Lebron 61 replicate 3 | Non-toxic | + |

*Key: NG (no growth) - (no pink color) +(faint pink color) ++(obvious pink color) +++ (dark pink color)

#### Gene *62*; Score *0*

Images taken after 5 days at 37 ºC

- + 1 2 3   - + 1 2 3   - + 1 2 3

Dilution factor: $10^{0}$, $10^{-1}$, $10^{-2}$, $10^{-3}$, $10^{-4}$, $10^{-5}$

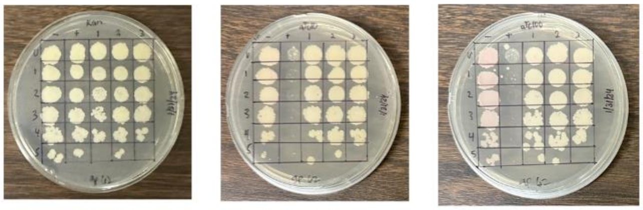

0 ng/ml aTc   10 ng/ml aTc   100 ng/ml aTc

| Lane | Gene ID | Plasmid name | Gene name | Toxic/Non-toxic | Colony color on 100 ng/ml aTc plate* |
|---|---|---|---|---|---|
| - Non-toxic control | -- | pExTra03 | Fruitloop *52* mutant | Non-toxic | + |
| + Toxic control | -- | pExTra02 | Fruitloop *52* | Toxic | - |
| 1 | | pExTra-Lebron 62 | Lebron *62* replicate 1 | Non-toxic | - |
| 2 | | pExTra-Lebron 62 | Lebron *62* replicate 2 | Non-toxic | - |
| 3 | | pExTra-Lebron 62 | Lebron *62* replicate 3 | Non-toxic | - |

*Key: NG (no growth) - (no pink color) +(faint pink color) ++(obvious pink color) +++ (dark pink color)

#### Gene *63*; Score *0*

Images taken after 5 days at 37 ºC

- + 1 2 3   - + 1 2 3   - + 1 2 3

Dilution factor: $10^{0}$, $10^{-1}$, $10^{-2}$, $10^{-3}$, $10^{-4}$, $10^{-5}$

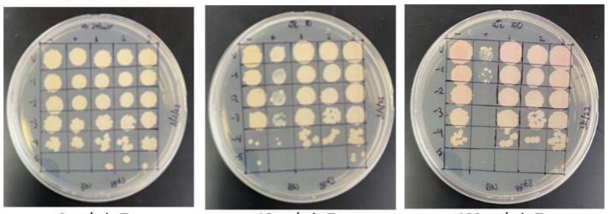

0 ng/ml aTc   10 ng/ml aTc   100 ng/ml aTc

| Lane | Gene ID | Plasmid name | Gene name | Toxic/Non-toxic | Colony color on 100 ng/ml aTc plate* |
|---|---|---|---|---|---|
| - Non-toxic control | -- | pExTra03 | Fruitloop *52* mutant | Non-toxic | + |
| + Toxic control | -- | pExTra02 | Fruitloop *52* | Toxic | - |
| 1 | | pExTra-Lebron 63 | Lebron *63* replicate 1 | Non-toxic | ++ |
| 2 | | pExTra-Lebron 63 | Lebron *63* replicate 2 | Non-toxic | ++ |
| 3 | | pExTra-Lebron 63 | Lebron *63* replicate 3 | Non-toxic | ++ |

*Key: NG (no growth) - (no pink color) +(faint pink color) ++(obvious pink color) +++ (dark pink color)

#### Gene *64*; Score *1*

Images taken after 5 days at 37 ºC

+ - 1 2 3   + - 1 2 3   + - 1 2 3

Dilution factor: $10^{0}$, $10^{-1}$, $10^{-2}$, $10^{-3}$, $10^{-4}$, $10^{-5}$

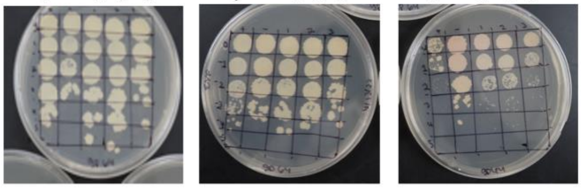

0 ng/ml aTc   10 ng/ml aTc   100 ng/ml aTc

| Lane | Gene ID | Plasmid name | Gene name | Toxic/Non-toxic | Colony color on 100 ng/ml aTc plate* |
|---|---|---|---|---|---|
| + Toxic control | -- | pExTra02 | Fruitloop *52* | Toxic | - |
| - Non-toxic control | -- | pExTra03 | Fruitloop *52* mutant | Non-toxic | ++ |
| 1 | | pExTra-Lebron 64 | Lebron *64* replicate 1 | Toxic | + |
| 2 | | pExTra-Lebron 64 | Lebron *64* replicate 2 | Toxic | + |
| 3 | | pExTra-Lebron 64 | Lebron *64* replicate 3 | Toxic | + |

*Key: NG (no growth) - (no pink color) +(faint pink color) ++(obvious pink color) +++ (dark pink color)

#### Gene *65*; Score *0*

Images taken after 5 days at 37 ºC

- + 1 2 3   - + 1 2 3   - + 1 2 3

Dilution factor: $10^{0}$, $10^{-1}$, $10^{-2}$, $10^{-3}$, $10^{-4}$, $10^{-5}$

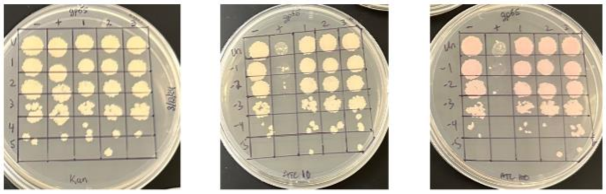

0 ng/ml aTc   10 ng/ml aTc   100 ng/ml aTc

| Lane | Gene ID | Plasmid name | Gene name | Toxic/Non-toxic | Colony color on 100 ng/ml aTc plate* |
|---|---|---|---|---|---|
| - Non-toxic control | -- | pExTra03 | Fruitloop *52* mutant | Non-toxic | ++ |
| + Toxic control | -- | pExTra02 | Fruitloop *52* | Toxic | - |
| 1 | | pExTra-Lebron 65 | Lebron *65* replicate 1 | Non-toxic | ++ |
| 2 | | pExTra-Lebron 65 | Lebron *65* replicate 2 | Non-toxic | ++ |
| 3 | | pExTra-Lebron 65 | Lebron *65* replicate 3 | Non-toxic | ++ |

*Key: NG (no growth) - (no pink color) +(faint pink color) ++(obvious pink color) +++ (dark pink color)

Images taken after 5 days at 37 ºC

#### Gene *66*; Score *1*

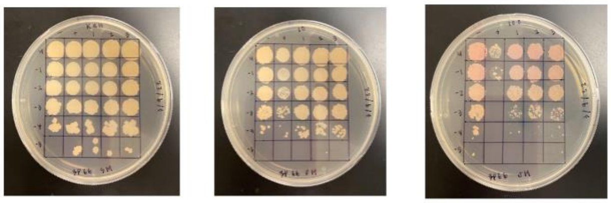


0 ng/ml aTc | 10 ng/ml aTc | 100 ng/ml aTc

| Lane | Gene ID | Plasmid name | Gene name | Toxic/Non-toxic | Colony color on 100 ng/ml aTc plate* |
|---|---|---|---|---|---|
| + Toxic control | -- | pExTra02 | Fruitloop *52* | Toxic | - |
| - Non-toxic control | -- | pExTra03 | Fruitloop *52* mutant | Non-toxic | ++ |
| 1 | | pExTra-Lebron 66 | Lebron *66* replicate 1 | Toxic | ++ |
| 2 | | pExTra-Lebron 66 | Lebron *66* replicate 2 | Toxic | ++ |
| 3 | | pExTra-Lebron 66 | Lebron 66 replicate 3 | Toxic | ++ |

*Key: NG (no growth) - (no pink color) +(faint pink color) ++(obvious pink color) +++ (dark pink color)

Images taken after 5 days at 37 ºC

#### Gene *67*; Score *1*

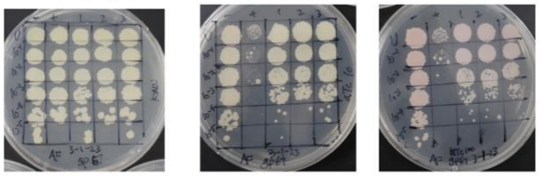


0 ng/ml aTc | 10 ng/ml aTc | 100 ng/ml aTc

| Lane | Gene ID | Plasmid name | Gene name | Toxic/Non-toxic | Colony color on 100 ng/ml aTc plate* |
|---|---|---|---|---|---|
| + Toxic control | -- | pExTra02 | Fruitloop *52* | Toxic | - |
| - Non-toxic control | -- | pExTra03 | Fruitloop *52* mutant | Non-toxic | ++ |
| 1 | | pExTra-Lebron *67* | Lebron *67* replicate 1 | Toxic | + |
| 2 | | pExTra-Lebron *67* | Lebron *67* replicate 2 | Toxic | + |
| 3 | | pExTra-Lebron *67* | Lebron *67* replicate 3 | Toxic | + |

*Key: NG (no growth) - (no pink color) +(faint pink color) ++(obvious pink color) +++ (dark pink color)

Images taken after 5 days at 37 ºC

#### Gene *68*; Score *1*

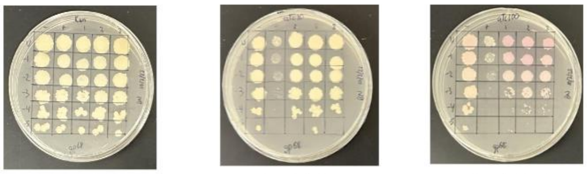


0 ng/ml aTc | 10 ng/ml aTc | 100 ng/ml aTc

| Lane | Gene ID | Plasmid name | Gene name | Toxic/Non-toxic | Colony color on 100 ng/ml aTc plate* |
|---|---|---|---|---|---|
| - Non-toxic control | -- | pExTra03 | Fruitloop *52* mutant | Non-toxic | + |
| + Toxic control | -- | pExTra02 | Fruitloop *52* | Toxic | - |
| 1 | | pExTra-Lebron 68 | Lebron *68* replicate 1 | Toxic | ++ |
| 2 | | pExTra-Lebron 68 | Lebron *68* replicate 2 | Toxic | ++ |
| 3 | | pExTra-Lebron 68 | Lebron *68* replicate 3 | Toxic | ++ |

*Key: NG (no growth) - (no pink color) +(faint pink color) ++(obvious pink color) +++ (dark pink color)

Images taken after 5 days at 37 ºC

#### Gene *69*; Score *0*

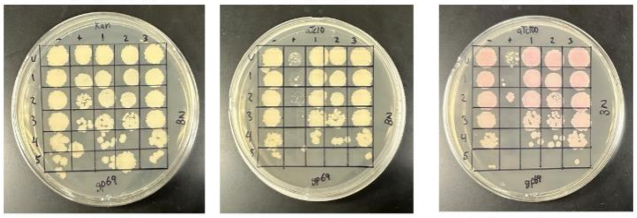


0 ng/ml aTc | 10 ng/ml aTc | 100 ng/ml aTc

| Lane | Gene ID | Plasmid name | Gene name | Toxic/Non-toxic | Colony color on 100 ng/ml aTc plate* |
|---|---|---|---|---|---|
| - Non-toxic control | -- | pExTra03 | Fruitloop *52* mutant | Non-toxic | ++ |
| + Toxic control | -- | pExTra02 | Fruitloop *52* | Toxic | ++ |
| 1 | | pExTra-Lebron 69 | Lebron *69* replicate 1 | Non-toxic | ++ |
| 2 | | pExTra-Lebron 69 | Lebron *69* replicate 2 | Non-toxic | ++ |
| 3 | | pExTra-Lebron 69 | Lebron *69* replicate 3 | Non-toxic | ++ |

*Key: NG (no growth) - (no pink color) +(faint pink color) ++(obvious pink color) +++ (dark pink color)

Images taken after 5 days at 37 ºC

#### Gene *70*; Score *2*

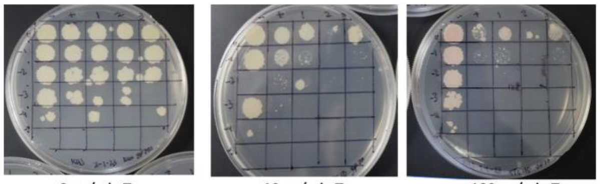


0 ng/ml aTc | 10 ng/ml aTc | 100 ng/ml aTc

| Lane | Gene ID | Plasmid name | Gene name | Toxic/Non-toxic | Colony color on 100 ng/ml aTc plate* |
|---|---|---|---|---|---|
| + Toxic control | -- | pExTra02 | Fruitloop *52* | Toxic | - |
| - Non-toxic control | -- | pExTra03 | Fruitloop *52* mutant | Non-toxic | + |
| 1 | | pExTra-Lebron *70* | Lebron *70* replicate 1 | Toxic | + |
| 2 | | pExTra-Lebron *70* | Lebron *70* replicate 2 | Toxic | - |
| 3 | | pExTra-Lebron *70* | Lebron *70* replicate 3 | Toxic | - |

*Key: NG (no growth) - (no pink color) +(faint pink color) ++(obvious pink color) +++ (dark pink color)

Images taken after 5 days at 37 ºC

#### Gene *71*; Score *3*

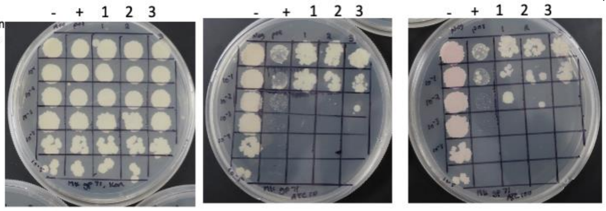


0 ng/ml aTc | 10 ng/ml aTc | 100 ng/ml aTc

| Lane | Gene ID | Plasmid name | Gene name | Toxic/Non-toxic | Colony color on 100 ng/ml aTc plate* |
|---|---|---|---|---|---|
| + Toxic control | -- | pExTra02 | Fruitloop 52 | Toxic | - |
| - Non-toxic control | -- | pExTra03 | Fruitloop 52 mutant | Non-toxic | + |
| 1 | | pExTra-Lebron 71 | Lebron *71* replicate 1 | Toxic | - |
| 2 | | pExTra-Lebron 71 | Lebron *71* replicate 2 | Toxic | - |
| 3 | | pExTra-Lebron 71 | Lebron *71* replicate 3 | Toxic | - |

*Key: NG (no growth) - (no pink color) +(faint pink color) ++(obvious pink color) +++ (dark pink color)

Images taken after 5 days at 37 ºC

#### Gene *72*; Score *0*

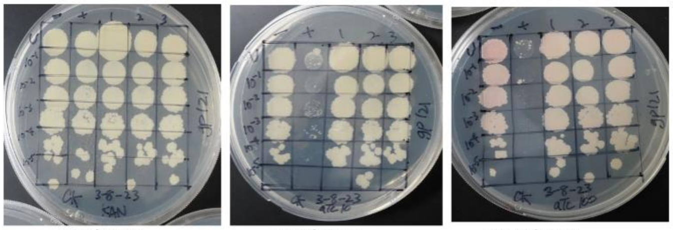


0 ng/ml aTc | 10 ng/ml aTc | 100 ng/ml aTc

| Lane | Gene ID | Plasmid name | Gene name | Toxic/Non-toxic | Colony color on 100 ng/ml aTc plate* |
|---|---|---|---|---|---|
| - Non-toxic control | -- | pExTra03 | Fruitloop *52* mutant | Non-toxic | ++ |
| + Toxic control | -- | pExTra02 | Fruitloop *52* | Toxic | – |
| 1 | – | pExTra-Lebron 72 | Lebron *72* replicate 1 | Non-toxic | – |
| 2 | – | pExTra-Lebron 72 | Lebron *72* replicate 2 | Non-toxic | – |
| 3 | – | pExTra-Lebron 72 | Lebron *72* replicate 3 | Non-toxic | – |

*Key: NG (no growth) - (no pink color) +(faint pink color) ++(obvious pink color) +++ (dark pink color)

Images taken after 5 days at 37 ºC

#### Gene *73*; Score *2*

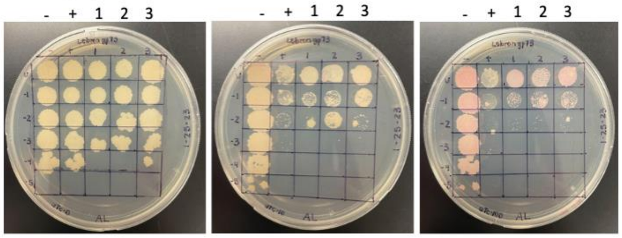


0 ng/ml aTc | 10 ng/ml aTc | 100 ng/ml aTc

| Lane | Gene ID | Plasmid name | Gene name | Toxic/Non-toxic | Colony color on 100 ng/ml aTc plate* |
|---|---|---|---|---|---|
| + Toxic control | -- | pExTra02 | Fruitloop *52* | Toxic | - |
| - Non-toxic control | -- | pExTra03 | Fruitloop *52* mutant | Non-toxic | + |
| 1 | | pExTra-Lebron73 | Lebron 73 replicate 1 | Toxic | ++ |
| 2 | | pExTra-Lebron73 | Lebron 73 replicate 2 | Toxic | ++ |
| 3 | | pExTra-Lebron73 | Lebron 73 replicate 3 | Toxic | ++ |

*Key: NG (no growth) - (no pink color) +(faint pink color) ++(obvious pink color) +++ (dark pink color)

#### Gene *74*; Score *1*

Images taken after 5 days at 37 ºC

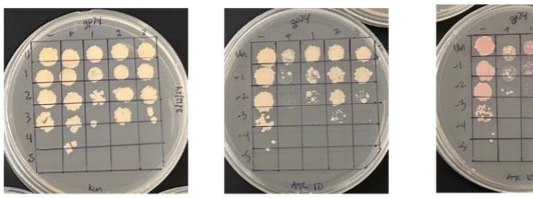


| Lane | Gene ID | Plasmid name | Gene name | Toxic/Non-toxic | Colony color on 100 ng/ml aTc plate* |
|---|---|---|---|---|---|
| - Non-toxic control | -- | pExTra03 | Fruitloop *52* mutant | Non-toxic | ++ |
| + Toxic control | -- | pExTra02 | Fruitloop *52* | Toxic | - |
| 1 | | pExTra-Lebron 74 | Lebron *74* replicate 1 | Toxic | +++ |
| 2 | | pExTra-Lebron 74 | Lebron *74* replicate 2 | Toxic | +++ |
| 3 | | pExTra-Lebron 74 | Lebron *74* replicate 3 | Toxic | +++ |

*Key: NG (no growth) - (no pink color) +(faint pink color) ++(obvious pink color) +++ (dark pink color)

#### Gene *75*; Score *0*

Images taken after 5 days at 37 ºC

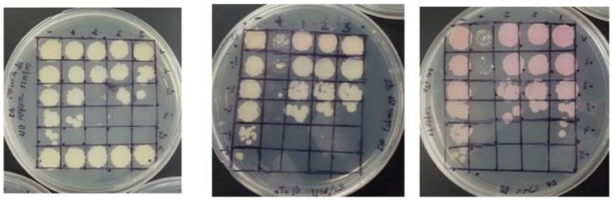


| Lane | Gene ID | Plasmid name | Gene name | Toxic/Non-toxic | Colony color on 100 ng/ml aTc plate* |
|---|---|---|---|---|---|
| + Toxic control | -- | pExTra02 | Fruitloop *52* | Toxic | - |
| - Non-toxic control | -- | pExTra03 | Fruitloop *52* mutant | Non-toxic | ++ |
| 1 | | pExTra-Lebron 75 | Lebron *75* replicate 1 | Non-toxic | +++ |
| 2 | | pExTra-Lebron 75 | Lebron *75* replicate 2 | Non-toxic | +++ |
| 3 | | pExTra-Lebron 75 | Lebron *75* replicate 3 | Non-toxic | +++ |

*Key: NG (no growth) - (no pink color) +(faint pink color) ++(obvious pink color) +++ (dark pink color)

#### Gene *76*; Score *3*

Images taken after 5 days at 37 ºC

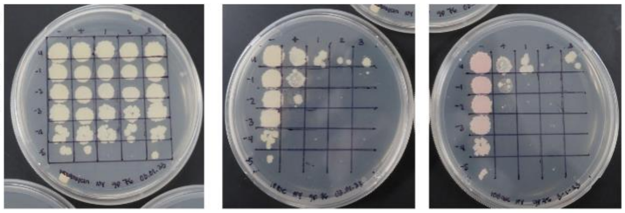


| Lane | Gene ID | Plasmid name | Gene name | Toxic/Non-toxic | Colony color on 100 ng/ml aTc plate* |
|---|---|---|---|---|---|
| + Toxic control | -- | pExTra02 | Fruitloop *52* | Toxic | - |
| - Non-toxic control | -- | pExTra03 | Fruitloop *52* mutant | Non-toxic | + |
| 1 | 131436 | pExTra-Lebron 76 | Lebron 76 replicate 1 | Toxic | - |
| 2 | 131436 | pExTra-Lebron 76 | Lebron 76 replicate 2 | Toxic | - |
| 3 | 131436 | pExTra-Lebron 76 | Lebron 76 replicate 3 | Toxic | - |

*Key: NG (no growth) - (no pink color) +(faint pink color) ++(obvious pink color) +++ (dark pink color)

#### Gene *77*; Score *0*

Images taken after 5 days at 37 ºC

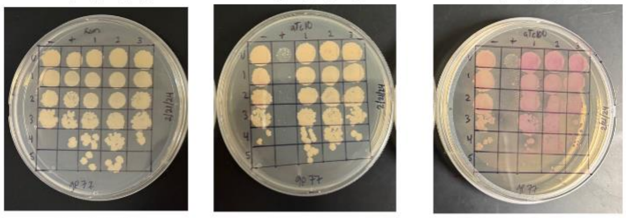


| Lane | Gene ID | Plasmid name | Gene name | Toxic/Non-toxic | Colony color on 100 ng/ml aTc plate* |
|---|---|---|---|---|---|
| - Non-toxic control | -- | pExTra03 | Fruitloop *52* mutant | Non-toxic | ++ |
| + Toxic control | -- | pExTra02 | Fruitloop *52* | Toxic | - |
| 1 | | pExTra-Lebron 77 | Lebron *77* replicate 1 | Non-toxic | +++ |
| 2 | | pExTra-Lebron 77 | Lebron *77* replicate 2 | Non-toxic | +++ |
| 3 | | pExTra-Lebron 77 | Lebron *77* replicate 3 | Non-toxic | +++ |

*Key: NG (no growth) - (no pink color) +(faint pink color) ++(obvious pink color) +++ (dark pink color)

#### Gene *78*; Score *3*

Images taken after 5 days at 37 ºC

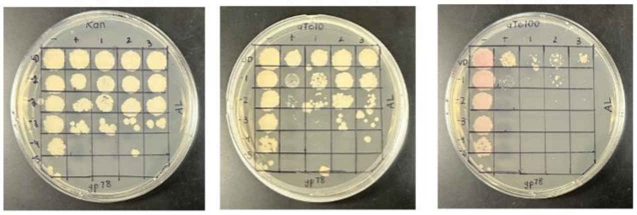


| Lane | Gene ID | Plasmid name | Gene name | Toxic/Non-toxic | Colony color on 100 ng/ml aTc plate* |
|---|---|---|---|---|---|
| - Non-toxic control | -- | pExTra03 | Fruitloop *52* mutant | Non-toxic | ++ |
| + Toxic control | -- | pExTra02 | Fruitloop *52* | Toxic | - |
| 1 | | pExTra-Lebron 78 | Lebron *78* replicate 1 | Toxic | - |
| 2 | | pExTra-Lebron 78 | Lebron *78* replicate 2 | Toxic | - |
| 3 | | pExTra-Lebron 78 | Lebron *78* replicate 3 | Toxic | - |

*Key: NG (no growth) - (no pink color) +(faint pink color) ++(obvious pink color) +++ (dark pink color)

#### Gene *79*; Score *0*

Images taken after 5 days at 37 ºC

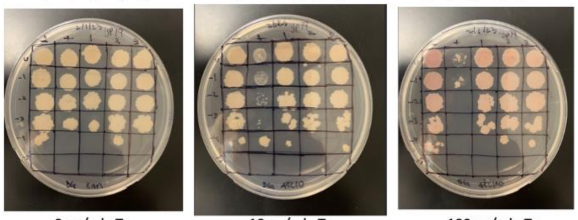


| Lane | Gene ID | Plasmid name | Gene name | Toxic/Non-toxic | Colony color on 100 ng/ml aTc plate* |
|---|---|---|---|---|---|
| + Toxic control | -- | pExTra02 | Fruitloop *52* | Toxic | - |
| - Non-toxic control | -- | pExTra03 | Fruitloop *52* mutant | Non-toxic | +++ |
| 1 | 131436 | pExtra-Lebron 79 | Lebron 79 replicate 1 | Non-toxic | ++ |
| 2 | 131436 | pExTra-Lebron 79 | Lebron 79 replicate 2 | Non-toxic | ++ |
| 3 | 131436 | pExTra-Lebron 79 | Lebron 79 replicate 3 | Non-toxic | ++ |

*Key: NG (no growth) - (no pink color) +(faint pink color) ++(obvious pink color) +++ (dark pink color)

#### Gene *80*; Score *1*

Images taken after 5 days at 37 ºC

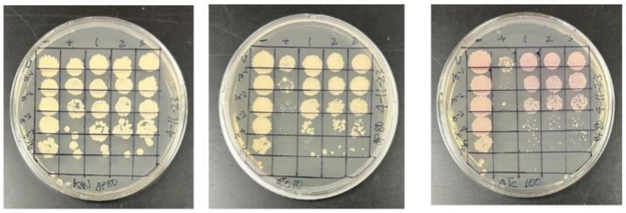


| Lane | Gene ID | Plasmid name | Gene name | Toxic/Non-toxic | Colony color on 100 ng/ml aTc plate* |
|---|---|---|---|---|---|
| - Non-toxic control | -- | pExTra03 | Fruitloop *52* mutant | Non-toxic | ++ |
| + Toxic control | -- | pExTra02 | Fruitloop *52* | Toxic | - |
| 1 | | pExTra-Lebron 80 | Lebron *80* replicate 1 | Toxic | ++ |
| 2 | | pExTra-Lebron 80 | Lebron *80* replicate 2 | Toxic | ++ |
| 3 | | pExTra-Lebron 80 | Lebron *80* replicate 3 | Toxic | ++ |

*Key: NG (no growth) - (no pink color) +(faint pink color) ++(obvious pink color) +++ (dark pink color)

#### Gene *81*; Score *3*

Images taken after 5 days at 37 ºC

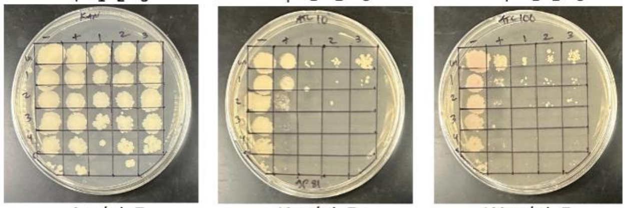


| Lane | Gene ID | Plasmid name | Gene name | Toxic/Non-toxic | Colony color on 100 ng/ml aTc plate* |
|---|---|---|---|---|---|
| - Non-toxic control | -- | pExTra03 | Fruitloop *52* mutant | Non-toxic | + |
| + Toxic control | -- | pExTra02 | Fruitloop *52* | Toxic | - |
| 1 | | pExTra-Lebron 81 | Lebron *81* replicate 1 | Toxic | - |
| 2 | | pExTra-Lebron 81 | Lebron *81* replicate 2 | Toxic | - |
| 3 | | pExTra-Lebron 81 | Lebron *81* replicate 3 | Toxic | - |

*Key: NG (no growth) - (no pink color) +(faint pink color) ++(obvious pink color) +++ (dark pink color)

#### Gene *82*; Score *0*

Images taken after 5 days at 37 ºC

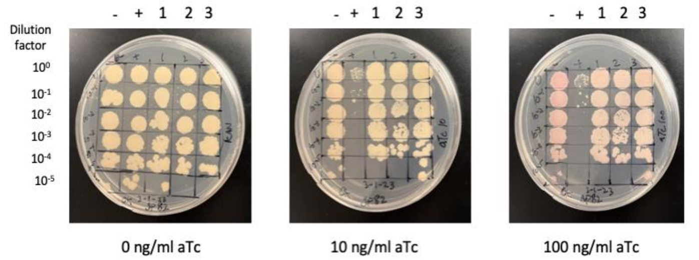


| Lane | Gene ID | Plasmid name | Gene name | Toxic/Non-toxic | Colony color on 100 ng/ml aTc plate* |
|---|---|---|---|---|---|
| - Non-toxic control | -- | pExTra03 | Fruitloop *52* mutant | Non-toxic | ++ |
| + Toxic control | -- | pExTra02 | Fruitloop *52* | Toxic | – |
| 1 | – | pExTra-Lebron 82 | Lebron *82* replicate 1 | Non-toxic | + |
| 2 | – | pExTra-Lebron 82 | Lebron *82* replicate 2 | Non-toxic | + |
| 3 | – | pExTra-Lebron 82 | Lebron *82* replicate 3 | Non-toxic | + |

*Key: NG (no growth) - (no pink color) +(faint pink color) ++(obvious pink color) +++ (dark pink color)

#### Gene *83*; Score *0*

Images taken after 5 days at 37 ºC

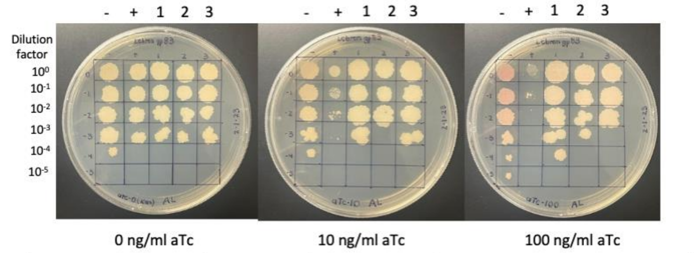


| Lane | Gene ID | Plasmid name | Gene name | Toxic/Non-toxic | Colony color on 100 ng/ml aTc plate* |
|---|---|---|---|---|---|
| + Toxic control | -- | pExTra02 | Fruitloop *52* | Toxic | - |
| - Non-toxic control | -- | pExTra03 | Fruitloop *52* mutant | Non-toxic | + |
| 1 | | pExTra-Lebron83 | Lebron 83 replicate 1 | Non-toxic | - |
| 2 | | pExTra-Lebron83 | Lebron 83 replicate 2 | Non-toxic | - |
| 3 | | pExTra-Lebron83 | Lebron 83 replicate 3 | Non-toxic | - |

*Key: NG (no growth) - (no pink color) +(faint pink color) ++(obvious pink color) +++ (dark pink color)

#### Gene *84*; Score *0*

Images taken after 5 days at 37 ºC

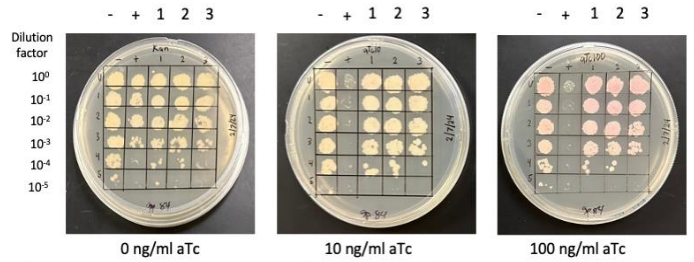


| Lane | Gene ID | Plasmid name | Gene name | Toxic/Non-toxic | Colony color on 100 ng/ml aTc plate* |
|---|---|---|---|---|---|
| - Non-toxic control | -- | pExTra03 | Fruitloop *52* mutant | Non-toxic | ++ |
| + Toxic control | -- | pExTra02 | Fruitloop *52* | Toxic | - |
| 1 | | pExTra-Lebron 84 | Lebron *84* replicate 1 | Non-toxic | ++ |
| 2 | | pExTra-Lebron 84 | Lebron *84* replicate 2 | Non-toxic | ++ |
| 3 | | pExTra-Lebron 84 | Lebron *84* replicate 3 | Non-toxic | ++ |

*Key: NG (no growth) - (no pink color) +(faint pink color) ++(obvious pink color) +++ (dark pink color)

#### Gene *85*; Score *3*

Images taken after 5 days at 37 ºC

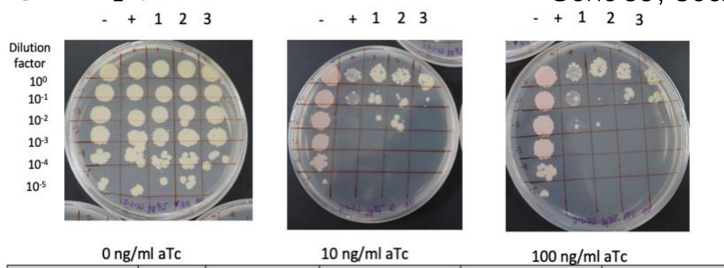


| Lane | Gene ID | Plasmid name | Gene name | Toxic/Non-toxic | Colony color on 100 ng/ml aTc plate* |
|---|---|---|---|---|---|
| + Toxic control | -- | pExTra02 | Fruitloop *52* | Toxic | - |
| - Non-toxic control | -- | pExTra03 | Fruitloop *52* mutant | Non-toxic | ++ |
| 1 | | pExTra-Lebron 85 | Lebron *85* replicate 1 | Toxic | - |
| 2 | | pExTra-Lebron 85 | Lebron *85* replicate 2 | Toxic | - |
| 3 | | pExTra-Lebron 85 | Lebron *85* replicate 3 | Toxic | - |

*Key: NG (no growth) - (no pink color) +(faint pink color) ++(obvious pink color) +++ (dark pink color)

#### Gene *86*; Score *1*

Images taken after 5 days at 37 ºC

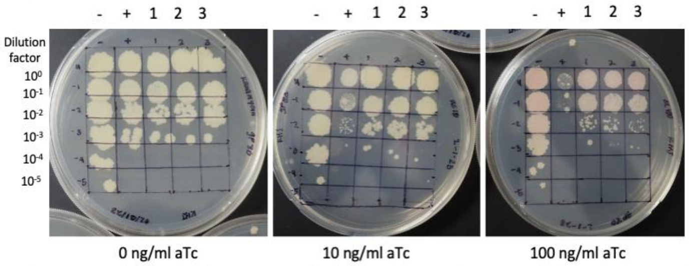


| Lane | Gene ID | Plasmid name | Gene name | Toxic/Non-toxic | Colony color on 100 ng/ml aTc plate* |
|---|---|---|---|---|---|
| + Toxic control | -- | pExTra02 | Fruitloop *52* | Toxic | - |
| - Non-toxic control | -- | pExTra03 | Fruitloop *52* mutant | Non-toxic | + |
| 1 | | pExTra-Lebron 86 | Lebron 86 replicate 1 | Toxic | + |
| 2 | | pExTra-Lebron 86 | Lebron 86 replicate 2 | Toxic | + |
| 3 | | pExTra-Lebron 86 | Lebron 86 replicate 3 | Toxic | + |

*Key: NG (no growth) - (no pink color) +(faint pink color) ++(obvious pink color) +++ (dark pink color)

#### Gene *87*; Score *0*

Images taken after 5 days at 37 ºC

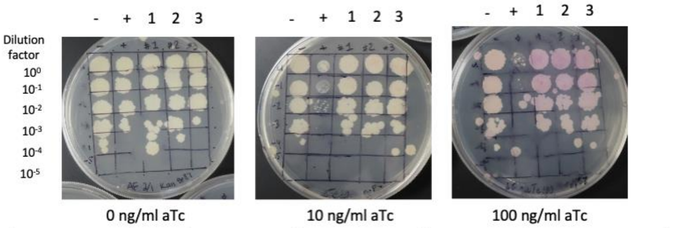


| Lane | Gene ID | Plasmid name | Gene name | Toxic/Non-toxic | Colony color on 100 ng/ml aTc plate* |
|---|---|---|---|---|---|
| + Toxic control | -- | pExTra02 | Fruitloop *52* | Toxic | - |
| - Non-toxic control | -- | pExTra03 | Fruitloop *52* mutant | Non-toxic | + |
| 1 | | pExTra-Lebron *87* | Lebron *87* replicate 1 | Non-toxic | ++ |
| 2 | | pExTra-Lebron *87* | Lebron *87* replicate 2 | Non-toxic | ++ |
| 3 | | pExTra-Lebron *87* | Lebron *87* replicate 3 | Non-toxic | ++ |

*Key: NG (no growth) - (no pink color) +(faint pink color) ++(obvious pink color) +++ (dark pink color)

#### Gene *88*; Score *0*

Images taken after 5 days at 37 ºC

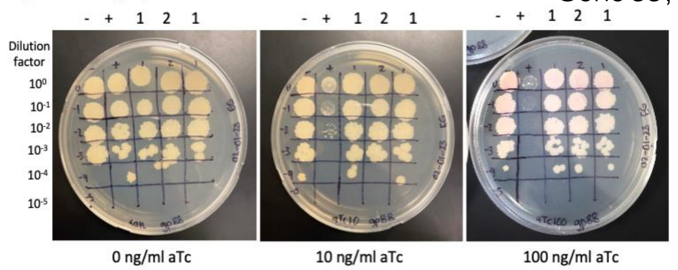


| Lane | Gene ID | Plasmid name | Gene name | Toxic/Non-toxic | Colony color on 100 ng/ml aTc plate* |
|---|---|---|---|---|---|
| + Toxic control | -- | pExTra02 | Fruitloop *52* | Toxic | + |
| - Non-toxic control | -- | pExTra03 | Fruitloop *52* mutant | Non-toxic | - |
| 1 | | pExTra-Lebron 88 | Lebron 88 replicate 1 | Non-toxic | + |
| 2 | | pExTra-Lebron 88 | Lebron 88 replicate 2 | Non-toxic | + |
| 3 | | pExTra-Lebron 88 | Lebron 88 replicate 1 | Non-toxic | + |

*Key: NG (no growth) - (no pink color) +(faint pink color) ++(obvious pink color) +++ (dark pink color)

#### Gene *89*; Score *2*

Images taken after 5 days at 37 ºC

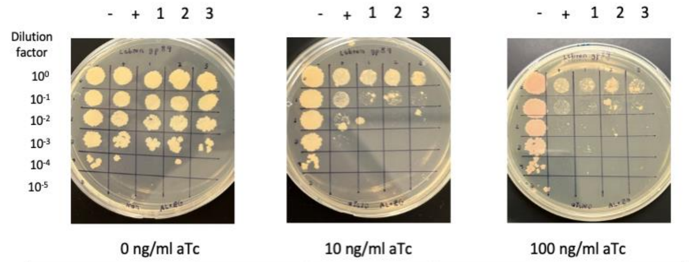


| Lane | Gene ID | Plasmid name | Gene name | Toxic/Non-toxic | Colony color on 100 ng/ml aTc plate* |
|---|---|---|---|---|---|
| - Non-toxic control | -- | pExTra03 | Fruitloop *52* mutant | Non-toxic | ++ |
| + Toxic control | -- | pExTra02 | Fruitloop *52* | Toxic | - |
| 1 | | pExTra-Lebron 89 | Lebron *89* replicate 1 | Toxic | + |
| 2 | | pExTra-Lebron 89 | Lebron *89* replicate 2 | Toxic | + |
| 3 | | pExTra-Lebron 89 | Lebron *89* replicate 3 | Toxic | + |

*Key: NG (no growth) - (no pink color) +(faint pink color) ++(obvious pink color) +++ (dark pink color)

#### Gene *90*; Score *0*

Images taken after 5 days at 37 ºC

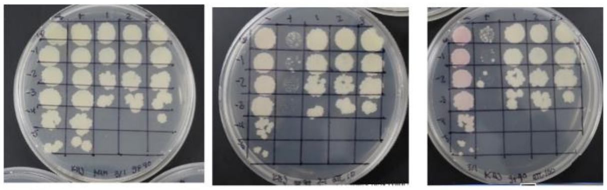


| Lane | Gene ID | Plasmid name | Gene name | Toxic/Non-toxic | Colony color on 100 ng/ml aTc plate* |
|---|---|---|---|---|---|
| + Toxic control | -- | pExTra02 | Fruitloop *52* | Toxic | - |
| - Non-toxic control | -- | pExTra03 | Fruitloop *52* mutant | Non-toxic | ++ |
| 1 | | pExTra-Lebron *90* | Lebron *90* replicate 1 | Non-toxic | - |
| 2 | | pExTra-Lebron *90* | Lebron *90* replicate 2 | Non-toxic | - |
| 3 | | pExTra-Lebron *90* | Lebron *90* replicate 3 | Non-toxic | - |

*Key: NG (no growth) - (no pink color) +(faint pink color) ++(obvious pink color) +++ (dark pink color)

#### Gene *91* Score *0*

Images taken after 5 days at 37 ºC

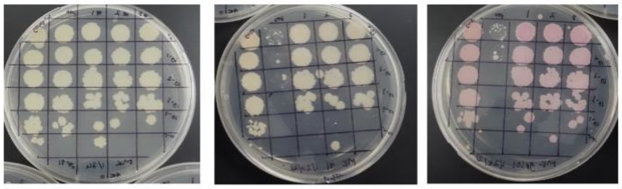


| Lane | Gene ID | Plasmid name | Gene name | Toxic/Non-toxic | Colony color on 100 ng/ml aTc plate* |
|---|---|---|---|---|---|
| + Toxic control | -- | pExTra02 | Fruitloop 52 | Toxic | - |
| - Non-toxic control | -- | pExTra03 | Fruitloop 52 mutant X | Non-toxic | ++ |
| 1 | | pExTra-Lebron 91 | Lebron *91* replicate 1 | Non-toxic | ++ |
| 2 | | pExTra-Lebron 91 | Lebron *91* replicate 2 | Non-toxic | ++ |
| 3 | | pExTra-Lebron 91 | Lebron *91* replicate 3 | Non-toxic | ++ |

*Key: NG (no growth) - (no pink color) +(faint pink color) ++(obvious pink color) +++ (dark pink color)

#### Gene *92*; Score *0*

Images taken after 5 days at 37 ºC

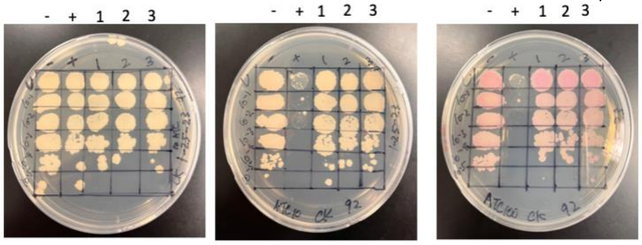


| Lane | Gene ID | Plasmid name | Gene name | Toxic/Non-toxic | Colony color on 100 ng/ml aTc plate* |
|---|---|---|---|---|---|
| - Non-toxic control | -- | pExTra03 | Fruitloop *52* mutant | Non-toxic | + |
| + Toxic control | -- | pExTra02 | Fruitloop *52* | Toxic | - |
| 1 | – | pExTra-Lebron 92 | Lebron *92* replicate 1 | Non-toxic | ++ |
| 2 | – | pExTra-Lebron 92 | Lebron *92* replicate 2 | Non-toxic | ++ |
| 3 | – | pExTra-Lebron 92 | Lebron *92* replicate 3 | Non-toxic | ++ |

*Key: NG (no growth) - (no pink color) +(faint pink color) ++(obvious pink color) +++ (dark pink color)

#### Gene *93*; Score *3*

Images taken after 5 days at 37 ºC

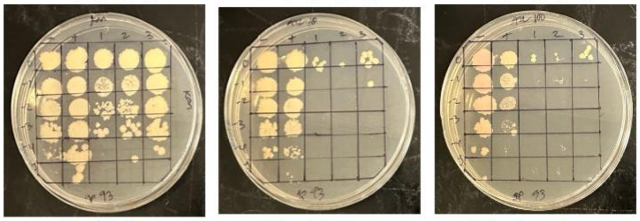


| Lane | Gene ID | Plasmid name | Gene name | Toxic/Non-toxic | Colony color on 100 ng/ml aTc plate* |
|---|---|---|---|---|---|
| - Non-toxic control | -- | pExTra03 | Fruitloop *52* mutant | Non-toxic | + |
| + Toxic control | -- | pExTra02 | Fruitloop *52* | Toxic | - |
| 1 | | pExTra-Lebron 93 | Lebron *93* replicate 1 | Toxic | + |
| 2 | | pExTra-Lebron 93 | Lebron *93* replicate 2 | Toxic | + |
| 3 | | pExTra-Lebron 93 | Lebron *93* replicate 3 | Toxic | + |

*Key: NG (no growth) - (no pink color) +(faint pink color) ++(obvious pink color) +++ (dark pink color)

#### Gene *94*; Score *0*

Images taken after 5 days at 37 ºC

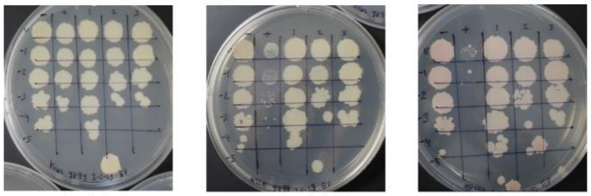


| Lane | Gene ID | Plasmid name | Gene name | Toxic/Non-toxic | Colony color on 100 ng/ml aTc plate* |
|---|---|---|---|---|---|
| - Non-toxic control | -- | pExTra03 | Fruitloop *52* mutant | Non-toxic | + |
| + Toxic control | -- | pExTra02 | Fruitloop *52* | Toxic | - |
| 1 | – | pExTra-Lebron 94 | Lebron *94* replicate 1 | Non-toxic | + |
| 2 | – | pExTra-Lebron 94 | Lebron *94* replicate 2 | Non-toxic | + |
| 3 | – | pExTra-Lebron 94 | Lebron *94* replicate 3 | Non-toxic | + |

*Key: NG (no growth) - (no pink color) +(faint pink color) ++(obvious pink color) +++ (dark pink color)

#### Gene *95*; Score *0*

Images taken after 5 days at 37 ºC

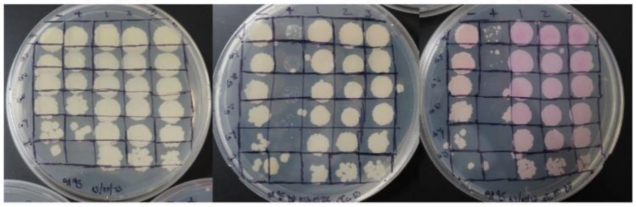


| Lane | Gene ID | Plasmid name | Gene name | Toxic/Non-toxic | Colony color on 100 ng/ml aTc plate* |
|---|---|---|---|---|---|
| + Toxic control | -- | pExTra02 | Fruitloop *52* | Toxic | - |
| - Non-toxic control | -- | pExTra03 | Fruitloop *52* mutant | Non-toxic | ++ |
| 1 | -- | pExTra-Lebron 95 | Lebron *95* replicate 1 | Non-toxic | +++ |
| 2 | -- | pExTra-Lebron 95 | Lebron *95* replicate 2 | Non-toxic | +++ |
| 3 | -- | pExTra-Lebron 95 | Lebron *95* replicate 3 | Non-toxic | +++ |

*Key: NG (no growth) - (no pink color) +(faint pink color) ++(obvious pink color) +++ (dark pink color)

#### Gene *96*; Score *0*

Images taken after 5 days at 37 ºC

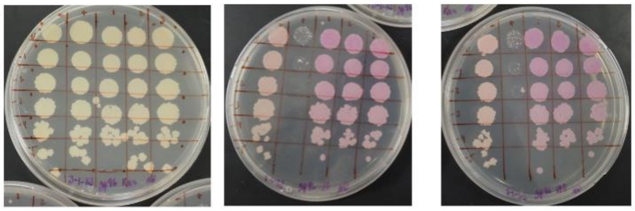


| Lane | Gene ID | Plasmid name | Gene name | Toxic/Non-toxic | Colony color on 100 ng/ml aTc plate* |
|---|---|---|---|---|---|
| + Toxic control | -- | pExTra02 | Fruitloop *52* | Toxic | + |
| - Non-toxic control | -- | pExTra03 | Fruitloop 52 mutant | Non-toxic | - |
| 1 | | pExTra-Lebron 96 | Lebron 96 replicate 1 | Non-toxic | +++ |
| 2 | | pExTra-Lebron 96 | Lebron 96 replicate 2 | Non-toxic | +++ |
| 3 | | pExTra-Lebron 96 | Lebron 96 replicate 3 | Non-toxic | +++ |

*Key: NG (no growth) - (no pink color) +(faint pink color) ++(obvious pink color) +++ (dark pink color)

#### Gene *97*; Score *0*

Images taken after 5 days at 37 ºC

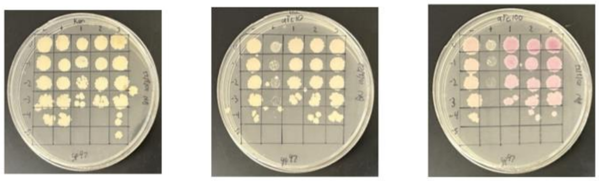


| Lane | Gene ID | Plasmid name | Gene name | Toxic/Non-toxic | Colony color on 100 ng/ml aTc plate* |
|---|---|---|---|---|---|
| - Non-toxic control | -- | pExTra03 | Fruitloop *52* mutant | Non-toxic | + |
| + Toxic control | -- | pExTra02 | Fruitloop *52* | Toxic | - |
| 1 | | pExTra-Lebron 97 | Lebron *97* replicate 1 | Non-toxic | ++ |
| 2 | | pExTra-Lebron 97 | Lebron *97* replicate 2 | Non-toxic | ++ |
| 3 | | pExTra-Lebron 97 | Lebron *97* replicate 3 | Non-toxic | +++ |

*Key: NG (no growth) - (no pink color) +(faint pink color) ++(obvious pink color) +++ (dark pink color)

#### Gene *98*; Score *0*

Images taken after 5 days at 37 ºC

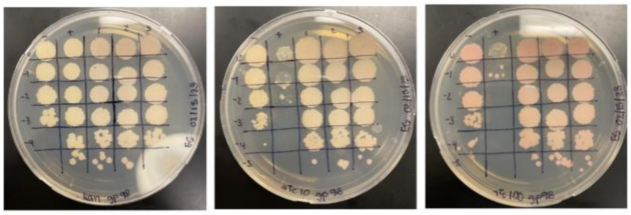


0 ng/ml aTc 10 ng/ml aTc 100 ng/ml aTc

| Lane | Gene ID | Plasmid name | Gene name | Toxic/Non-toxic | Colony color on 100 ng/ml aTc plate* |
|---|---|---|---|---|---|
| + Toxic control | -- | pExTra02 | Fruitloop *52* | Toxic | - |
| - Non-toxic control | -- | pExTra03 | Fruitloop *52* mutant | Non-toxic | + |
| 1 | | pExTra-Lebron 98 | Lebron 98 replicate 1 | Non-Toxic | + |
| 2 | | pExTra-Lebron 98 | Lebron 98 replicate 2 | Non-Toxic | + |
| 3 | | pExTra-Lebron 98 | Lebron *98* replicate 3 | Non-Toxic | + |

*Key: NG (no growth) - (no pink color) +(faint pink color) ++(obvious pink color) +++ (dark pink color)

#### Gene *99*; Score *3*

Images taken after 5 days at 37 ºC

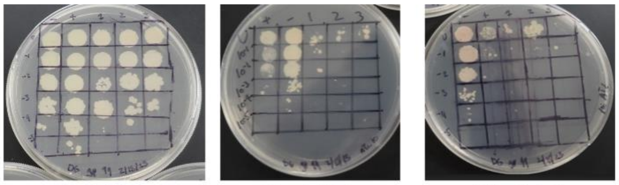


0 ng/ml aTc 10 ng/ml aTc 100 ng/ml aTc

| Lane | Gene ID | Plasmid name | Gene name | Toxic/Non-toxic | Colony color on 100 ng/ml aTc plate* |
|---|---|---|---|---|---|
| + Toxic control | -- | pExTra02 | Fruitloop *52* | Toxic | - |
| - Non-toxic control | -- | pExTra03 | Fruitloop *52* mutant | Non-toxic | + |
| 1 | | pExtra-Lebron 99 | Lebron 99 replicate 1 | Toxic | - |
| 2 | | pExTra-Lebron 99 | Lebron 99 replicate 2 | Toxic | - |
| 3 | | pExTra-Lebron 99 | Lebron 99 replicate 3 | Toxic | NG |

*Key: NG (no growth) - (no pink color) +(faint pink color) ++(obvious pink color) +++ (dark pink color)

#### Gene *100*; Score *1*

Images taken after 5 days at 37 ºC

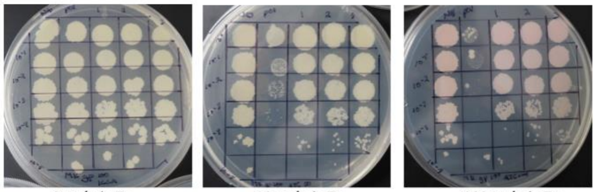


0 ng/ml aTc 10 ng/ml aTc 100 ng/ml aTc

| Lane | Gene ID | Plasmid name | Gene name | Toxic/Non-toxic | Colony color on 100 ng/ml aTc plate* |
|---|---|---|---|---|---|
| + Toxic control | -- | pExTra02 | Fruitloop *52* | Toxic | - |
| - Non-toxic control | -- | pExTra03 | Fruitloop *52* mutant | Non-toxic | + |
| 1 | | pExTra-Lebron *100* | Lebron *100* replicate 1 | Toxic | ++ |
| 2 | | pExTra-Lebron *100* | Lebron *100* replicate 2 | Toxic | ++ |
| 3 | | pExTra-Lebron *100* | Lebron *100* replicate 3 | Toxic | ++ |

*Key: NG (no growth) - (no pink color) +(faint pink color) ++(obvious pink color) +++ (dark pink color)

#### Gene *101*; Score *0*

Images taken after 5 days at 37 ºC

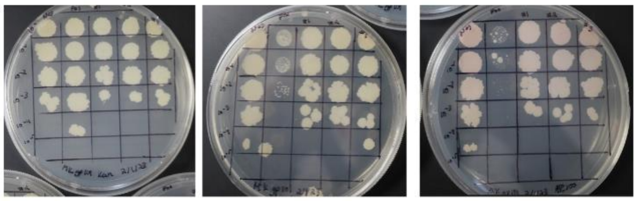


0 ng/ml aTc 10 ng/ml aTc 100 ng/ml aTc

| Lane | Gene ID | Plasmid name | Gene name | Toxic/Non-toxic | Colony color on 100 ng/ml aTc plate* |
|---|---|---|---|---|---|
| + Toxic control | -- | pExTra02 | Fruitloop 52 | Toxic | - |
| - Non-toxic control | -- | pExTra03 | Fruitloop 52 mutant X | Non-toxic | + |
| 1 | | pExTra-Lebron 101 | Lebron *101* replicate 1 | Non-toxic | + |
| 2 | | pExTra-Lebron 101 | Lebron *101* replicate 2 | Non-toxic | + |
| 3 | | pExTra-Lebron 101 | Lebron *101* replicate 3 | Non-toxic | + |

*Key: NG (no growth) - (no pink color) +(faint pink color) ++(obvious pink color) +++ (dark pink color)

#### Gene 104; Score *0*

Images taken after 5 days at 37 ºC

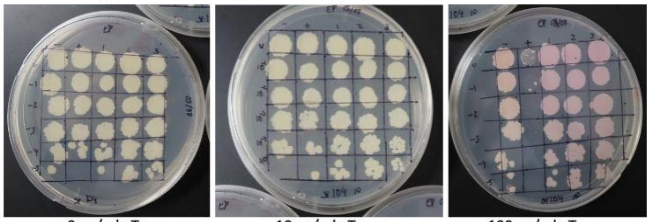


0 ng/ml aTc 10 ng/ml aTc 100 ng/ml aTc

| Lane | Gene ID | Plasmid name | Gene name | Toxic/Non-toxic | Colony color on 100 ng/ml aTc plate* |
|---|---|---|---|---|---|
| - Non-toxic control | -- | pExTra03 | Fruitloop *52* mutant | Non-toxic | + |
| + Toxic control | -- | pExTra02 | Fruitloop *52* | Toxic | - |
| 1 | – | pExTra-Lebron 104 | Lebron *104* replicate 1 | Non-toxic | ++ |
| 2 | – | pExTra-Lebron 104 | Lebron *104* replicate 2 | Non-toxic | ++ |
| 3 | – | pExTra-Lebron 104 | Lebron *104* replicate 3 | Non-toxic | ++ |

*Key: NG (no growth) - (no pink color) +(faint pink color) ++(obvious pink color) +++ (dark pink color)

#### Gene *110*; Score *0*

Images taken after 5 days at 37 ºC

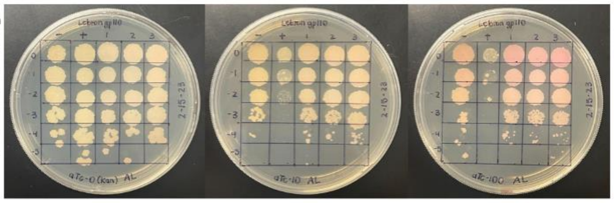


0 ng/ml aTc 10 ng/ml aTc 100 ng/ml aTc

| Lane | Gene ID | Plasmid name | Gene name | Toxic/Non-toxic | Colony color on 100 ng/ml aTc plate* |
|---|---|---|---|---|---|
| + Toxic control | -- | pExTra02 | Fruitloop *52* | Toxic | - |
| - Non-toxic control | -- | pExTra03 | Fruitloop *52* mutant | Non-toxic | + |
| 1 | | pExTra-Lebron110 | Lebron 110 replicate 1 | Non-toxic | - |
| 2 | | pExTra-Lebron110 | Lebron 110 replicate 2 | Non-toxic | - |
| 3 | | pExTra-Lebron110 | Lebron 110 replicate 3 | Non-toxic | + |

*Key: NG (no growth) - (no pink color) +(faint pink color) ++(obvious pink color) +++ (dark pink color)

#### Gene *111*; Score *0*

Images taken after 5 days at 37 ºC

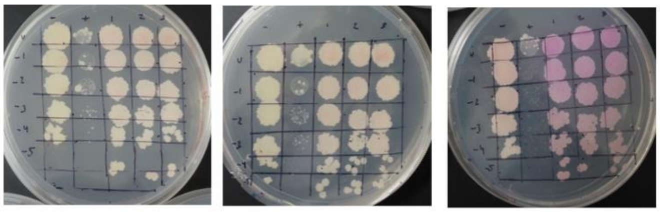


0 ng/ml aTc 10 ng/ml aTc 100 ng/ml aTc

| Lane | Gene ID | Plasmid name | Gene name | Toxic/Non-toxic | Colony color on 100 ng/ml aTc plate* |
|---|---|---|---|---|---|
| - Non-toxic control | -- | pExTra03 | Fruitloop *52* mutant | Non-toxic | + |
| + Toxic control | -- | pExTra02 | Fruitloop *52* | Toxic | - |
| 1 | – | pExTra-Lebron 111 | Lebron *111* replicate 1 | Non-toxic | +++ |
| 2 | – | pExTra-Lebron 111 | Lebron *111* replicate 2 | Non-toxic | +++ |
| 3 | – | pExTra-Lebron 111 | Lebron *111* replicate 3 | Non-toxic | +++ |

*Key: NG (no growth) - (no pink color) +(faint pink color) ++(obvious pink color) +++ (dark pink color)

#### Gene *113*; Score *1*

Images taken after 5 days at 37 ºC

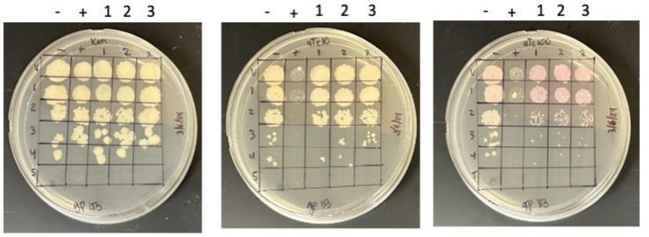


0 ng/ml aTc 10 ng/ml aTc 100 ng/ml aTc

| Lane | Gene ID | Plasmid name | Gene name | Toxic/Non-toxic | Colony color on 100 ng/ml aTc plate* |
|---|---|---|---|---|---|
| - Non-toxic control | -- | pExTra03 | Fruitloop *52* mutant | Non-toxic | + |
| + Toxic control | -- | pExTra02 | Fruitloop *52* | Toxic | - |
| 1 | | pExTra-Lebron 113 | Lebron *113* replicate 1 | Toxic | ++ |
| 2 | | pExTra-Lebron 113 | Lebron *113* replicate 2 | Toxic | ++ |
| 3 | | pExTra-Lebron 113 | Lebron *113* replicate 3 | Toxic | ++ |

*Key: NG (no growth) - (no pink color) +(faint pink color) ++(obvious pink color) +++ (dark pink color)

#### Gene 115; Score *0*

Images taken after 5 days at 37 ºC

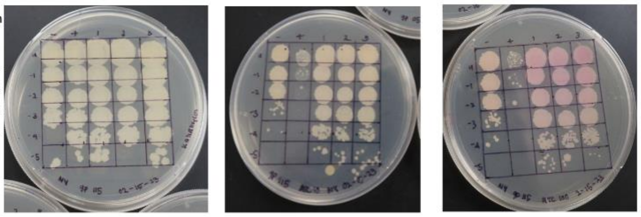


| Lane | Gene ID | Plasmid name | Gene name | Toxic/Non-toxic | Colony color on 100 ng/ml aTc plate* |
|---|---|---|---|---|---|
| + Toxic control | -- | pExTra02 | Fruitloop *52* | Toxic | - |
| - Non-toxic control | -- | pExTra03 | Fruitloop *52* mutant | Non-toxic | + |
| 1 | | pExTra-Lebron 115 | Lebron 115 replicate 1 | Non-toxic | ++ |
| 2 | | pExTra-Lebron 115 | Lebron 115 replicate 2 | Non-toxic | ++ |
| 3 | | pExTra-Lebron 115 | Lebron 115 replicate 3 | Non-toxic | ++ |

*Key: NG (no growth) - (no pink color) +(faint pink color) ++(obvious pink color) +++ (dark pink color)

#### Gene *116*; Score *0*

Images taken after 5 days at 37 ºC

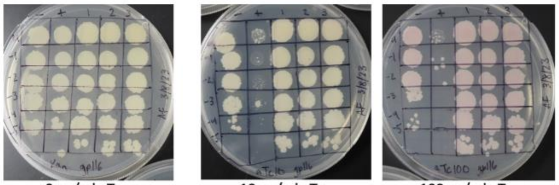


| Lane | Gene ID | Plasmid name | Gene name | Toxic/Non-toxic | Colony color on 100 ng/ml aTc plate* |
|---|---|---|---|---|---|
| + Toxic control | -- | pExTra02 | Fruitloop *52* | Toxic | - |
| - Non-toxic control | -- | pExTra03 | Fruitloop *52* mutant | Non-toxic | + |
| 1 | | pExTra-Lebron *116* | Lebron *116* replicate 1 | Non-toxic | + |
| 2 | | pExTra-Lebron *116* | Lebron *116* replicate 2 | Non-toxic | + |
| 3 | | pExTra-Lebron *116* | Lebron *116* replicate 3 | Non-toxic | + |

*Key: NG (no growth) - (no pink color) +(faint pink color) ++(obvious pink color) +++ (dark pink color)

#### Gene *117*; Score *1*

Images taken after 5 days at 37 ºC

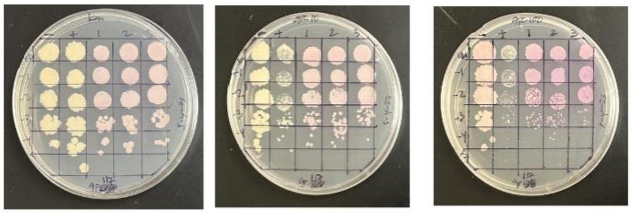


| Lane | Gene ID | Plasmid name | Gene name | Toxic/Non-toxic | Colony color on 100 ng/ml aTc plate* |
|---|---|---|---|---|---|
| - Non-toxic control | -- | pExTra03 | Fruitloop *52* mutant | Non-toxic | + |
| + Toxic control | -- | pExTra02 | Fruitloop *52* | Toxic | - |
| 1 | | pExTra-Lebron 117 | Lebron *117* replicate 1 | Toxic | ++ |
| 2 | | pExTra-Lebron 117 | Lebron *117* replicate 2 | Toxic | ++ |
| 3 | | pExTra-Lebron 117 | Lebron *117* replicate 3 | Toxic | ++ |

*Key: NG (no growth) - (no pink color) +(faint pink color) ++(obvious pink color) +++ (dark pink color)

#### Gene *118*; Score *0*

Images taken after 5 days at 37 ºC

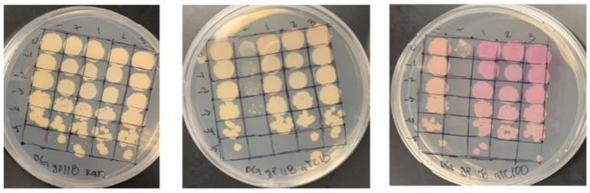


| Lane | Gene ID | Plasmid name | Gene name | Toxic/Non-toxic | Colony color on 100 ng/ml aTc plate* |
|---|---|---|---|---|---|
| + Toxic control | -- | pExTra02 | Fruitloop *52* | Toxic | - |
| - Non-toxic control | -- | pExTra03 | Fruitloop *52* mutant | Non-toxic | ++ |
| 1 | 131436 | pExtra-Lebron 118 | Lebron 118 replicate 1 | Non-toxic | ++ |
| 2 | 131436 | pExTra-Lebron 118 | Lebron 118 replicate 2 | Non-toxic | ++ |
| 3 | 131436 | pExTra-Lebron 118 | Lebron 118 replicate 3 | Non-toxic | +++ |

*Key: NG (no growth) - (no pink color) +(faint pink color) ++(obvious pink color) +++ (dark pink color)

#### Gene 119; Score *1*

Images taken after 5 days at 37 ºC

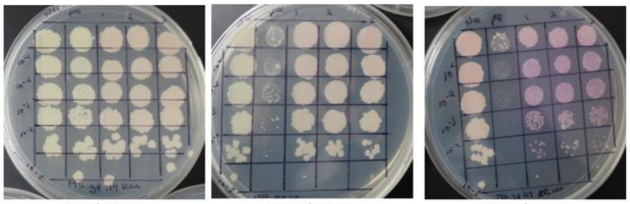


| Lane | Gene ID | Plasmid name | Gene name | Toxic/Non-toxic | Colony color on 100 ng/ml aTc plate* |
|---|---|---|---|---|---|
| + Toxic control | -- | pExTra02 | Fruitloop *52* | Toxic | - |
| - Non-toxic control | -- | pExTra03 | Fruitloop *52* mutant | Non-toxic | + |
| 1 | | pExTra-Lebron *119* | Lebron *119* replicate 1 | toxic | ++ |
| 2 | | pExTra-Lebron *119* | Lebron *119* replicate 2 | toxic | ++ |
| 3 | | pExTra-Lebron *119* | Lebron *119* replicate 3 | toxic | ++ |

*Key: NG (no growth) - (no pink color) +(faint pink color) ++(obvious pink color) +++ (dark pink color)

#### Gene *120*; Score *0*

Images taken after 5 days at 37 ºC

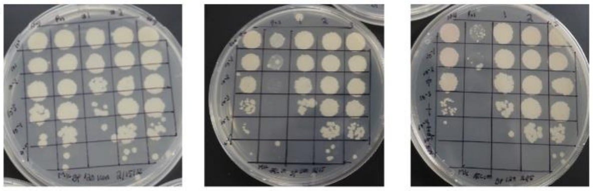


| Lane | Gene ID | Plasmid name | Gene name | Toxic/Non-toxic | Colony color on 100 ng/ml aTc plate* |
|---|---|---|---|---|---|
| + Toxic control | -- | pExTra02 | Fruitloop 52 | Toxic | - |
| - Non-toxic control | -- | pExTra03 | Fruitloop 52 mutant | Non-toxic | + |
| 1 | | pExTra-Lebron 120 | Lebron *120* replicate 1 | Non-toxic | - |
| 2 | | pExTra-Lebron 120 | Lebron *120* replicate 2 | Non-toxic | - |
| 3 | | pExTra-Lebron 120 | Lebron *120* replicate 3 | Non-toxic | - |

*Key: NG (no growth) - (no pink color) +(faint pink color) ++(obvious pink color) +++ (dark pink color)

#### Gene *121*; Score *0*

Images taken after 5 days at 37 ºC

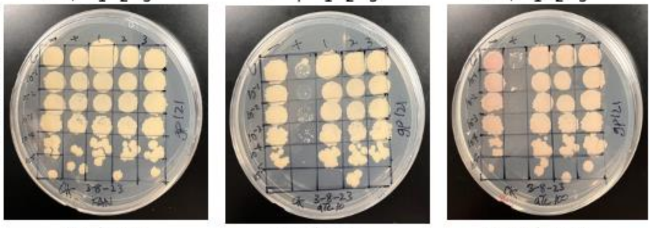


| Lane | Gene ID | Plasmid name | Gene name | Toxic/Non-toxic | Colony color on 100 ng/ml aTc plate* |
|---|---|---|---|---|---|
| - Non-toxic control | -- | pExTra03 | Fruitloop *52* mutant | Non-toxic | + |
| + Toxic control | -- | pExTra02 | Fruitloop *52* | Toxic | – |
| 1 | – | pExTra-Lebron 121 | Lebron *121* replicate 1 | Non-toxic | + |
| 2 | – | pExTra-Lebron 121 | Lebron *121* replicate 2 | Non-toxic | + |
| 3 | – | pExTra-Lebron 121 | Lebron *121* replicate 3 | Non-toxic | + |

*Key: NG (no growth) - (no pink color) +(faint pink color) ++(obvious pink color) +++ (dark pink color)

#### Gene *122*; Score *0*

Images taken after 5 days at 37 ºC

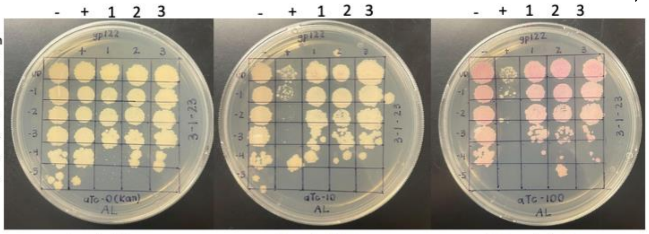


| Lane | Gene ID | Plasmid name | Gene name | Toxic/Non-toxic | Colony color on 100 ng/ml aTc plate* |
|---|---|---|---|---|---|
| + Toxic control | -- | pExTra02 | Fruitloop *52* | Toxic | - |
| - Non-toxic control | -- | pExTra03 | Fruitloop *52* mutant | Non-toxic | ++ |
| 1 | | pExTra-Lebron122 | Lebron 122 replicate 1 | Non-toxic | ++ |
| 2 | | pExTra-Lebron122 | Lebron 122 replicate 2 | Non-toxic | ++ |
| 3 | | pExTra-Lebron122 | Lebron 122 replicate 3 | Non-toxic | ++ |

*Key: NG (no growth) - (no pink color) +(faint pink color) ++(obvious pink color) +++ (dark pink color)

#### Gene 123: Score *0*

Images taken after 5 days at 37 ºC

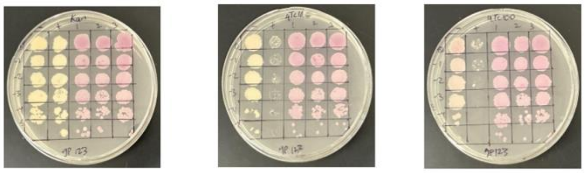


| Lane | Gene ID | Plasmid name | Gene name | Toxic/Non-toxic | Colony color on 100 ng/ml aTc plate* |
|---|---|---|---|---|---|
| - Non-toxic control | -- | pExTra03 | Fruitloop *52* mutant | Non-toxic | + |
| + Toxic control | -- | pExTra02 | Fruitloop *52* | Toxic | - |
| 1 | | pExTra-Lebron 123 | Lebron *123* replicate 1 | Non-toxic | ++ |
| 2 | | pExTra-Lebron 123 | Lebron *123* replicate 2 | Non-toxic | ++ |
| 3 | | pExTra-Lebron 123 | Lebron *123* replicate 3 | Non-toxic | ++ |

*Key: NG (no growth) - (no pink color) +(faint pink color) ++(obvious pink color) +++ (dark pink color)

#### Gene *124*; Score *0*

Images taken after 5 days at 37 ºC

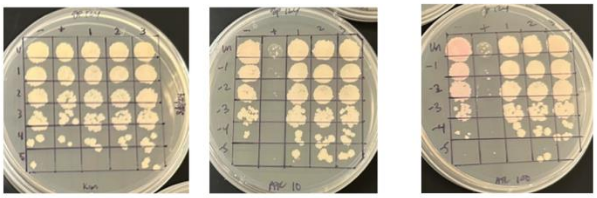


| Lane | Gene ID | Plasmid name | Gene name | Toxic/Non-toxic | Colony color on 100 ng/ml aTc plate* |
|---|---|---|---|---|---|
| - Non-toxic control | -- | pExTra03 | Fruitloop *52* mutant | Non-toxic | ++ |
| + Toxic control | -- | pExTra02 | Fruitloop *52* | Toxic | - |
| 1 | | pExTra-Lebron 124 | Lebron *124* replicate 1 | Non-toxic | + |
| 2 | | pExTra-Lebron 124 | Lebron *124* replicate 2 | Non-toxic | + |
| 3 | | pExTra-Lebron 124 | Lebron *124* replicate 3 | Non-toxic | + |

*Key: NG (no growth) - (no pink color) +(faint pink color) ++(obvious pink color) +++ (dark pink color)

#### Gene *125*; Score *1*

Images taken after 5 days at 37 ºC

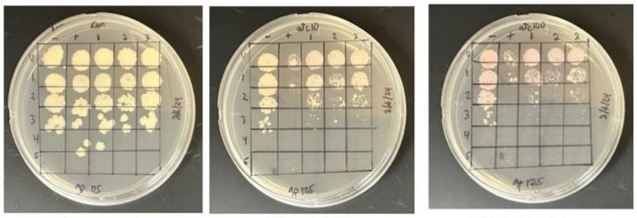


| Lane | Gene ID | Plasmid name | Gene name | Toxic/Non-toxic | Colony color on 100 ng/ml aTc plate* |
|---|---|---|---|---|---|
| - Non-toxic control | -- | pExTra03 | Fruitloop *52* mutant | Non-toxic | + |
| + Toxic control | -- | pExTra02 | Fruitloop *52* | Toxic | - |
| 1 | | pExTra-Lebron 125 | Lebron *125* replicate 1 | Toxic | + |
| 2 | | pExTra-Lebron 125 | Lebron *125* replicate 2 | Toxic | + |
| 3 | | pExTra-Lebron 125 | Lebron *125* replicate 3 | Toxic | + |

*Key: NG (no growth) - (no pink color) +(faint pink color) ++(obvious pink color) +++ (dark pink color)

#### Gene *126*; Score *0*

Images taken after 5 days at 37 ºC

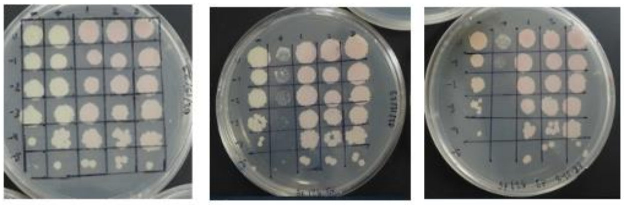


| Lane | Gene ID | Plasmid name | Gene name | Toxic/Non-toxic | Colony color on 100 ng/ml aTc plate* |
|---|---|---|---|---|---|
| - Non-toxic control | -- | pExTra03 | Fruitloop *52* mutant | Non-toxic | + |
| + Toxic control | -- | pExTra02 | Fruitloop *52* | Toxic | - |
| 1 | – | pExTra-Lebron 126 | Lebron *126* replicate 1 | Non-toxic | + |
| 2 | – | pExTra-Lebron 126 | Lebron *126* replicate 2 | Non-toxic | + |
| 3 | – | pExTra-Lebron 126 | Lebron *126* replicate 3 | Non-toxic | + |

*Key: NG (no growth) - (no pink color) +(faint pink color) ++(obvious pink color) +++ (dark pink color)

#### Gene 127; Score *0*

Images taken after 5 days at 37 ºC

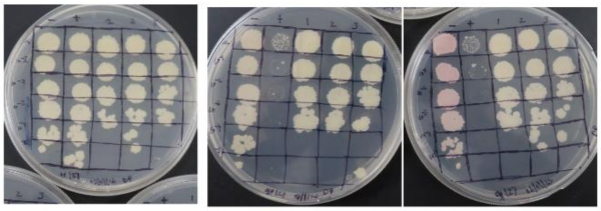


| Lane | Gene ID | Plasmid name | Gene name | Toxic/Non-toxic | Colony color on 100 ng/ml aTc plate* |
|---|---|---|---|---|---|
| + Toxic control | -- | pExTra02 | Fruitloop *52* | Toxic | - |
| - Non-toxic control | -- | pExTra03 | Fruitloop *52* mutant | Non-toxic | ++ |
| 1 | -- | pExTra-Lebron 127 | Lebron *127* replicate 1 | Non-toxic | - |
| 2 | -- | pExTra-Lebron 127 | Lebron *127* replicate 2 | Non-toxic | - |
| 3 | -- | pExTra-Lebron 127 | Lebron *127* replicate 3 | Non-toxic | - |

*Key: NG (no growth) - (no pink color) +(faint pink color) ++(obvious pink color) +++ (dark pink color)

#### Gene *128*; Score *3*

Images taken after 5 days at 37 ºC

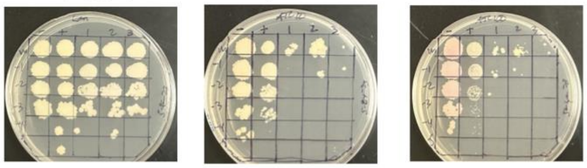


| Lane | Gene ID | Plasmid name | Gene name | Toxic/Non-toxic | Colony color on 100 ng/ml aTc plate* |
|---|---|---|---|---|---|
| - Non-toxic control | -- | pExTra03 | Fruitloop *52* mutant | Non-toxic | + |
| + Toxic control | -- | pExTra02 | Fruitloop *52* | Toxic | - |
| 1 | | pExTra-Lebron 128 | Lebron *128* replicate 1 | Toxic | - |
| 2 | | pExTra-Lebron 128 | Lebron *128* replicate 2 | Toxic | - |
| 3 | | pExTra-Lebron 128 | Lebron *128* replicate 3 | Toxic | - |

*Key: NG (no growth) - (no pink color) +(faint pink color) ++(obvious pink color) +++ (dark pink color)

#### Gene *129*; Score *0*

Images taken after 5 days at 37 ºC

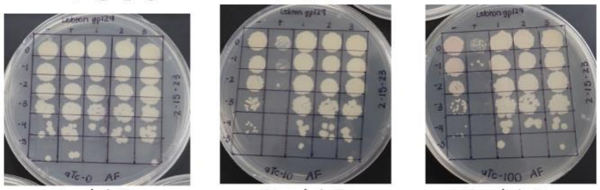


| Lane | Gene ID | Plasmid name | Gene name | Toxic/Non-toxic | Colony color on 100 ng/ml aTc plate* |
|---|---|---|---|---|---|
| + Toxic control | -- | pExTra02 | Fruitloop *52* | Toxic | - |
| - Non-toxic control | -- | pExTra03 | Fruitloop *52* mutant | Non-toxic | + |
| 1 | 131436 | pExTra-Lebron *129* | Lebron *129* replicate 1 | Non-toxic | - |
| 2 | 131436 | pExTra-Lebron *129* | Lebron *129* replicate 2 | Non-toxic | - |
| 3 | 131436 | pExTra-Lebron *129* | Lebron *129* replicate 3 | Non-toxic | - |

*Key: NG (no growth) - (no pink color) +(faint pink color) ++(obvious pink color) +++ (dark pink color)

#### Gene *130*; Score *0*

Images taken after 5 days at 37 ºC

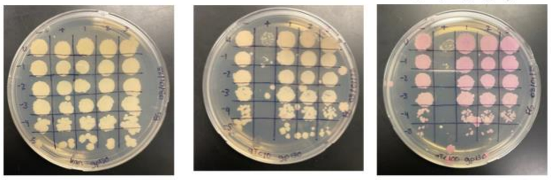


| Lane | Gene ID | Plasmid name | Gene name | Toxic/Non-toxic | Colony color on 100 ng/ml aTc plate* |
|---|---|---|---|---|---|
| + Toxic control | -- | pExTra02 | Fruitloop *52* | Toxic | - |
| - Non-toxic control | -- | pExTra03 | Fruitloop *52* mutant | Non-toxic | ++ |
| 1 | | pExTra-Lebron 130 | Lebron 130 replicate 1 | Non-Toxic | +++ |
| 2 | | pExTra-Lebron 130 | Lebron 130 replicate 2 | Non-Toxic | +++ |
| 3 | | pExTra-Lebron 130 | Lebron *130* replicate 3 | Non-Toxic | +++ |

*Key: NG (no growth) - (no pink color) +(faint pink color) ++(obvious pink color) +++ (dark pink color)

Gene 131: Score *1*

Images taken after 5 days at 37 ºC

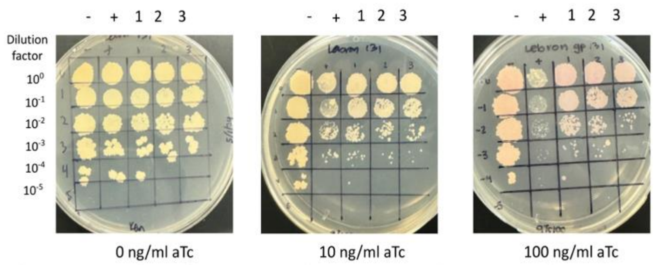


| Lane | Gene ID | Plasmid name | Gene name | Toxic/Non-toxic | Colony color on 100 ng/ml aTc plate* |
|---|---|---|---|---|---|
| - Non-toxic control | -- | pExTra03 | Fruitloop *52* mutant | Non-toxic | + |
| + Toxic control | -- | pExTra02 | Fruitloop *52* | Toxic | - |
| 1 | | pExTra-Lebron 131 | Lebron *131* replicate 1 | Toxic | ++ |
| 2 | | pExTra-Lebron 131 | Lebron *131* replicate 2 | Toxic | ++ |
| 3 | | pExTra-Lebron 131 | Lebron *131* replicate 3 | Toxic | ++ |

*Key: NG (no growth) - (no pink color) +(faint pink color) ++(obvious pink color) +++ (dark pink color)

Gene 132: Score *0*

Images taken after 5 days at 37 ºC

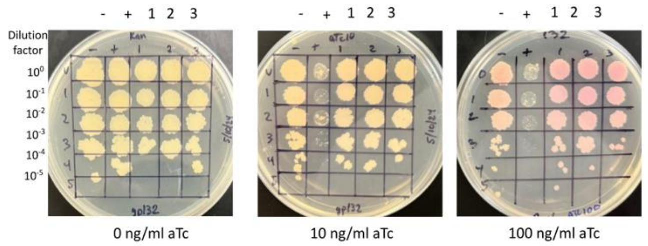


| Lane | Gene ID | Plasmid name | Gene name | Toxic/Non-toxic | Colony color on 100 ng/ml aTc plate* |
|---|---|---|---|---|---|
| - Non-toxic control | -- | pExTra03 | Fruitloop *52* mutant | Non-toxic | + |
| + Toxic control | -- | pExTra02 | Fruitloop *52* | Toxic | - |
| 1 | | pExTra-Lebron 132 | Lebron *132* replicate 1 | Non-toxic | +++ |
| 2 | | pExTra-Lebron 132 | Lebron *132* replicate 2 | Non-toxic | +++ |
| 3 | | pExTra-Lebron 132 | Lebron *132* replicate 3 | Non-toxic | +++ |

*Key: NG (no growth) - (no pink color) +(faint pink color) ++(obvious pink color) +++ (dark pink color)

**Supplemental Figure 1**. Shown are the results of representative cytotoxicity assays for 122 LeBron genes screened in this study. Each strain was spotted in triplicate alongside *M. smegmatis*/pExtra-Fruitloop 52 (+) and pExtra-Fruitloop 52I170S (-) control strains in 7H10 Kan supplemented with 0, 10, or 100 ng/ml aTc. In all experiments, 100 to 10-5 dilutions are shown. Plates were monitored over 5 days at 37C, with results shown to be illustrate effects on colony color and size. Colony color was scored using the indicated key shown at the bottom of each data card.

**Supplemental Table 1: Primer sequences used in this study to amplify genes**

| Primer Name | Primer Sequence (5' to 3') |
|---|---|
| oLebron1_F | atgcggaggaatcacttccatATGGGTAGAGGACGTGGTAAGG |
| oLebron1_R | tgcaggatccgactcgagtgtcgacTCACTTGCCGTACTTCTCAAG |
| oLebron2_F | atgcggaggaatcacttccatATGGGGGATGGCGTTAAAAAC |
| oLebron2_R | tgcaggatccgactcgagtgtcgacTCAGGACGCCTGAGCG |
| oLebron3_F | atgcggaggaatcacttccatATGCAGTCCATTCGTTCGG |
| oLebron3_R | tgcaggatccgactcgagtgtcgacTCAATCATCACCATTACCAATCTC |
| oLebron4_F | atgcggaggaatcacttccatATGACGGTTATTCCGTCAATTCCC |
| oLebron4_R | tgcaggatccgactcgagtgtcgacTCAATAGCCATTAGTCGACCCC |
| oLebron5_F | atgcggaggaatcacttccatATGGCTATTGAGCTACCCG |
| oLebron5_R | tgcaggatccgactcgagtgtcgacTCAGGCAGCCTTACTGGC |
| oLebron6_F | atgcggaggaatcacttccatATGGCTGAGGCTGACGAG |
| oLebron6_R | tgcaggatccgactcgagtgtcgacTCAAGCTGAAGCGCTTAGTTTCC |
| oLebron7_F | atgcggaggaatcacttccatATGACTGGCTTAAGTGAAGGAAAAT |
| oLebron7_R | tgcaggatccgactcgagtgtcgacTCAGACGCCGCCGCG |
| oLebron8_F | atgcggaggaatcacttccatATGGCGCACATTTTTGTGAAGC |
| oLebron8_R | tgcaggatccgactcgagtgtcgacTCAGGCAGGGACGGTG |
| oLebron9_F | atgcggaggaatcacttccatATGCAGACCCTCATGTCG |
| oLebron9_R | tgcaggatccgactcgagtgtcgacTCACCAGGGCACCACC |
| oLebron10_F | atgcggaggaatcacttccatATGAACGAAGAGACCATAACGG |
| oLebron10_R | tgcaggatccgactcgagtgtcgacTCACCCCGTCACCGCC |
| oLebron11_F | atgcggaggaatcacttccatATGCCTGCTGCGGGTAC |
| oLebron11_R | tgcaggatccgactcgagtgtcgacTCACAGCCCATTCTTAGCC |
| oLebron12_F | atgcggaggaatcacttccatATGGCTCTGGCGCTGC |
| oLebron12_R | tgcaggatccgactcgagtgtcgacTCAAGAACGAAACAGCTCATAAATG |
| oLebron13_F | atgcggaggaatcacttccatATGGCTGATTTCTACACGATTAAGG |
| oLebron13_R | tgcaggatccgactcgagtgtcgacTCACGCCGAGACGGTC |
| oLebron14_F | atgcggaggaatcacttccatATGGCTGAAAAGAGCAAGG |
| oLebron14_R | tgcaggatccgactcgagtgtcgacTCACTTTCCCGAGTCCTTATCC |
| oLebron15_F | atgcggaggaatcacttccatATGGCTGAAAAGAGCAAGG |
| oLebron15_R | tgcaggatccgactcgagtgtcgacTCATAGGGTGACCCATTTCG |
| oLebron16_F | atgcggaggaatcacttccatATGGCTGAATACGTTGCC |
| oLebron16_R | tgcaggatccgactcgagtgtcgacTCATCGCATTGGCAGGATGG |
| oLebron17_F | atgcggaggaatcacttccatATGGAAAGACAACTGAATATGTCCG |
| oLebron17_R | tgcaggatccgactcgagtgtcgacTCAGCGCAGCACCCCC |
| oLebron18_F | atgcggaggaatcacttccatATGGCTGACACGCAGTATTGG |
| oLebron18_R | tgcaggatccgactcgagtgtcgacTCAACTCTGGATAATGTGGACACC |
| oLebron19_F | atgcggaggaatcacttccatATGGCTAAAACGCAGTCGG |

| | |
|---|---|
| oLebron19_R | tgcaggatccgactcgagtgtcgacTCAGGTCAACCTCCCCTCC |
| oLebron20_F | atgcggaggaatcacttccatATGACGATGCCTAACGGGG |
| oLebron20_R | tgcaggatccgactcgagtgtcgacTCAATTACCAGACAAATCCATCGCC |
| oLebron21_F | atgcggaggaatcacttccatATGATGACGTGGGCGG |
| oLebron21_R | tgcaggatccgactcgagtgtcgacTCAAGAATTGCCCCATTGAAAGG |
| oLebron22_F | atgcggaggaatcacttccatATGTTGGATCACGCGTTCAGG |
| oLebron22_R | tgcaggatccgactcgagtgtcgacTCAAGCCGCAATCGGC |
| oLebron23_F | atgcggaggaatcacttccatATGAACTATCAGGCTGGCC |
| oLebron23_R | tgcaggatccgactcgagtgtcgacTCAAGCATTGAACCACCCC |
| oLebron24_F | atgcggaggaatcacttccatATGCTTAACATCGGCCCG |
| oLebron24_R | tgcaggatccgactcgagtgtcgacTCAAACACCGTCCTTCGC |
| oLebron25_F | atgcggaggaatcacttccatATGGCGCTACTGGGAAGC |
| oLebron25_R | tgcaggatccgactcgagtgtcgacTCAGCTCGCCGCTTTCAG |
| oLebron26_F | atgcggaggaatcacttccatATGGCGAAGGGTCTCAATG |
| oLebron26_R | tgcaggatccgactcgagtgtcgacTCAAACAACACTCCTTAGGTAATC |
| oLebron27_F | atgcggaggaatcacttccatATGTTGGTTTACGACGAGAAG |
| oLebron27_R | tgcaggatccgactcgagtgtcgacTCACTCACCGTCCTTACC |
| oLebron28_F | atgcggaggaatcacttccatATGAAGATTCTCGGTCACAAGC |
| oLebron28_R | tgcaggatccgactcgagtgtcgacTCATCCCTTCTCAATAAAGGGCG |
| oLebron29_F | atgcggaggaatcacttccatATGGACCCGAGCGATATGG |
| oLebron29_R | tgcaggatccgactcgagtgtcgacTCACTTCCAAATGTCCAATTCC |
| oLebron30_F | atgcggaggaatcacttccatATGGAAGTAAAAGTTTATTCGCCG |
| oLebron30_R | tgcaggatccgactcgagtgtcgacTCAGGCCGCTGTCTGG |
| oLebron31_F | atgcggaggaatcacttccatATGATTGAGGCTGTGGTGC |
| oLebron31_R | tgcaggatccgactcgagtgtcgacTCAGAGTTTGGCGGGTGG |
| oLebron32_F | atgcggaggaatcacttccatATGCAGCCGCTGATCATTGG |
| oLebron32_R | tgcaggatccgactcgagtgtcgacTCATGCCGCCAGTTTATCTC |
| oLebron33_F | atgcggaggaatcacttccatATGGGTAAGCGCGTAACC |
| oLebron33_R | tgcaggatccgactcgagtgtcgacTCAGGACTGCGCTGCG |
| oLebron34_F | atgcggaggaatcacttccatATGAGAATCCGTTGGCC |
| oLebron34_R | tgcaggatccgactcgagtgtcgacTCACCCATCGCTTTCGTATCC |
| oLebron35_F | atgcggaggaatcacttccatATGCCTGGTCAGAGCGG |
| oLebron35_R | tgcaggatccgactcgagtgtcgacTCACTCAACCCCCTGC |
| oLebron36_F | atgcggaggaatcacttccatATGGCCGAGACGGACTC |
| oLebron36_R | tgcaggatccgactcgagtgtcgacTCAAACACTGGGTAACAGAGC |
| oLebron37_F | atgcggaggaatcacttccatATGCATACGACGCACTCTG |
| oLebron37_R | tgcaggatccgactcgagtgtcgacTCAGTGCGCGGTCTTGTG |
| oLebron38_F | atgcggaggaatcacttccatATGCGCATGTCCCAAAAATCG |
| oLebron38_R | tgcaggatccgactcgagtgtcgacTCACAGGGGAGGAATATCTG |

| | |
|---|---|
| oLebron39_F | atgcggaggaatcacttccatATGCCTTCCGAACTTTGTCG |
| oLebron39_R | tgcaggatccgactcgagtgtcgacTCACACGGCCCTCCGG |
| oLebron40_F | atgcggaggaatcacttccatATGACCGCGCCCACAG |
| oLebron40_R | tgcaggatccgactcgagtgtcgacTCACAGGTCTGCCTCC |
| oLebron41_F | atgcggaggaatcacttccatATGACCGCCGCGCAAG |
| oLebron41_R | tgcaggatccgactcgagtgtcgacTCACCCCTGATTGCCG |
| oLebron42_F | atgcggaggaatcacttccatATGTGCGACCCGGCAATC |
| oLebron42_R | tgcaggatccgactcgagtgtcgacTCATACCGTCCAGCCC |
| oLebron43_F | atgcggaggaatcacttccatATGAGTACAGCGGGTGTAAAAAATC |
| oLebron43_R | tgcaggatccgactcgagtgtcgacTCAGATGGGAGTCGCG |
| oLebron44_F | atgcggaggaatcacttccatATGCGCGGTTATCGGG |
| oLebron44_R | tgcaggatccgactcgagtgtcgacTCAGCTCTCCCTTCCC |
| oLebron45_F | atgcggaggaatcacttccatATGACCGTTATCTATTTGGGC |
| oLebron45_R | tgcaggatccgactcgagtgtcgacTCAGAGGTAGTCGTCGC |
| oLebron46_F | atgcggaggaatcacttccatATGGACAAAGACTGGCTGTTTTCAG |
| oLebron46_R | tgcaggatccgactcgagtgtcgacTCAGCCTTCATCGGGC |
| oLebron47_F | atgcggaggaatcacttccatATGAAGGCTAGCCAGCGG |
| oLebron47_R | tgcaggatccgactcgagtgtcgacTCAGCCCGCTTGAGGC |
| oLebron48_F | atgcggaggaatcacttccatATGCGAAACGCCGTATTGG |
| oLebron48_R | tgcaggatccgactcgagtgtcgacTCAGGGGAGGCTAGGAG |
| oLebron49_F | atgcggaggaatcacttccatATGGAAGGAACCATTTTCTTGAACC |
| oLebron49_R | tgcaggatccgactcgagtgtcgacTCAGCGAGCCGGGGCAC |
| oLebron50_F | atgcggaggaatcacttccatATGTCGCAAAAGAGCGTGTTCG |
| oLebron50_R | tgcaggatccgactcgagtgtcgacTCATCTCTTAGCCCCCACTCC |
| oLebron51_F | atgcggaggaatcacttccatATGATCGGTTCTGTTTGGCG |
| oLebron51_R | tgcaggatccgactcgagtgtcgacTCAAAAGCTGGCACCGTAAACC |
| oLebron52_F | atgcggaggaatcacttccatATGCCAGCTTTTGAGGGTG |
| oLebron52_R | tgcaggatccgactcgagtgtcgacTCATTTCCCGACGTAAAGG |
| oLebron53_F | atgcggaggaatcacttccatATGACGACATTCCTTGACAGTC |
| oLebron53_R | tgcaggatccgactcgagtgtcgacTCACAACGTGATCTGCGAGTACC |
| oLebron54_F | atgcggaggaatcacttccatATGGCTAAGACGGTTCGAGAG |
| oLebron54_R | tgcaggatccgactcgagtgtcgacTCACCGGCGCTCCACG |
| oLebron55_F | atgcggaggaatcacttccatATGGTGGTCCTGCTCC |
| oLebron55_R | tgcaggatccgactcgagtgtcgacTCAAAGCTCCCATCGAAAGC |
| oLebron56_F | atgcggaggaatcacttccatATGAGCGGCTACTACGAGG |
| oLebron56_R | tgcaggatccgactcgagtgtcgacTCATTCATCCCTGTATTCCTTAACG |
| oLebron57_F | atgcggaggaatcacttccatATGAATAAGCCGGTCGGC |
| oLebron57_R | tgcaggatccgactcgagtgtcgacTCATCCTTCAATGATCACAGTG |
| oLebron58_F | atgcggaggaatcacttccatATGGTTTACCCACCGGACTTTACC |

| | |
|---|---|
| oLebron58_R | tgcaggatccgactcgagtgtcgacTCACCAGTGATAACGGCC |
| oLebron59_F | atgcggaggaatcacttccatATGGCCGTTATCACTGGTAATCG |
| oLebron59_R | tgcaggatccgactcgagtgtcgacTCACTTAGCTCCGTTCTTAGCTTTC |
| oLebron60_F | atgcggaggaatcacttccatATGACAGCCACGAAAGCTG |
| oLebron60_R | tgcaggatccgactcgagtgtcgacTCAGAGAGTCACCATCCCC |
| oLebron61_F | atgcggaggaatcacttccatATGCCGCCTCCTGTTAAGG |
| oLebron61_R | tgcaggatccgactcgagtgtcgacTCATGTGAGACCTTGCCG |
| oLebron62_F | atgcggaggaatcacttccatATGCGTGGCATGTCTATTCC |
| oLebron62_R | tgcaggatccgactcgagtgtcgacTCAACCGGCTAGCGCAAAG |
| oLebron63_F | atgcggaggaatcacttccatATGAGCGATACCTACGAGTACAACC |
| oLebron63_R | tgcaggatccgactcgagtgtcgacTCACCATTCACCCCTGC |
| oLebron64_F | atgcggaggaatcacttccatATGGTGAAAAAGATATTTGCGGC |
| oLebron64_R | tgcaggatccgactcgagtgtcgacTCATGGCTCTTCCGGTTCATC |
| oLebron65_F | atgcggaggaatcacttccatATGTGGACCCCCGATAAC |
| oLebron65_R | tgcaggatccgactcgagtgtcgacTCACTGCACTGGCCCAC |
| oLebron66_F | atgcggaggaatcacttccatATGAGCGACTTCGGGAAAATCC |
| oLebron66_R | tgcaggatccgactcgagtgtcgacTCAAGCCGCTACCCTTTCC |
| oLebron67_F | atgcggaggaatcacttccatATGACGGAGGCTCAGC |
| oLebron67_R | tgcaggatccgactcgagtgtcgacTCAAGTGATTCGCTGGTAGTGC |
| oLebron68_F | atgcggaggaatcacttccatATGGAGGGCAAGCCGG |
| oLebron68_R | tgcaggatccgactcgagtgtcgacTCATGCTGAATTGTCACGCG |
| oLebron69_F | atgcggaggaatcacttccatATGGAGATGGCTAAAGCCAAGG |
| oLebron69_R | tgcaggatccgactcgagtgtcgacTCACCATTTGTCCCCCAACC |
| oLebron70_F | atgcggaggaatcacttccatATGGTAGATCCCGATGAGGAC |
| oLebron70_R | tgcaggatccgactcgagtgtcgacTCAGATAAACGGCCATGAATTCGG |
| oLebron71_F | atgcggaggaatcacttccatATGGCCGTTTATCTGAGGGC |
| oLebron71_R | tgcaggatccgactcgagtgtcgacTCAGTGCTCACTTAACCCCTTAACC |
| oLebron72_F | atgcggaggaatcacttccatATGAGCACTAAGTTCGCGCC |
| oLebron72_R | tgcaggatccgactcgagtgtcgacTCATTGGCTACTCGCCAAAAC |
| oLebron73_F | atgcggaggaatcacttccatATGGCAACCACGCCGAAG |
| oLebron73_R | tgcaggatccgactcgagtgtcgacTCACTGGATACCGGCTTTACC |
| oLebron74_F | atgcggaggaatcacttccatATGAGCGAGCTGTCGTTAAC |
| oLebron74_R | tgcaggatccgactcgagtgtcgacTCAGAAATCTCCCGGCTGAAC |
| oLebron75_F | atgcggaggaatcacttccatATGAAACTGGGAAACGTCGTG |
| oLebron75_R | tgcaggatccgactcgagtgtcgacTCACTTGAACTCGATAGACACCTGC |
| oLebron76_F | atgcggaggaatcacttccatATGGAGCACCTACTTGTAACG |
| oLebron76_R | tgcaggatccgactcgagtgtcgacTCAGCCAGCTTGCGCC |
| oLebron77_F | atgcggaggaatcacttccatATGTTTTTCCCCCTAAAGGGC |
| oLebron77_R | tgcaggatccgactcgagtgtcgacTCACGCAACCTCCATCAAATCG |

| | |
|---|---|
| oLebron78_F | atgcggaggaatcacttccatATGGCAGAACACGTGACGG |
| oLebron78_R | tgcaggatccgactcgagtgtcgacTCAAGCAGCATGTCGCTC |
| oLebron79_F | atgcggaggaatcacttccatATGCTGCTTAGCACCAAAATTAAGC |
| oLebron79_R | tgcaggatccgactcgagtgtcgacTCACGCATACCGTGGCAAC |
| oLebron80_F | atgcggaggaatcacttccatATGGGACTAGGGGAGC |
| oLebron80_R | tgcaggatccgactcgagtgtcgacTCAGCCACGCTTAACCTTCTC |
| oLebron81_F | atgcggaggaatcacttccatATGGCTAGTGTTGACATTCAG |
| oLebron81_R | tgcaggatccgactcgagtgtcgacTCAGAACGGAGGCTCTTCC |
| oLebron82_F | atgcggaggaatcacttccatATGCTGCGCAAAATCGG |
| oLebron82_R | tgcaggatccgactcgagtgtcgacTCATTCCCGCCACCAG |
| oLebron83_F | atgcggaggaatcacttccatATGATCGGGTTCCTGATTTTCC |
| oLebron83_R | tgcaggatccgactcgagtgtcgacTCATTCTTCGTCGTCCTCCTTAGC |
| oLebron84_F | atgcggaggaatcacttccatATGAAGAGAATCCTAGTAACCGG |
| oLebron84_R | tgcaggatccgactcgagtgtcgacTCAGTTGTCTCCGTAGTTGATTACG |
| oLebron85_F | atgcggaggaatcacttccatATGAGTTTTGACGCGCC |
| oLebron85_R | tgcaggatccgactcgagtgtcgacTCAAGCCGCTACCTCCATATCC |
| oLebron86_F | atgcggaggaatcacttccatATGAGTTACGACTACAAGGCTCC |
| oLebron86_R | tgcaggatccgactcgagtgtcgacTCAGCCGAGCACCACC |
| oLebron87_F | atgcggaggaatcacttccatATGGTCAATTACCTGTCGGG |
| oLebron87_R | tgcaggatccgactcgagtgtcgacTCATTCCTCTGCCCTCC |
| oLebron88_F | atgcggaggaatcacttccatATGAGTTACACGCTGTTCGC |
| oLebron88_R | tgcaggatccgactcgagtgtcgacTCAGGCGCGGACATGG |
| oLebron89_F | atgcggaggaatcacttccatATGTCCGCGCCTAAGTGG |
| oLebron89_R | tgcaggatccgactcgagtgtcgacTCACAAGTTCCCTTTGGTTGC |
| oLebron90_F | atgcggaggaatcacttccatATGTGGGTTGAGAGTCC |
| oLebron90_R | tgcaggatccgactcgagtgtcgacTCAAAGCACTTCCGGAAGC |
| oLebron91_F | atgcggaggaatcacttccatATGAGGCCGTCAGATGTTGC |
| oLebron91_R | tgcaggatccgactcgagtgtcgacTCAATTCTTTCCCCCTAAGC |
| oLebron92_F | atgcggaggaatcacttccatATGACCACGAGGGATGC |
| oLebron92_R | tgcaggatccgactcgagtgtcgacTCACGCGAACGCCCTC |
| oLebron93_F | atgcggaggaatcacttccatATGAGCGCGCGGGTCC |
| oLebron93_R | tgcaggatccgactcgagtgtcgacTCACACCGGCCTCAAC |
| oLebron94_F | atgcggaggaatcacttccatATGAGTTACATAAGCTTGGATCG |
| oLebron94_R | tgcaggatccgactcgagtgtcgacTCACCAAGCCTTCCCG |
| oLebron95_F | atgcggaggaatcacttccatATGATGGACGATCCTTACC |
| oLebron95_R | tgcaggatccgactcgagtgtcgacTCACTCATCGAGGTCAACC |
| oLebron96_F | atgcggaggaatcacttccatATGGAGACGACGGAGG |
| oLebron96_R | tgcaggatccgactcgagtgtcgacTCAAGCCGCTTGCGCAG |
| oLebron97_F | atgcggaggaatcacttccatATGCCGGGCAATCTGC |

| | |
|---|---|
| oLebron97_R | tgcaggatccgactcgagtgtcgacTCATGCGAACCAATCTCTCG |
| oLebron98_F | atgcggaggaatcacttccatATGTCGGATACCGAGAATCAGC |
| oLebron98_R | tgcaggatccgactcgagtgtcgacTCAATCATCAGCCGCCAAAATAGC |
| oLebron99_F | atgcggaggaatcacttccatATGATTGAGGTTTACAACTCGG |
| oLebron99_R | tgcaggatccgactcgagtgtcgacTCAGACGCACTCGGCG |
| oLebron100_F | atgcggaggaatcacttccatATGGAGTTCAAAGCTAATTGTTGC |
| oLebron100_R | tgcaggatccgactcgagtgtcgacTCAGCCCCTCTTGTCCTG |
| oLebron101_F | atgcggaggaatcacttccatATGGAGTTCGTGATGGAGTGC |
| oLebron101_R | tgcaggatccgactcgagtgtcgacTCACAGGACCATTGCAAAGAC |
| oLebron104_F | atgcggaggaatcacttccatATGACTGATGTTGATCCGATCTGG |
| oLebron104_R | tgcaggatccgactcgagtgtcgacTCAACTCTCATCAACCTCAGG |
| oLebron110_F | atgcggaggaatcacttccatATGTCTGAGGTTAAGAGCCTTTACG |
| oLebron110_R | tgcaggatccgactcgagtgtcgacTCACCCCGCCCCATTAG |
| oLebron111_F | atgcggaggaatcacttccatATGACGTTGTATCACCGGACG |
| oLebron111_R | tgcaggatccgactcgagtgtcgacTCACTTACGCTTCTTACGCGG |
| oLebron113_F | atgcggaggaatcacttccatATGCCCGCTTCAACGG |
| oLebron113_R | tgcaggatccgactcgagtgtcgacTCAGTCCGGTTCGATTCC |
| oLebron115_F | atgcggaggaatcacttccatATGACCTGTCAAATCCCGTGC |
| oLebron115_R | tgcaggatccgactcgagtgtcgacTCAGCAGTGCCCGCGTG |
| oLebron116_F | atgcggaggaatcacttccatATGGACCGGGTTGTGGTC |
| oLebron116_R | tgcaggatccgactcgagtgtcgacTCACTTGAGTGTGACGGTGG |
| oLebron117_F | atgcggaggaatcacttccatATGTACGGTAGAGGCATGAGC |
| oLebron117_R | tgcaggatccgactcgagtgtcgacTCAAGCGTCCCTGATGAGC |
| oLebron118_F | atgcggaggaatcacttccatATGCAGAGCGACAAATGCG |
| oLebron118_R | tgcaggatccgactcgagtgtcgacTCAACCCTGGTTTTTGCG |
| oLebron119_F | atgcggaggaatcacttccatATGGACTGCCTAGAAGGTCC |
| oLebron119_R | tgcaggatccgactcgagtgtcgacTCAGTCGTCCTCGTTCC |
| oLebron120_F | atgcggaggaatcacttccatATGCGCAAATTCATCAGCG |
| oLebron120_R | tgcaggatccgactcgagtgtcgacTCAGTTGGTGAGGCCG |
| oLebron121_F | atgcggaggaatcacttccatATGGACATGAGCAACGAAACG |
| oLebron121_R | tgcaggatccgactcgagtgtcgacTCAGCCCCCCATCTGC |
| oLebron122_F | atgcggaggaatcacttccatATGGTTTCTGACCCGACG |
| oLebron122_R | tgcaggatccgactcgagtgtcgacTCAAATCCGATCTTCAATGGAG |
| oLebron123_F | atgcggaggaatcacttccatATGCTGCCAAACATCCCAG |
| oLebron123_R | tgcaggatccgactcgagtgtcgacTCAGAAACCATGCTCGGAGAG |
| oLebron124_F | atgcggaggaatcacttccatATGGCCGCGTTCGCAAAG |
| oLebron124_R | tgcaggatccgactcgagtgtcgacTCAGCTGACGGCTTCTGC |
| oLebron125_F | atgcggaggaatcacttccatATGAAGCGCACACCGAAATG |
| oLebron125_R | tgcaggatccgactcgagtgtcgacTCATGCTGCTTCTCCTTCTGC |

| | |
|---|---|
| oLebron126_F | atgcggaggaatcacttccatATGGGCCGAGGTCGCG |
| oLebron126_R | tgcaggatccgactcgagtgtcgacTCACTTCTTAGCCCGGTC |
| oLebron127_F | atgcggaggaatcacttccatATGCGCGTAGGCACAC |
| oLebron127_R | tgcaggatccgactcgagtgtcgacTCACAGCAACCCAAGCTCG |
| oLebron128_F | atgcggaggaatcacttccatATGTCCGACATCCAGGC |
| oLebron128_R | tgcaggatccgactcgagtgtcgacTCAGGCCACCTTGCGC |
| oLebron129_F | atgcggaggaatcacttccatATGGTGAGAACAAAGATATACCCCG |
| oLebron129_R | tgcaggatccgactcgagtgtcgacTCAAGCGCAGGCCCCG |
| oLebron130_F | atgcggaggaatcacttccatATGAGCACCGCTATCCGC |
| oLebron130_R | tgcaggatccgactcgagtgtcgacTCAGACGGTTTCCAAAACG |
| oLebron131_F | atgcggaggaatcacttccatATGAGCACCACCCACACC |
| oLebron131_R | tgcaggatccgactcgagtgtcgacTCAGCGTGCGTACTGC |
| oLebron132_F | atgcggaggaatcacttccatATGGGTGTGTGTTTGCTGG |
| oLebron132_R | tgcaggatccgactcgagtgtcgacTCAAGCCAGTGCGCCTAAAG |
| pExTra_seqF | GTACCCGTGTGTACGACCAGC |
| pExTra_uniR | CCCTTCGAGACCATAGATCTGTTCC |
| oLebron4i_F | ggctggtactgaggattcg |
| oLebron5i_F | ggcgaggtcgagtatctgg |
| oLebron16ia_F | aagaggctatcgctgagg |
| oLebron16ib_F | acaatgacggtcgtcttg |
| oLebron16ic_F | agctccgttcggagattgatg |
| oLebron16id_R | ccggcattcaccgcatcc |
| oLebron16ie_R | gtgaccagcaatcgactgg |
| oLebron18i_F | ggattggcagtgtttcg |
| oLebron20ia_F | ttggtgtcggggccgaacg |
| oLebron20ib_F | agtcggcgatcacgttggc |
